# The Common Fund Data Ecosystem (CFDE)

**DOI:** 10.64898/2026.04.10.717672

**Authors:** Julie A. Jurgens, Andreas Bueckle, Jeet Vora, Mano R. Maurya, Taha Mohseni Ahooyi, Erika Zheng, Benjamin Stear, Ding Wang, Caitlin Ree, Srinivasan Ramachandran, Anton Nekrutenko, MacKenzie Brandes, Swathi Thaker, Daniel H. Katz, Monica C. Munoz-Torres, Ido Diamant, Hye-Jung E. Chun, J. Alan Simmons, Sarah K. Tasian, Sherry L. Jenkins, John Erol Evangelista, Hardik Dodia, Surya Saha, Martin A. Lindquist, Vennela Gajjala, Christopher Nemarich, Jimmy Zhen, Karen E. Ross, Anna I. Byrd, Alex Shilin, Vincent T. Metzger, Cristian G. Bologa, Sumana Srinivasan, Dongkeun Jang, Praveen Kumar, Lily D. Taub, Mia P. Levanto, Varduhi Petrosyan, Manju Anandakrishnan, Mariia Kim, Daniel J. B. Clarke, Adriana Ivich, Daniel J. Crichton, Shava Smallen, Dominic Bordelon, Chuming Chen, Andrew J. Schroeder, Ashish Mahabal, Ivan Cao-Berg, Sean Kim, Daniall Masood, Keyang Yu, Kyle J. Gaulton, David Jimenez-Morales, John Michael Rincon, Brendan J. Honick, Wei Wang, Cathy H. Wu, Aleksandar Milosavljevic, Philip D. Blood, Jyl Boline, Tudor I. Oprea, Christophe G. Lambert, Bernard de Bono, Peter J. Park, Jonathan C. Silverstein, Jason Flannick, Jeremy J. Yang, Jeffrey S. Grethe, Shankar Subramaniam, Michael Tiemeyer, Timothy Clark, Matthew T. Wheeler, Ari Kahn, Jennifer Burnette, Rene Ranzinger, Michael C. Schatz, LaFrancis Gibson, Noël P. Burtt, James P. Carson, Jake Y. Chen, Peipei Ping, Sean Davis, Deanne M. Taylor, Katy Börner, Allissa Dillman, Kelli Bursey, Avi Ma’ayan, The CFDE Consortium, Raja Mazumder, Matthew E. Roth, Casey S. Greene

## Abstract

The NIH Common Fund Data Ecosystem (CFDE) integrates data resources from 18 NIH Common Fund programs for discovery and integrative analysis. These programs generate valuable but heterogeneous datasets that can be difficult to discover, access, and reuse. CFDE aims to provide a collaborative, community-built infrastructure that links and enriches Common Fund programs. We describe the evolution, structure, and core technologies of CFDE, including practical approaches that support submission, integration, visualization, and public release of multimodal data. Training programs and workforce initiatives lower barriers to adoption. CFDE has devised solutions to critical issues facing cross-program initiatives, including data scale and heterogeneity, dataset integration, and long-term sustainability. We demonstrate the utility of linking Common Fund resources through integrative tools and cross-dataset queries to yield insights that would otherwise be infeasible. Collectively, CFDE shows that a standards-driven, federated approach enhances and unifies cross-disciplinary resources, fostering collaboration and data-driven discovery.

## Introduction

The NIH Common Fund^1,2^ supports ground-breaking, high-risk research programs that span multiple NIH institutes and that have the potential to accelerate discoveries across biomedical science. While these programs produce high-value datasets across a wide variety of projects and research consortia, their heterogeneity in formats, metadata, and access pathways has impeded data integration and reuse, a long-standing problem in the life sciences and other fields.^3–6^ The Common Fund Data Ecosystem (CFDE) is a collaborative, NIH Common Fund-supported initiative designed to enable community-based standardization, integration, and enhanced accessibility of Common Fund data, thereby facilitating research and discovery (see companion paper; Kano et al.).^7,8^ As of March 2026, CFDE offered access to 10,239,060 files, 2,101,705 biosamples, and 1,256,461 knowledge graph assertions from across 16 CFDE programs (https://data.cfde.cloud/processed). CFDE seeks to address interoperability challenges by connecting communities centered on partially federated yet distinct data platforms. CFDE programs retain autonomy while contributing harmonized metadata to shared discovery platforms for use by the research community. Here, we provide an overview of CFDE and summarize challenges associated with data integration and reuse that necessitated the development of this initiative. We also outline the approach that CFDE has taken to address these challenges and provide examples from the effort.

The Common Fund’s program portfolio encompasses multiple initiatives spanning common and rare disease phenotypes, clinical and basic data modalities, multiple species, and a multitude of biological processes. Data types include genetics and genomics, transcriptomics, epigenomics, proteomics, glycomics, metabolomics, drugs, imaging, and phenotyping, among others (Figure 1; interactive diagram available at https://cfde-sankey.github.io/). Collectively, Common Fund programs contribute to fundamental understanding of biological processes, clinical and translational science, generation of flagship reference maps and datasets, and development of standards for emerging areas such as artificial intelligence (Figure 2, Tables 1-2). Programs cover expansive ground in areas such as glycosylation, extracellular RNAs, the microbiome, molecular responses to exercise, rare disease genetics, bioelectronic medicine, signatures from massive perturbation studies, and illuminating understudied drug targets. Programs have independently developed their own Data Coordinating Centers (DCCs) as well as their own data models and portals, with the work of many spanning more than a decade. Since CFDE funding for each CFDE DCC is limited to 10 years, sustainability remains an ongoing challenge similar to that experienced by other federally funded efforts, despite various innovative approaches.^9–12^

**Figure 1.**
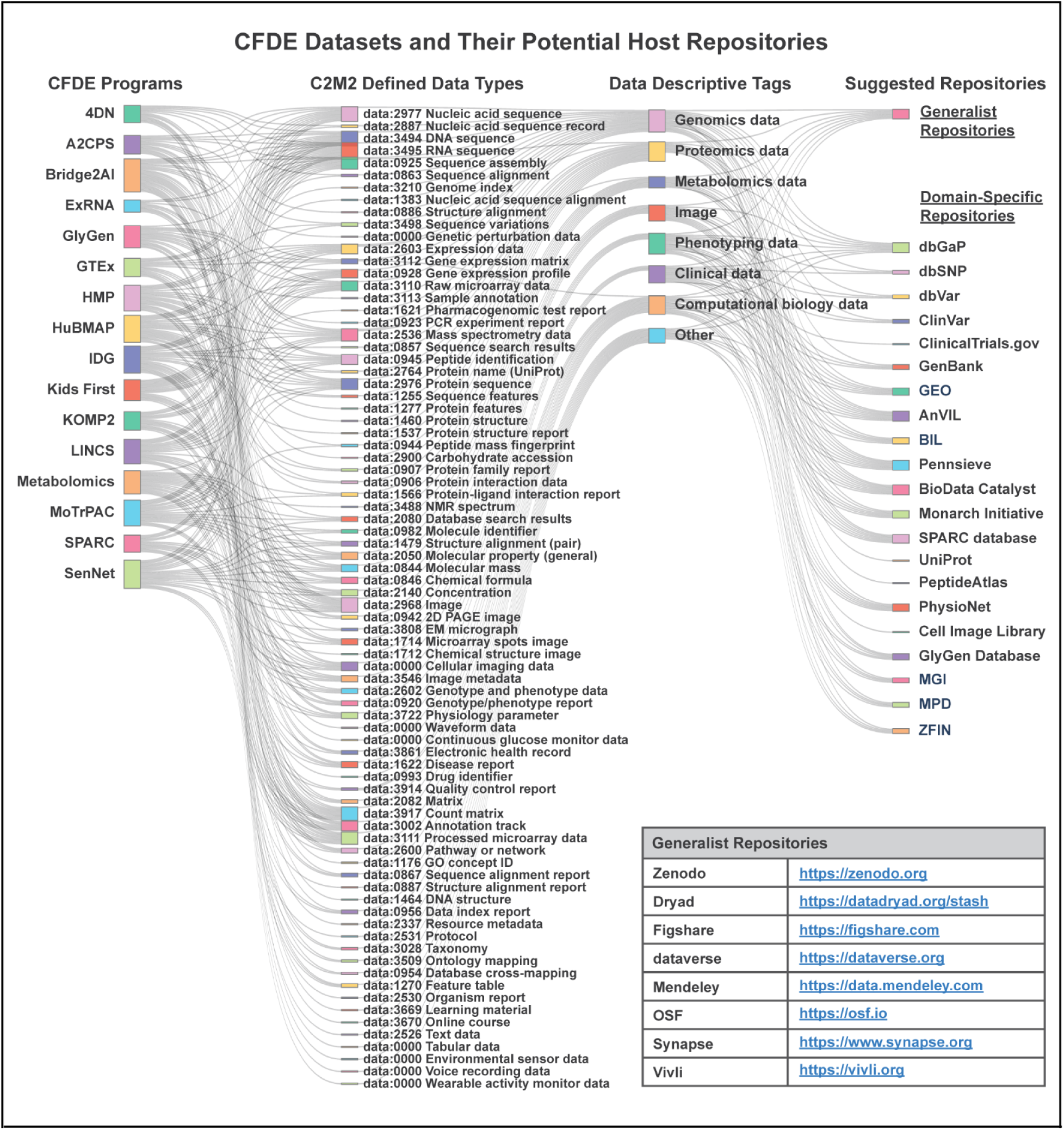
Summary of CFDE digital assets. This Sankey diagram visualizes CFDE digital assets, which currently include datasets generated by 16 CFDE-participating programs (**Column 1**). These multimodal datasets were annotated with EDAM (EMBRACE Data and Methods Ontology)-supported data type terms based on each DCC’s data classifications in their most recent C2M2 submissions (November 2025) and on their data portals. More than 80 data types (**Column 2**) were identified and further organized into eight higher-level categories (**Column 3**) using descriptors commonly adopted by the scientific community to support future data discovery and indexing. The diagram also incorporates 8 Generalist Repositories (**Table**, lower right) and 21 Domain-Specific Repositories (**Column 4**) as potential hosts for CFDE digital assets. An interactive version of the diagram is available at https://cfde-sankey.github.io/. This visualization represents a major CFDE ICC-SC effort supporting three core milestones: (1) assessing Common Fund program needs, (2) collaborating with DRCs to enhance the FAIRness of Common Fund digital assets, and (3) promoting outreach, reuse, and long-term sustainability of CFDE resources. Abbreviations: A2CPS-Acute to Chronic Pain Signatures, AnVIL-Analysis, Visualization, and Informatics Lab-space, BIL-Biomedical Imaging Library, BioData Catalyst-NHLBI BioData Catalyst Data Ecosystem, C2M2-Cross-Cut Metadata Model, CF-Common Fund, CFDE-Common Fund Data Ecosystem, ClinVar-Clinical Variation Database, dbGaP-Database of Genotypes and Phenotypes, dbSNP-Database of Single Nucleotide Polymorphisms, dbVar-Database of Genomic Structural Variation, DCC-Data Coordinating Center, EDAM-The EMBRACE Data and Methods (EDAM) Ontology, exRNA-The exRNA Research Portal, GEO-Gene Expression Omnibus, GTEx-The Genotype-Tissue Expression Project, HMP-The Human Microbiome Project, HuBMAP-The Human BioMolecular Atlas Program, IDG-Illuminating the Druggable Genome, Kids First-Gabriella Miller Kids First Pediatric Research Program, KOMP2-The Knockout Mouse Phenotyping Program, LINCS-Library of Integrated Network-based Cellular Signatures, Metabolomics-Metabolomics Workbench, MGI-Mouse Genome Informatics, Monarch Initiative-Monarch Initiative Integrative Biology Platform, MoTrPAC-The Molecular Transducers of Physical Activity Consortium, MPD-Mouse Phenome Database, OSF-Open Science Framework, Pennsieve-Pennsieve Data Platform, PeptideAtlas-Peptide Atlas Proteomics Resource, PhysioNet-Physiological Signal Archive Network, SenNet-Cellular Senescence Network, SPARC-Stimulating Peripheral Activity to Relieve Conditions, UniProt-Universal Protein Resource, ZFIN-Zebrafish Information Network, 4DN-The 4D Nucleome Program.

**Figure 2.**
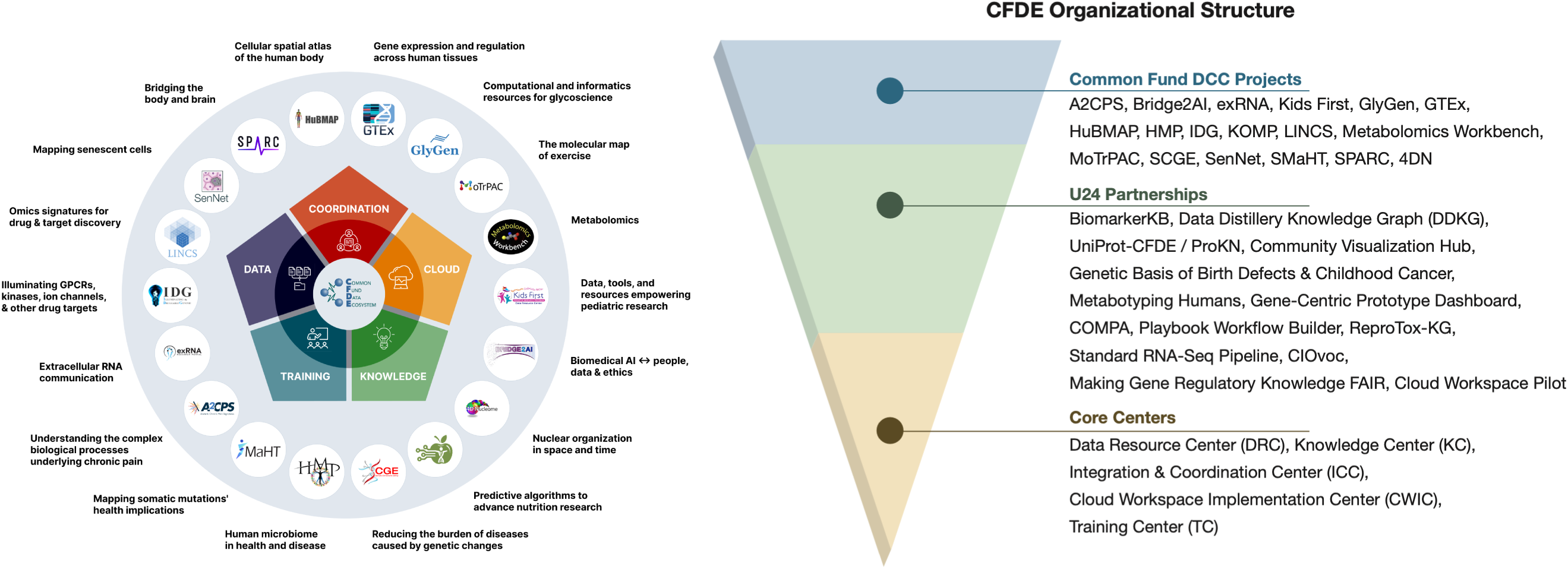
CFDE organization and scientific focus of individual programs. The CFDE consists of eighteen DCCs (circles; top layer of inverted pyramid) that are supported by five centers (spokes surrounding central hub; bottom layer of inverted pyramid). DCCs collaborate with one another and with non-CFDE entities to form partnerships (middle layer of inverted pyramid). Taglines summarize the major focus areas of each DCC. Abbreviations: A2CPS-Acute to Chronic Pain Signatures, B2AI-The Bridge2AI Consortium, CFDE-Common Fund Data Ecosystem, CWIC-Cloud Workspace Implementation Center, DCC-Data Coordinating Center, DRC-Data Resource Center, exRNA-The extracellular RNA Communication Program, GTEx-The Genotype-Tissue Expression Project, HMP-The Human Microbiome Project, HuBMAP-The Human BioMolecular Atlas Program, ICC-Integration and Coordination Center, IDG-Illuminating the Druggable Genome, KC-Knowledge Center, Kids First-Gabriella Miller Kids First Pediatric Research Program, KOMP-The Knockout Mouse Project, LINCS-Library of Integrated Network-based Cellular Signatures, MoTrPAC-The Molecular Transducers of Physical Activity Consortium, MW-Metabolomics Workbench, SCGE-Somatic Cell Genome Editing, SenNet-Cellular Senescence Network, SMaHT-The Somatic Mosaicism across Human Tissues Network, SPARC-Stimulating Peripheral Activity to Relieve Conditions, TC-Training Center, 4DN-The 4D Nucleome Program.

**Table 1.**
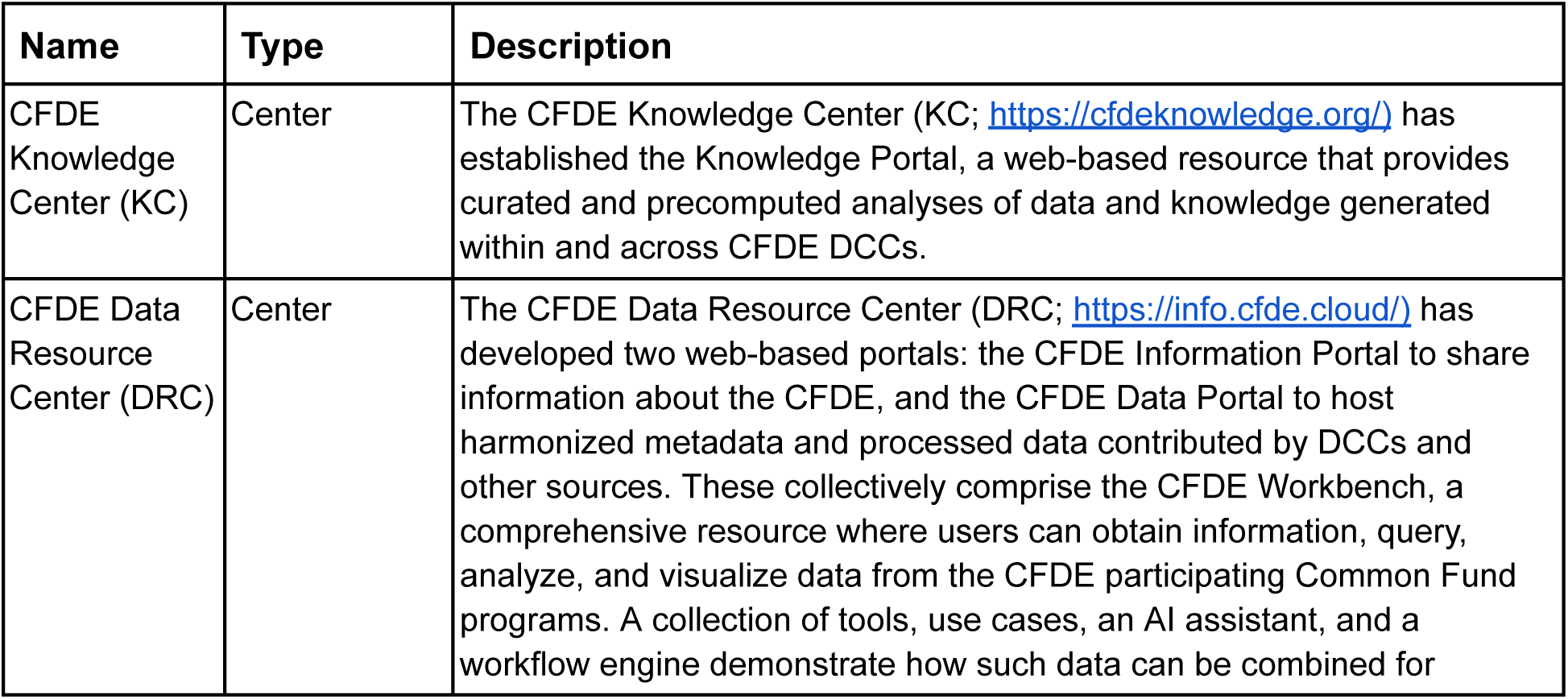
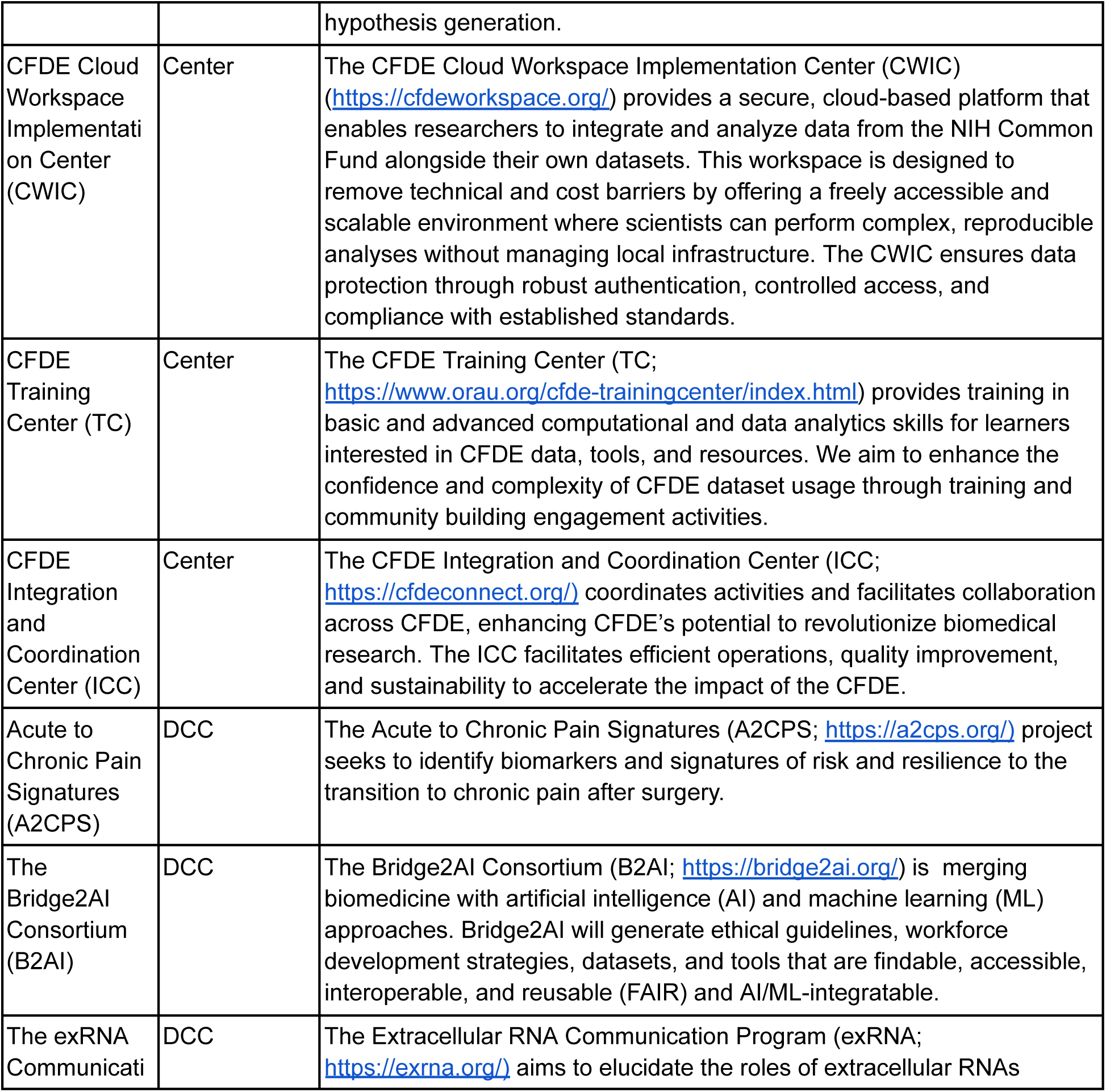
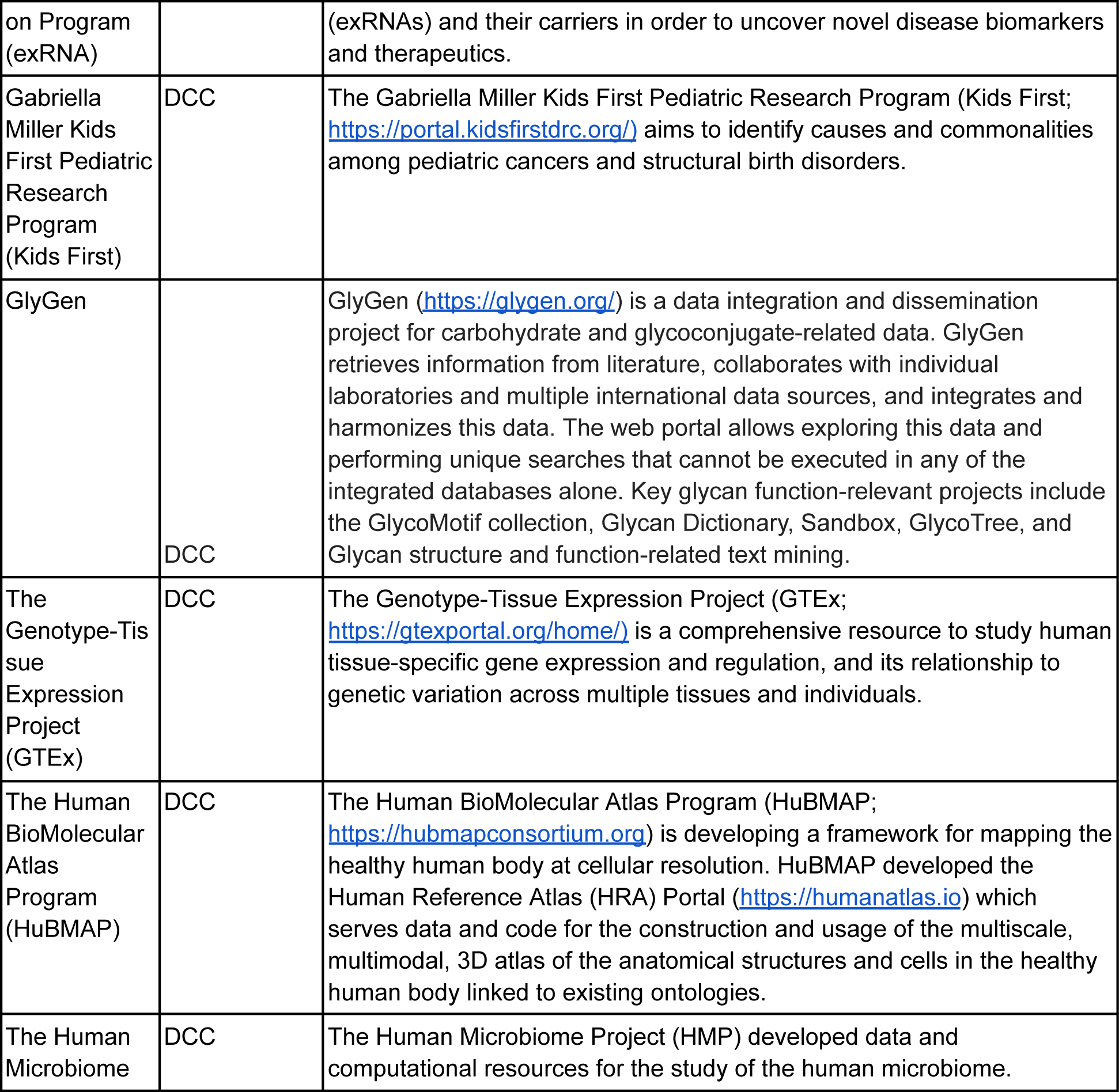
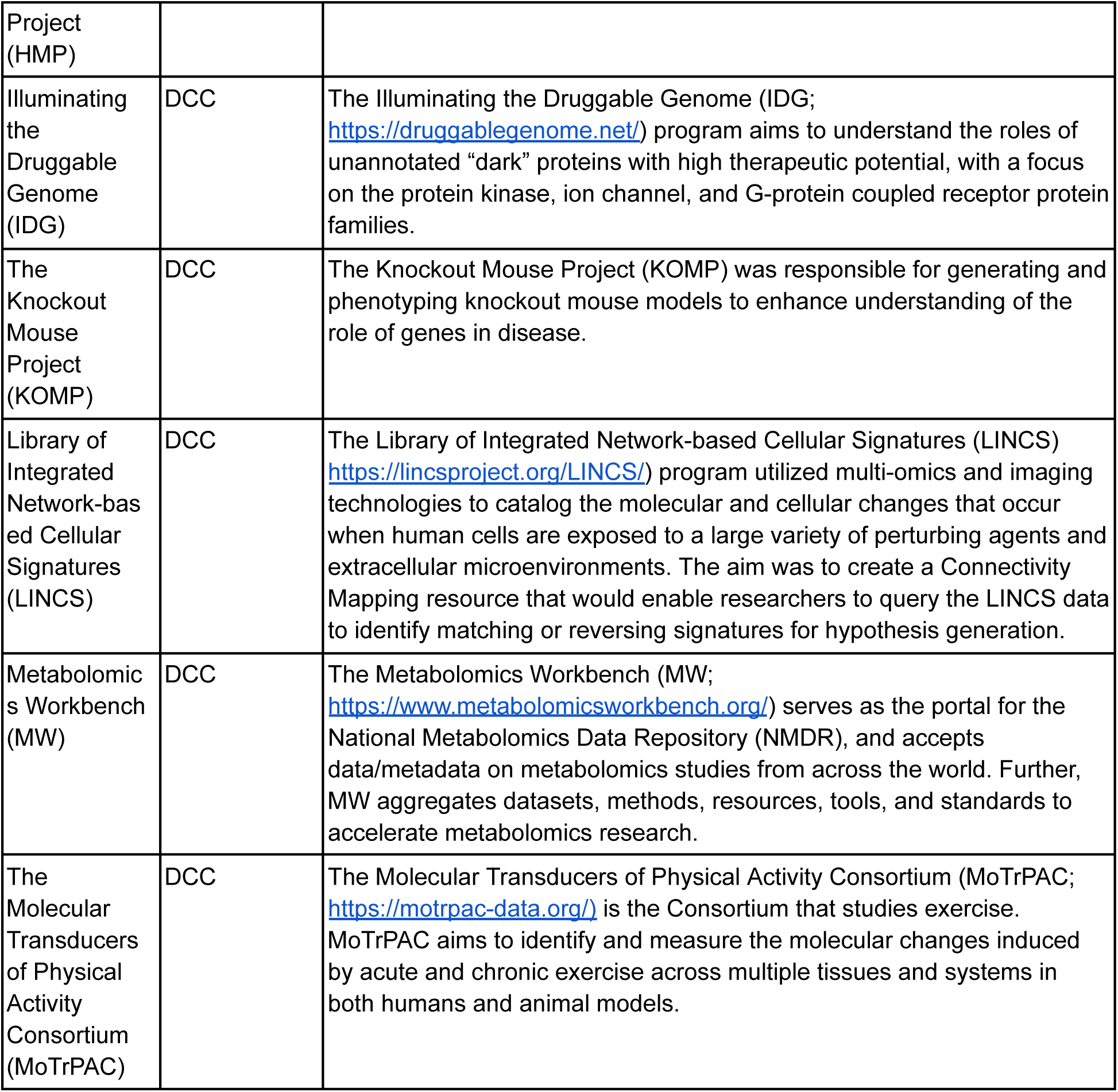
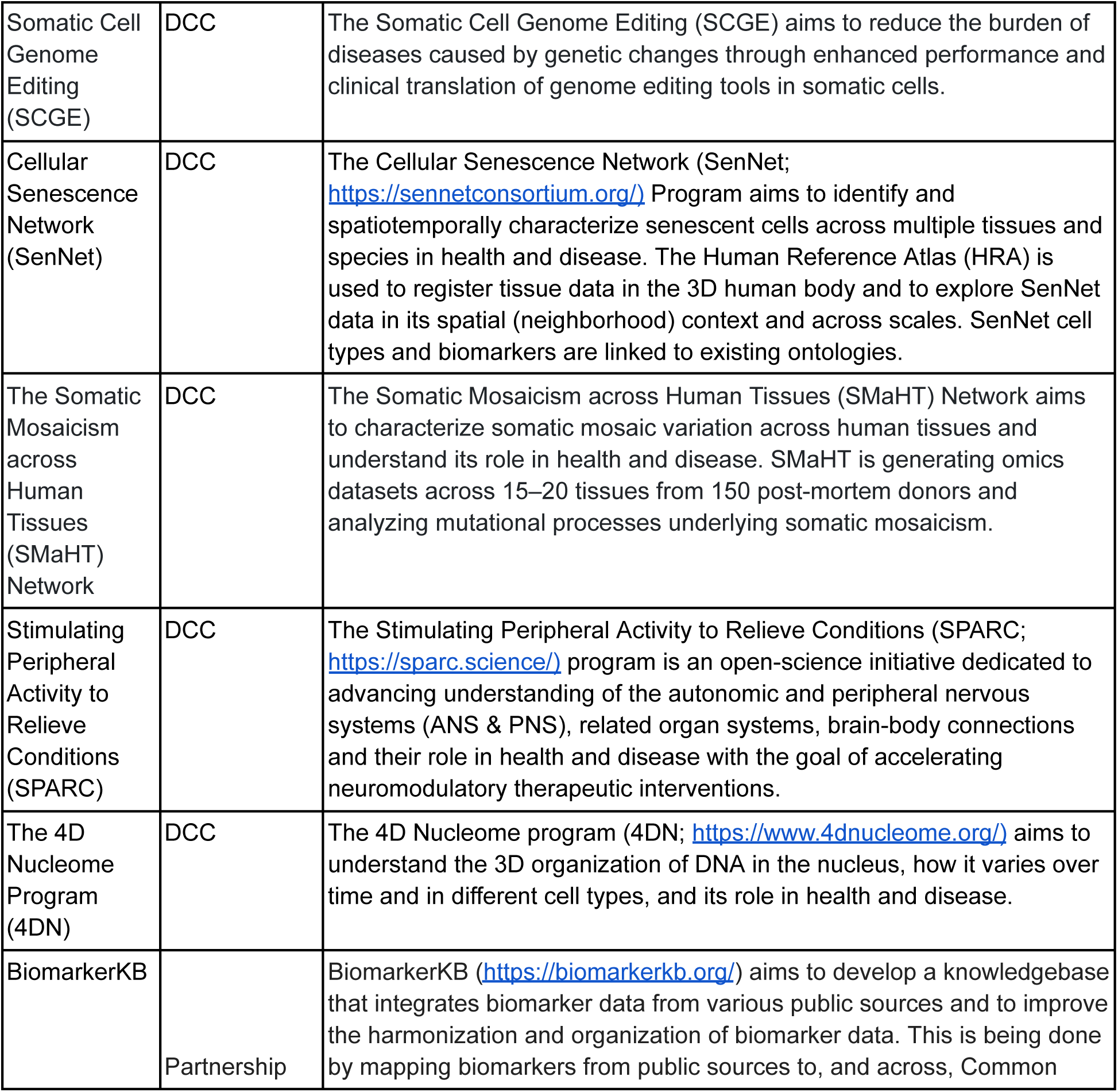
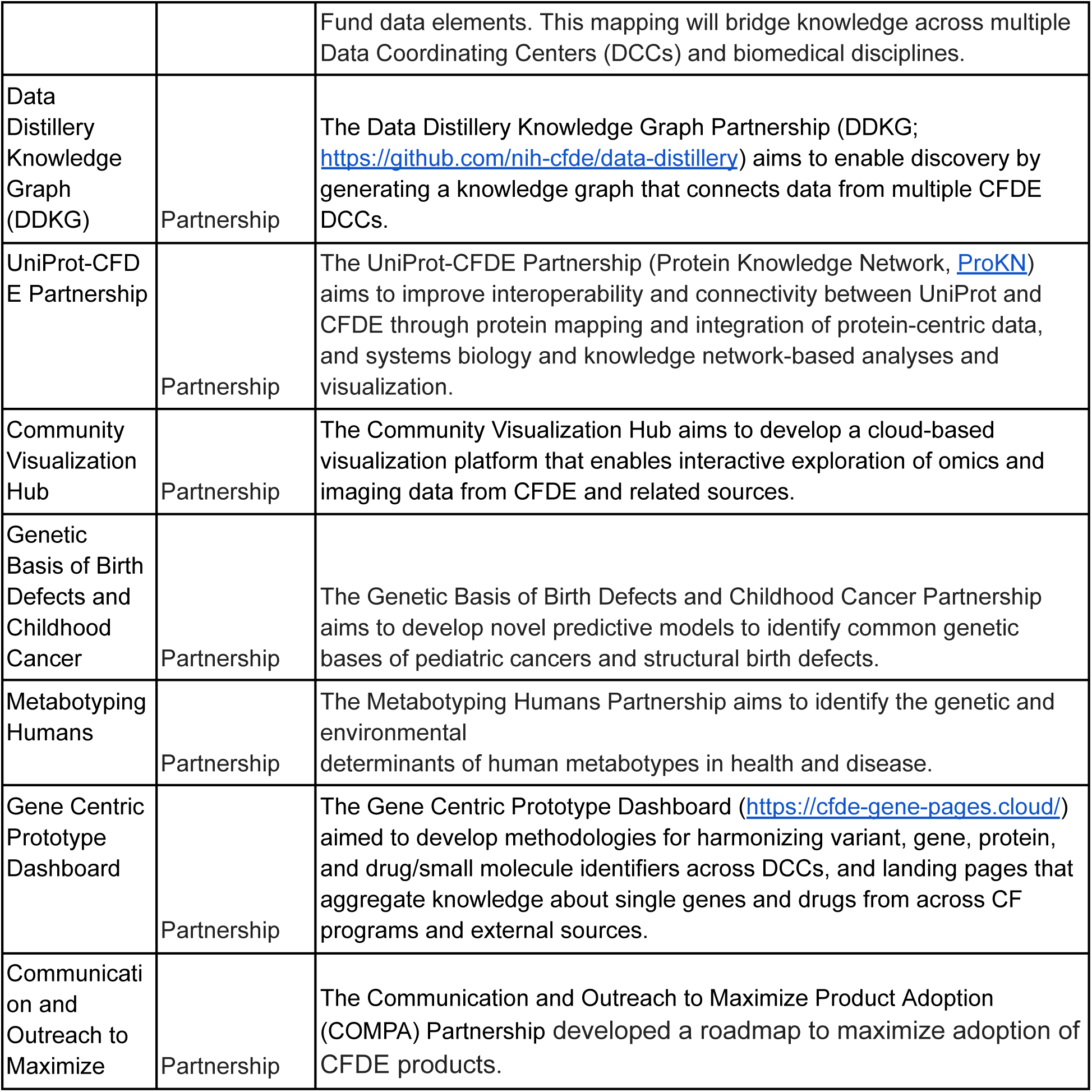
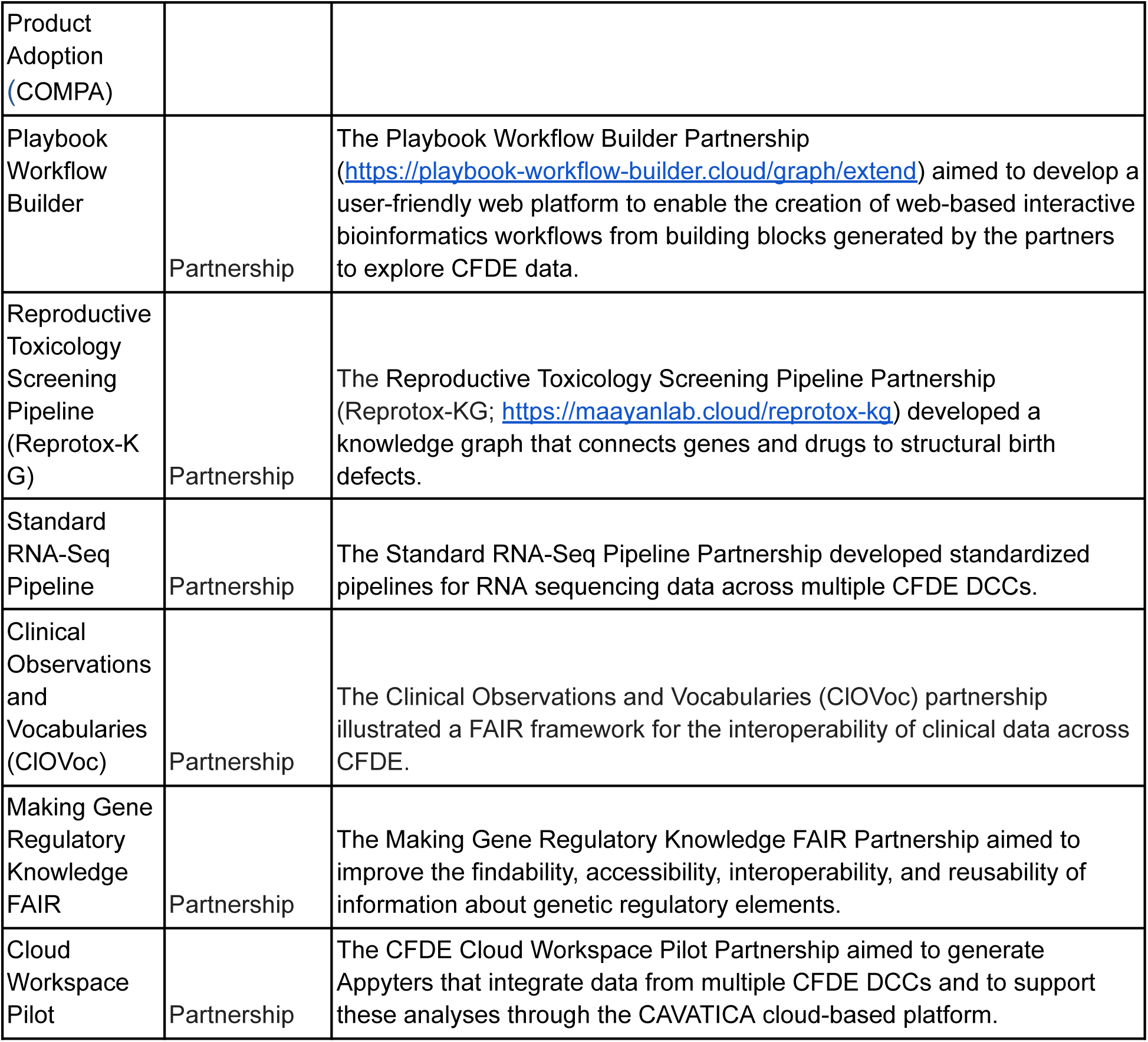
Overview of CFDE Centers, Data Coordinating Centers (DCCs), and Partnerships. Names, descriptions, and weblinks are provided for CFDE Centers, DCCs, and Partnerships. Abbreviations: A2CPS-Acute to Chronic Pain Signatures, B2AI-The Bridge2AI Consortium, CFDE-Common Fund Data Ecosystem, CWIC-Cloud Workspace Implementation Center, DCC-Data Coordinating Center, DRC-Data Resource Center, exRNA-The extracellular RNA Communication Program, GTEx-The Genotype-Tissue Expression Project, HMP-The Human Microbiome Project, HuBMAP-The Human BioMolecular Atlas Program, ICC-Integration and Coordination Center, IDG-Illuminating the Druggable Genome, KC-Knowledge Center, Kids First-Gabriella Miller Kids First Pediatric Research Program, KOMP-The Knockout Mouse Project, LINCS-Library of Integrated Network-based Cellular Signatures, MoTrPAC-The Molecular Transducers of Physical Activity Consortium, MW-Metabolomics Workbench, SCGE-Somatic Cell Genome Editing, SenNet-Cellular Senescence Network, SMaHT-The Somatic Mosaicism across Human Tissues Network, SPARC-Stimulating Peripheral Activity to Relieve Conditions, TC-Training Center, 4DN-The 4D Nucleome Program

**Table 2.**
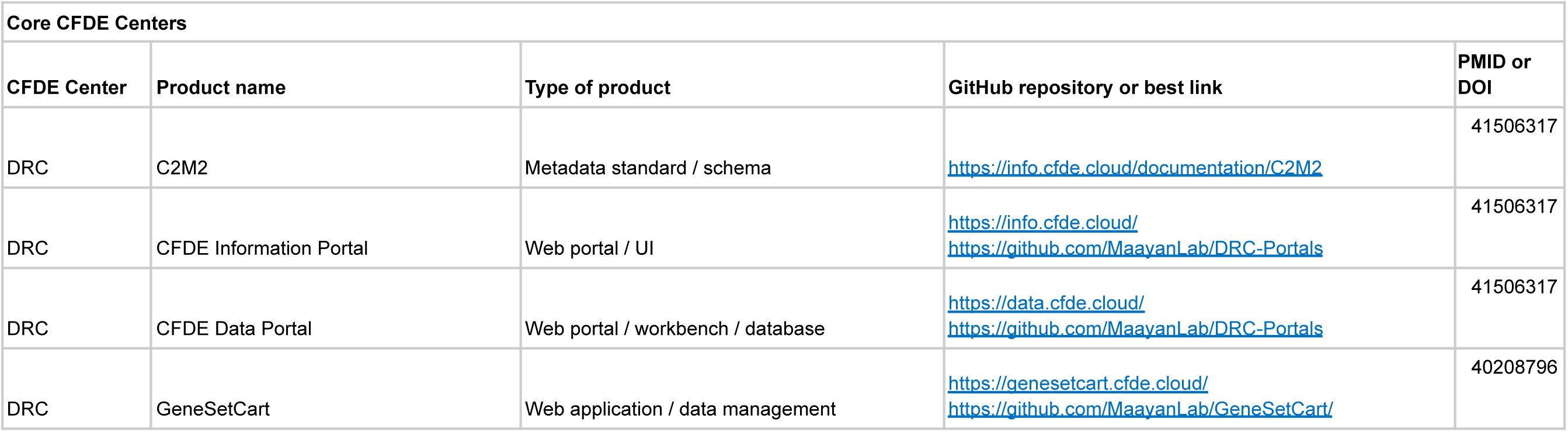
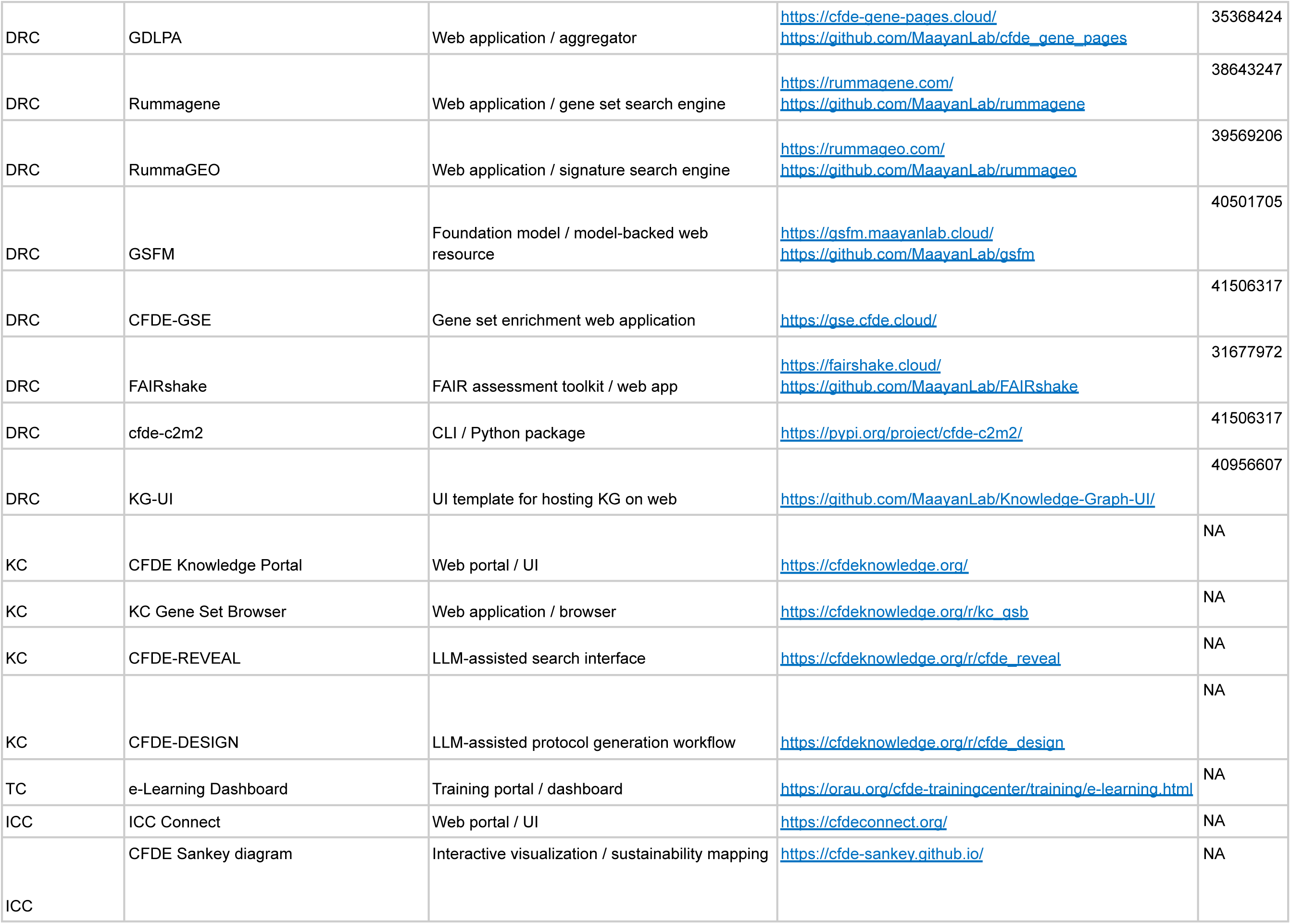
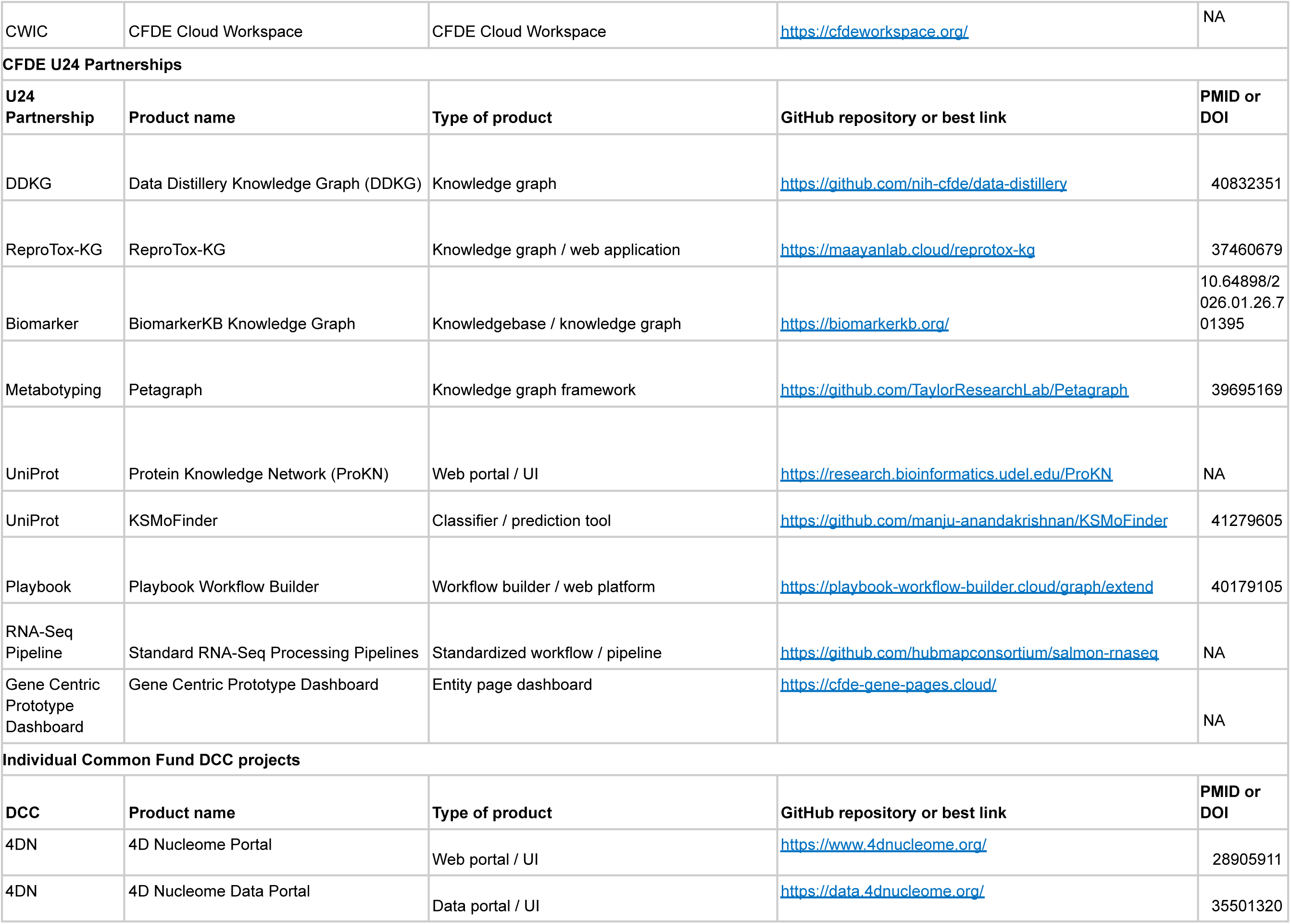
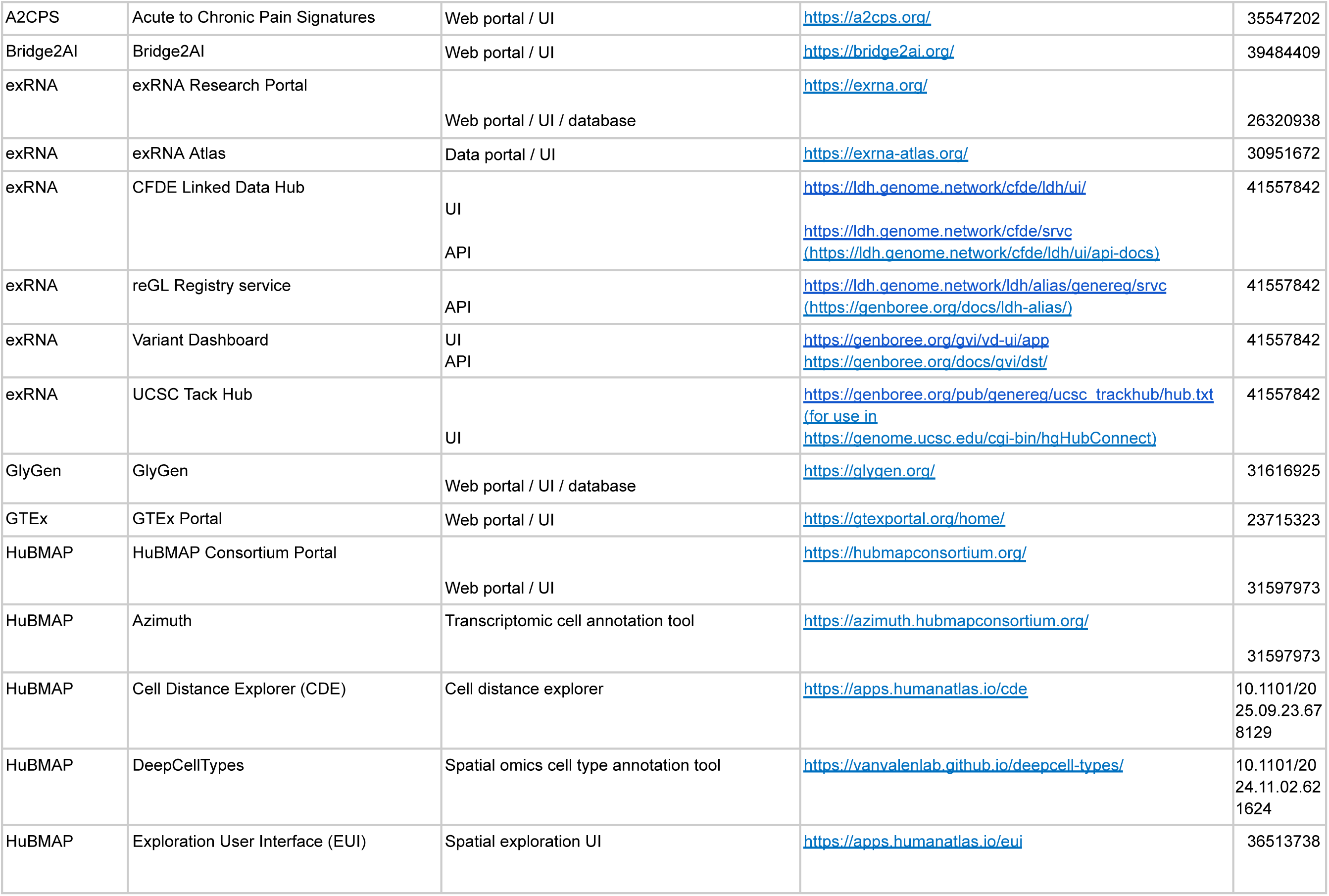
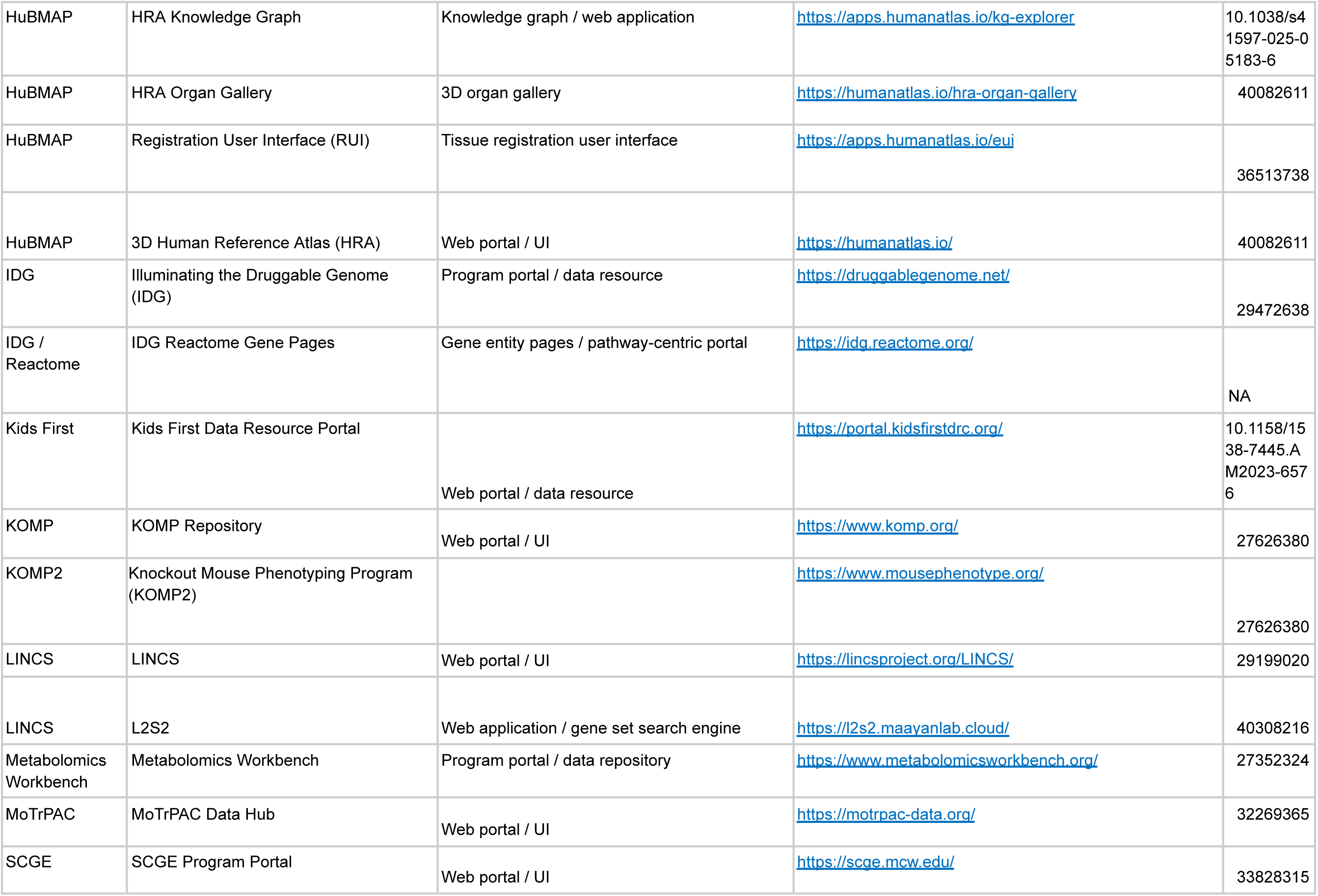
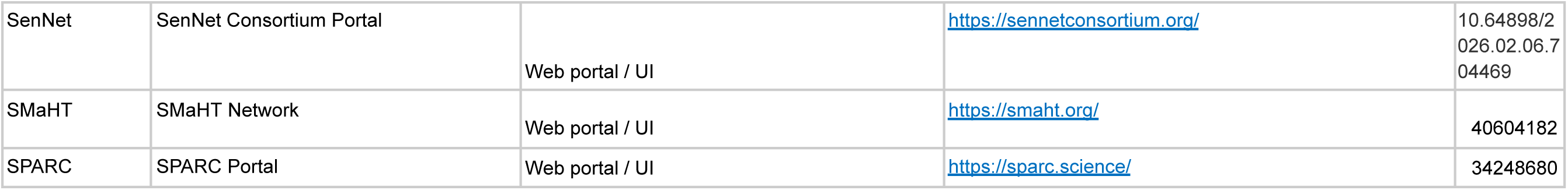
List of described CFDE products. Products described in the manuscript, listed alongside links to their websites and/or GitHub repositories and PMIDs or DOIs. Products are grouped based on their origination from CFDE Centers, U24 Partnership Projects, or DCCs. Abbreviations: A2CPS-Acute to Chronic Pain Signatures, C2M2-Cross-Cut Metadata Model, CDE-Cell Distance Explorer, CFDE-Common Fund Data Ecosystem, CFDE-GSE-Common Fund Data Ecosystem Gene Set Enrichment, CLI-Command Line Interface, CWIC-Cloud Workspace Implementation Center, DCC-Data Coordinating Center, DDKG-Data Distillery Knowledge Graph, DRC-Data Resource Center, DOI-digital object identifier, EUI-Exploration User Interface, exRNA-The exRNA Research Portal, GDLPA-Gene and Drug Landing Page Aggregator, GSFM-Gene Set Foundation Model, GTEx-The Genotype-Tissue Expression Project, HuBMAP-The Human BioMolecular Atlas Program, HRA-Human Reference Atlas, ICC-Integration and Coordination Center, IDG-Illuminating the Druggable Genome, KC-Knowledge Center, KG-Knowledge Graph, Kids First-Gabriella Miller Kids First Pediatric Research Program, KOMP-The Knockout Mouse Project, L2S2-LINCS L1000 Signatures Search, LINCS-Library of Integrated Network-based Cellular Signatures, MoTrPAC-The Molecular Transducers of Physical Activity Consortium, MW-Metabolomics Workbench, PMID-PubMed identifier, RUI-Registration User Interface, SCGE-Somatic Cell Genome Editing, SenNet-Cellular Senescence Network, SMaHT-The Somatic Mosaicism across Human Tissues Network, SPARC-Stimulating Peripheral Activity to Relieve Conditions, TC-Training Center, UI-user interface, 4DN-The 4D Nucleome Program.

Initially, CFDE programs were more inwardly focused with less emphasis on cross-program integration and interoperability. As CFDE matured and more programs were funded, CFDE sought to facilitate cross-disciplinary and collaborative data and code reuse across programs and projects. To enable this, CFDE developed the Cross-Cut Metadata Model (C2M2), a metadata standard that supports unified search across datasets.^7^ CFDE has also adopted community standards for representing highly processed data such as knowledge graphs^13,14^, gene regulatory information,^15^ gene matrix transpose (GMT) files, and Croissant files,^16^ with the goal of working toward FAIR (Findable, Accessible, Interoperable, and Reusable) principles.^17^

CFDE has undergone two phases in its development. During Phase I (2020-2022), CFDE was composed of Data Coordinating Centers (DCCs) from individual Common Fund programs and one CFDE Coordination Center.^18^ The DCCs and CFDE Coordination Center collaborated via partnerships and working groups. Phase I focused on laying the groundwork for an ecosystem through development of community, infrastructure, and a few key products such as initial knowledge graphs, a preliminary portal for query of Common Fund data and information, and an initial cloud workspace. Phase II (2022-2027) aims to rapidly build upon the framework developed in Phase I, with the goal of demonstrating broad use of Common Fund data and tools for discovery. To achieve this goal, the NIH established five Centers to build awareness of CFDE resources and provide sufficient training and infrastructure for their use: the Data Resource Center (DRC), Knowledge Center (KC), Integration and Coordination Center (ICC), Cloud Workspace Implementation Center (CWIC), and Training Center (TC). The current structure of CFDE, comprising these 5 centers and 18 DCCs (Figure 2) along with multiple partnerships, directly enables new avenues of research and provides multiple entry points for users to access the ecosystem. Additionally, the NIH has developed other funding mechanisms (e.g., pilot projects; https://commonfund.nih.gov/dataecosystem/FundedResearch) to further demonstrate Common Fund data utility.

CFDE supports the biomedical research community by fostering cross-program collaborations across Common Fund programs,^13,14,19–21^ illustrating the utility of developing and implementing standards for metadata harmonization, and lowering barriers to begin exploring the ecosystem. CFDE’s approach is designed to promote interoperability and long-term impact of Common Fund resources by meeting research communities where they are with the capabilities to connect to broader resources, ultimately enabling a sustainable ecosystem that enhances and accelerates answering integrative research questions with best-in-class data resources.

## Results

### The CFDE approach to (meta)data

The Common Fund programs have produced data resources that offer a unique opportunity to observe and analyze biomedical processes at a scale and in a way that is often unattainable from other data resources. However, since Common Fund programs were established independently and asynchronously, each program created its own data types, data model, data elements, and ontologies to represent its datasets. To realize the vision of easy access, integration, analysis, visualization, and exploration of data across publicly available Common Fund data resources, CFDE provides a thin layer of metadata harmonization through the C2M2 to enable integration and interoperability of multimodal datasets and exposes datasets ready to be merged for greater impact, as described below. Notably, some CFDE programs also rely on external repositories to house their data since they cannot be made accessible through CFDE. For example, human sequencing data from the Gabriella Miller Kids First Pediatric Research Program^22^ must be maintained through controlled access repositories.

CFDE is not a true runtime federated query system. It creates a centralized, harmonized metadata catalog from periodic C2M2 data package submissions for cross-DCC discovery. The datasets remain distributed and under the autonomy of the participating DCCs. After datasets are discovered via a unified metadata search, they can be retrieved from DCCs using the provided persistent_id/access_url/drs links. Therefore, CFDE is a hybrid federated data ecosystem rather than either a fully centralized repository or a pure runtime federated query system.

### CFDE (meta)data formats & ontologies promote interoperability

A key output of CFDE interoperability initiatives is the Cross-Cut Metadata Model (C2M2; Table 3; Figure 3; https://data.cfde.cloud/img/C2M2_ERD_no_FKlabel_edited.png; https://info.cfde.cloud/documentation/C2M2). C2M2 standardizes metadata on experimental resources and products while promoting the harmonization and search integration of domain-specific data repositories.^7^ Notably, C2M2 is a model of common CFDE metadata elements, not all metadata elements; it brings important shared attributes into the model but does not require all metadata elements across all projects to be aligned. This framework enables unified querying, aggregation, and integration of heterogeneous Common Fund datasets using harmonized metadata. C2M2 was initially designed for existing Common Fund datasets and has been adapted over time for data modalities from newer Common Fund projects, such as chromatin topology data from the 4D Nucleome project^23,24^ and human clinical data from Bridge2AI.^25^ C2M2 organizes metadata into core (Subject, Biosample, File), container (Collection, Project), and administrative (ID_Namespace, DCC) entities, enriched with ontology-driven attributes for taxonomy, anatomy, disease, assay, and data type (Figure 3). Figure 4 shows an example of metadata in the C2M2 format for a single project, PR000663 (Biomarkers of NAFLD progression: a lipidomics approach to an epidemic), from the Metabolomics Workbench (MW) DCC.^26^ Each piece of information in the C2M2 is accompanied by well-established ontologies, dictionaries, and mappings of DCC-provided namespaces to those defined by data providers. The DRC facilitates submission, ingestion and processing of C2M2 metadata to enable user queries for data discovery across the DCCs.^8^

**Figure 3.**
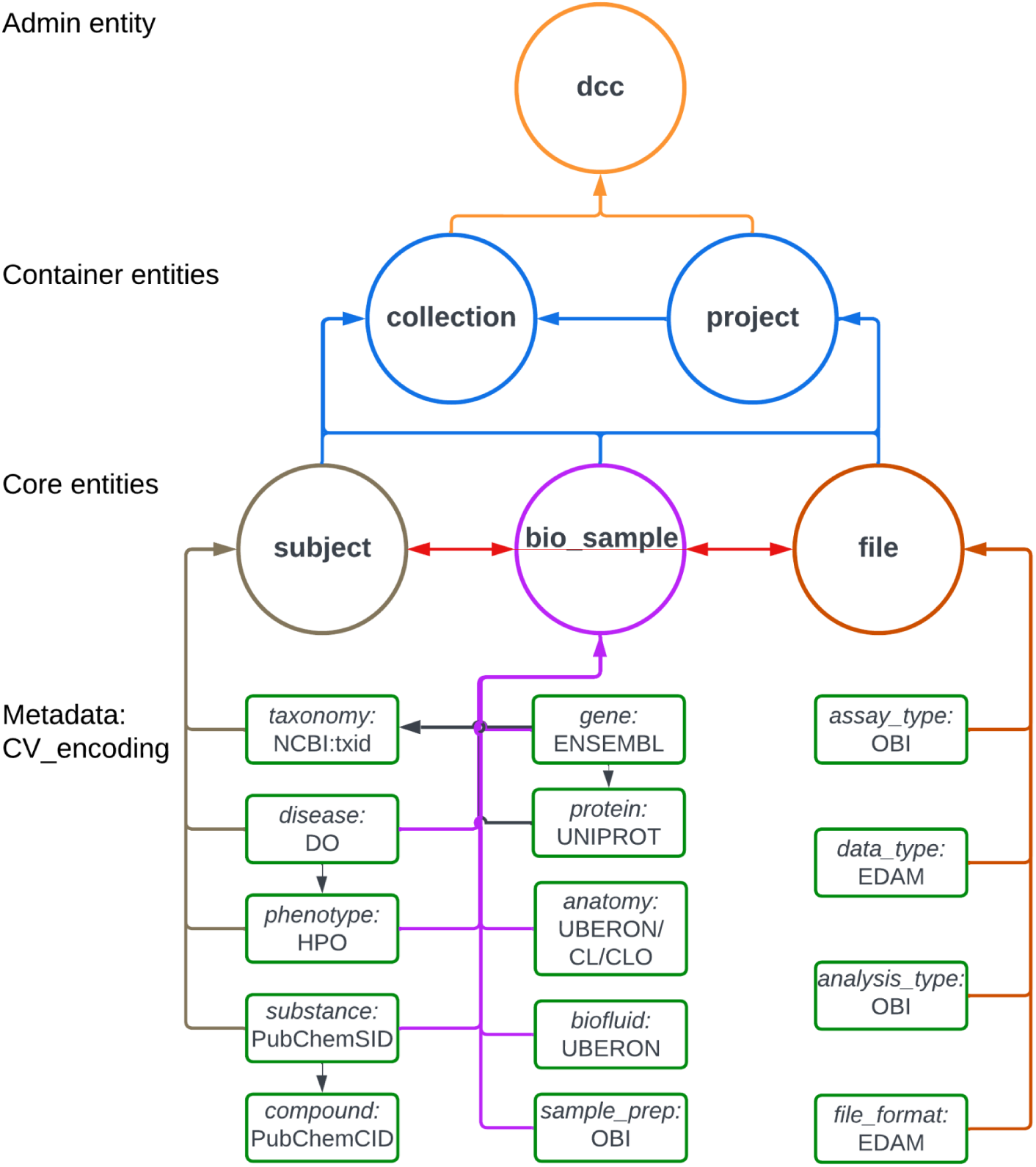
Ontologies/Controlled vocabularies used in the Cross-Cut Metadata Model (C2M2). A simplified schema showing the C2M2 entities and their metadata encoding (green boxes) in corresponding ontologies/controlled vocabularies.

**Figure 4.**
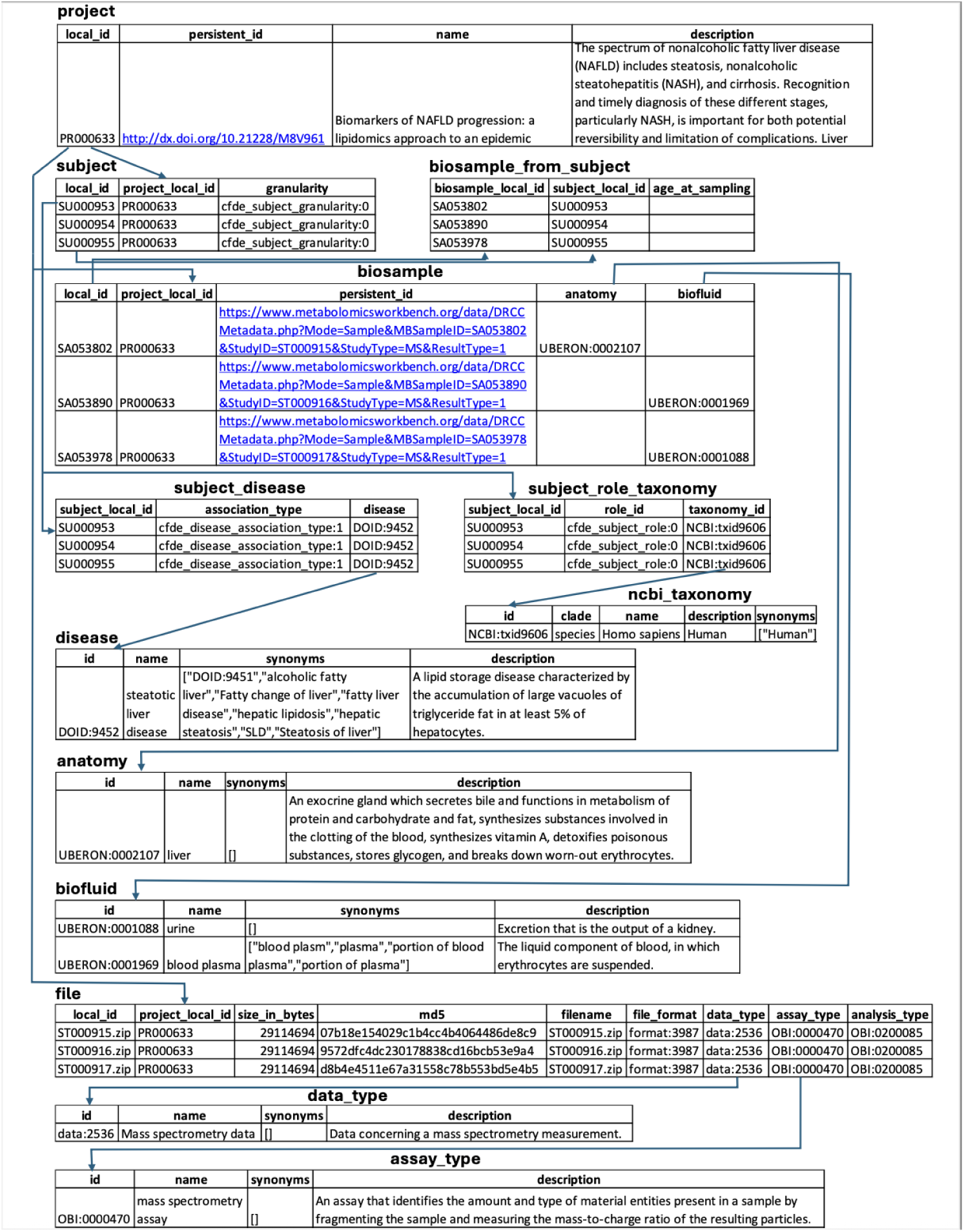
Example of metadata in the C2M2 format. This figure shows an example of metadata in the C2M2 format, including the connections between the tables, for a single project, PR000663 (http://dx.doi.org/10.21228/M8ZX0W), from the Metabolomics Workbench (MW) DCC. It can be noted that this is only partial information, and some empty and id_namespace columns have been omitted. This project is about the study of Nonalcoholic Fatty Liver Disease (NAFLD) in humans (the project table). As per the information at MW, there are three studies under this project, each study with its own lumped subject ID. Thus, there are a total of 3 subject IDs associated with this project (the subject table). There are a total of 264 biosamples collected in the project, of which only three are listed due to space constraints (the biosample table). For the subjects, associated information such as disease, e.g., steatotic liver disease, is captured in the tables subject_disease and disease, and taxonomy for humans is captured in the tables subject_role_taxonomy and ncbi_taxonomy. For the biosamples, anatomy and biofluid information are listed in the respective tables. On the MW website, for this project, three files can be accessed and downloaded, and the relevant C2M2 information is listed in the file table. Details about the relevant data_type and assay_type for these files are captured in the tables data_type and assay_type. Depending upon the type of metadata information from a DCC, there can be additional tables with suitable information, such as those related to collections. A given DCC may have information on several projects and/or collections and the overall information in various tables will be the union of such information from different projects and/or collections.

**Table 3.** List of Ontologies and Controlled Vocabularies used in the Cross-Cut Metadata Model (C2M2) The C2M2 model integrates standard ontologies and controlled vocabularies for a host of metadata fields, defined here. Abbreviations: C2M2-Cross-Cut Metadata Model, CL-Cell Ontology, CLO-Cell Line Ontology, DO-Disease Ontology, EDAM-The EMBRACE Data and Methods (EDAM) Ontology, HPO-Human Phenotype Ontology, MPO-Mammalian Phenotype Ontology, NCBI:txid-National Center for Biotechnology Information Taxonomy Identifier, OBI-The Ontology for Biomedical Investigations, PubChem-Public Chemical Information Resource, UniProtKB-UniProt Knowledgebase

| Metadata | Ontology/Controlled vocabulary |
| --- | --- |
| Analysis | OBI<sup>86</sup> |
| Anatomy | UBERON<sup>30</sup> |
| Assay | OBI<sup>86</sup> |
| Biofluid | UBERON<sup>30</sup> |
| Cell | CL<sup>87</sup> |
| Cell line | CLO<sup>88</sup> |
| Compound | PubChem<sup>89</sup> |
| Data Type | EDAM<sup>90</sup> |
| Disease | DO<sup>91</sup> |
| File Format | EDAM<sup>90</sup> |
| Gene | ENSEMBL<sup>92</sup> |
| Phenotype | HPO<sup>93</sup>/MPO<sup>94</sup> |
| Protein | UniProtKB<sup>39</sup> |
| Substance | PubChem<sup>89</sup> |
| Taxonomy | NCBI:txid<sup>95</sup> |

The Acute to Chronic Pain Signatures (A2CPS)^27^ project illustrates that C2M2 can be valuable as more than a downstream metadata exchange format. The program adopted the core C2M2 model internally because its primary entities and relationships matched much of what A2CPS needed to represent at a high level, including participants, biospecimens, assays, files, and study organization.^28^ Where A2CPS required additional expressivity, particularly for longitudinal study events, perioperative milestones, and repeated clinical observations, the model was extended or adapted to capture those program-specific concepts while preserving compatibility with CFDE-aligned metadata practices. Using a C2M2-informed internal representation in this way both supported local harmonization and substantially simplified later submission to CFDE by reducing the amount of downstream remapping required.

Metadata coverage for C2M2 submissions over time is shown for one CFDE DCC, the Gabriella Miller Kids First Pediatric Research Program (Table 4).^22^ These longitudinal C2M2 submission metrics reflect some variability due to two factors: firstly, variability driven by the standard ongoing data management activities of the Kids First DCC, including continuous maintenance and augmentation of study data; and secondly, variability driven by ongoing DCC C2M2 backend and pipeline improvements. For example, early incremental iterative testing and discovery in solidifying best data sharing practices and workflow improvements in transitioning from manual to automated pipelines led to initial fluctuations in metadata coverage. Data maintenance during Year 3 also led to a decrease in coverage for some entities during that time. However, results overall demonstrate progress in improving metadata coverage for certain aspects of C2M2 submissions over time and continuous improvement in the scope and scale of biosamples, files, projects, and subjects. Growth was driven by a combination of improved data measurement, tracking, and pipelines by the DCC, while data quality improved, in part, resulting from the validation and rigor introduced by the CFDE C2M2 manifest submission process.

**Table 4.**
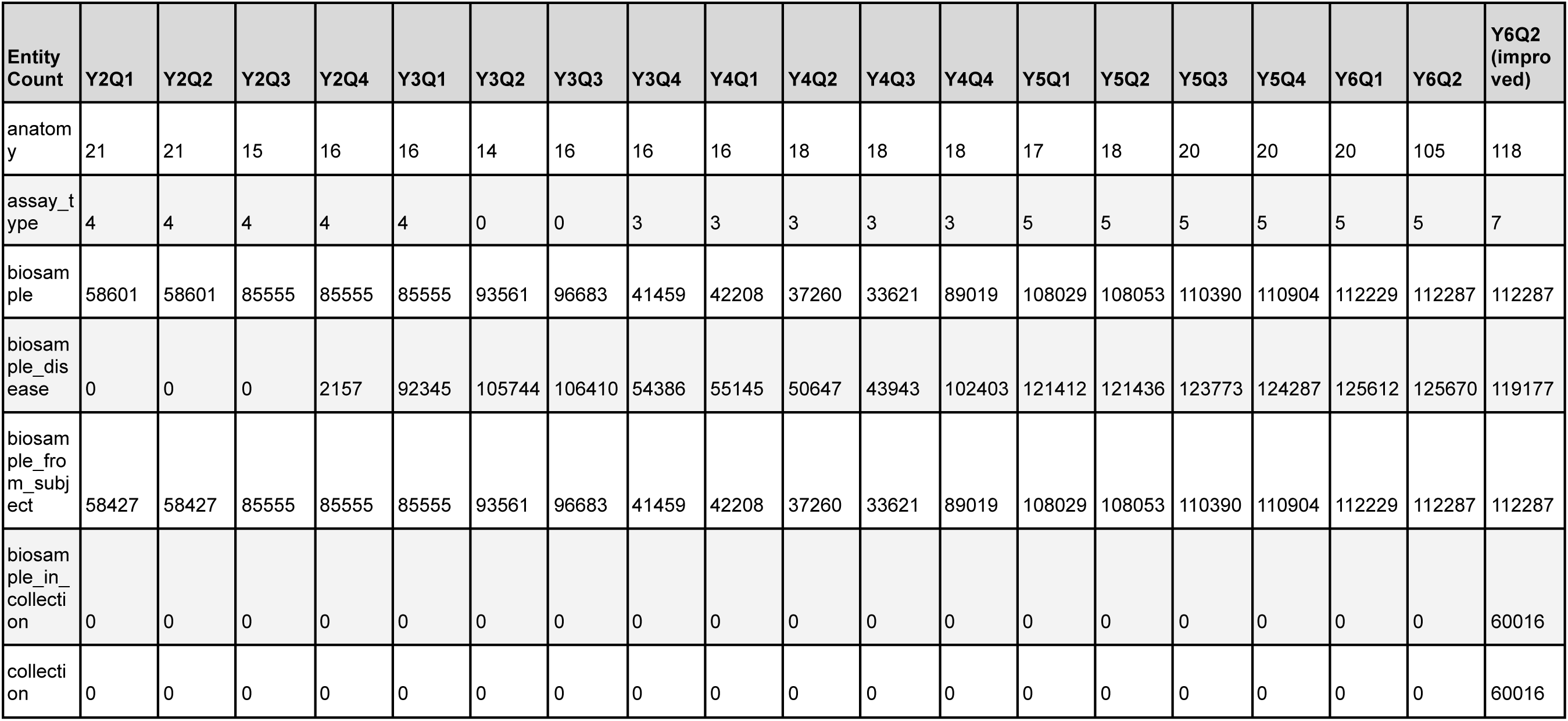
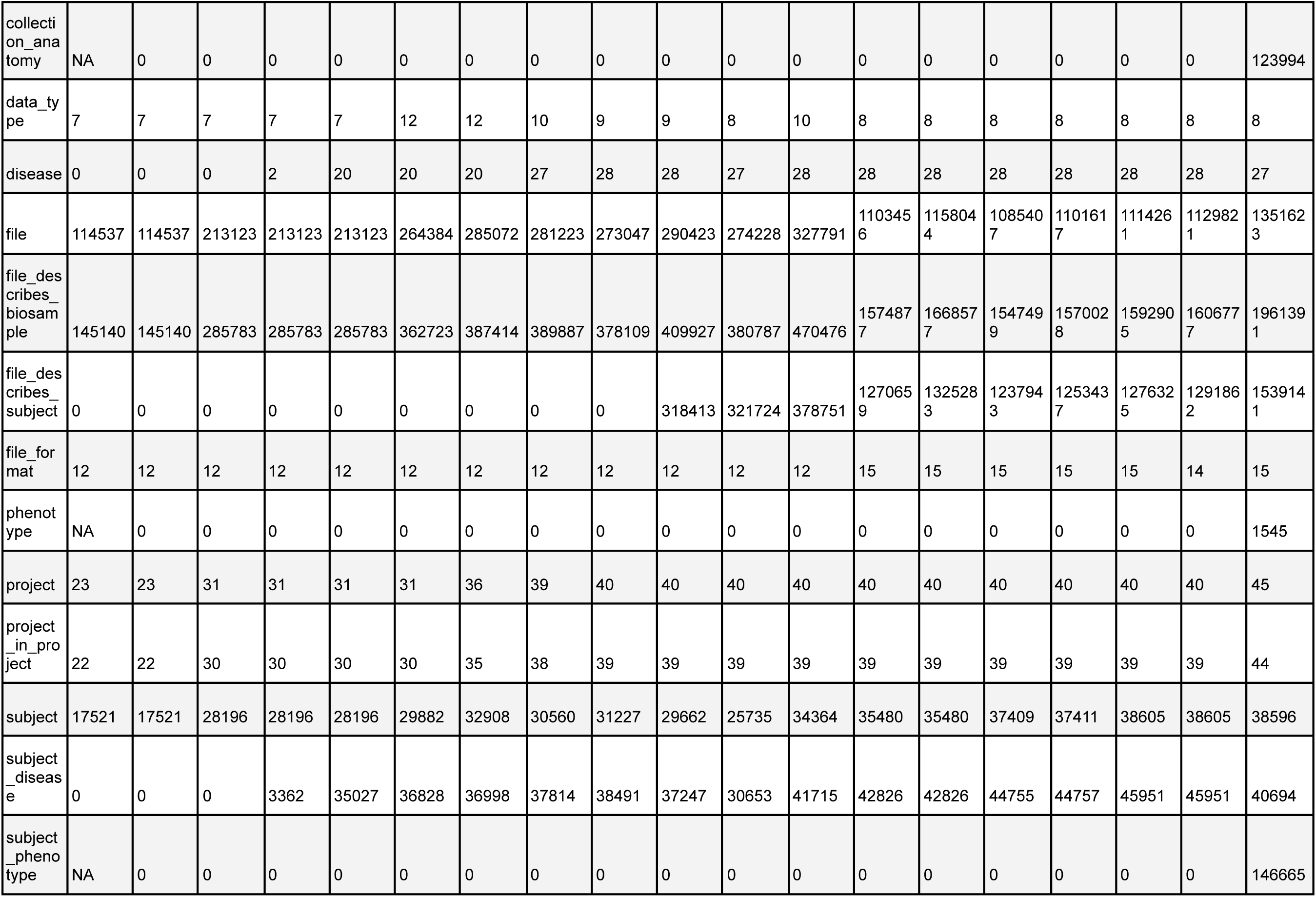
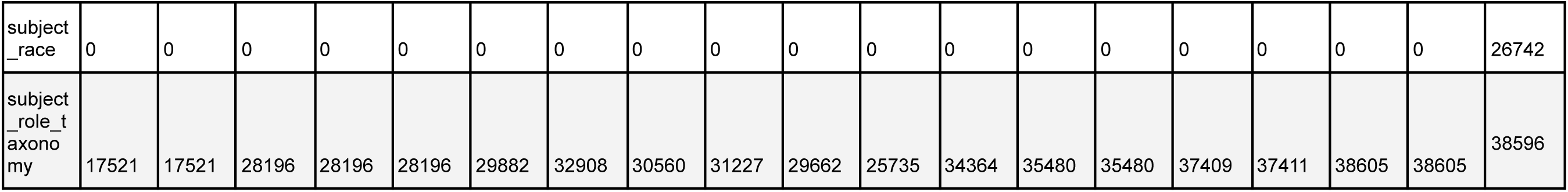
Tracked metadata coverage for C2M2 submissions over time for one CFDE DCC, the Gabriella Miller Kids First Pediatric Research Program. Metadata coverage for CFDE C2M2 submissions over time for the Gabriella Miller Kids First Pediatric Research Program DCC. CFDE C2M2 submissions were made on a quarterly basis for CFDE funding years beginning in January 2022 (Y2Q1) through the present (March 2026; Y6Q2). Counts are provided for each submitted metadata entity type. Some additional improvements have been planned but not yet deployed for the Year 6 Q2 submission; these are projected in the “Q2 Improved” column. Abbreviations: Q-quarter, Y-year.

A primary objective of CFDE was to develop standardized data and metadata formats, ontologies, and evidence attribution to enable interoperability.^7^ A metadata model for experimental data was developed by two working groups approaching different aspects of interoperability. The CFDE Ontology Working Group selected the most appropriate ontology or controlled vocabulary for metadata in C2M2 (Table 3, Figure 3; https://data.cfde.cloud/img/C2M2_ERD_no_FKlabel_edited.png). Ontologies and controlled vocabularies were selected to be open access, stable but not static, community-endorsed, cross-mappable with other standards, and easily adoptable across existing databases. They themselves need to follow FAIR principles.^17^ The CFDE Knowledge Graph Working Group assembled datasets and data products from eleven CFDE DCCs aligned with the ontologies and standards chosen by the Ontology Working Group into a unified analytic knowledge graph called the Data Distillery.

Additional standardized data and metadata formats have been developed primarily by individual CFDE DCCs with financial support from CFDE. These DCC initiatives promote broader integration of common data types across multiple CFDE DCCs and other non-CFDE programs. For example, in support of applications including multiscale reference atlas construction and exploration (Figure 5), a common coordinate framework (CCF) for Human Reference Atlas (HRA)^29^ anatomical data was developed. The international HRA Working Group, led by HuBMAP and SenNet, created the CCF that captures the partonomy of 4,800+ anatomical structures (linked to UBERON ontology)^30^ in more than 30 organs and the 1,300+ cell types located within them (linked to Cell Ontology), plus 2,070+ biomarkers (linked to HGNC) used to characterize those cell types.^29^ The HRA CCF has been used to spatially register tissue data and to explore single-cell and anatomical data from HuBMAP, SenNet, HTAN, SPARC, Kids First, and many other projects (Figure 6).^31,32^

**Figure 5.**
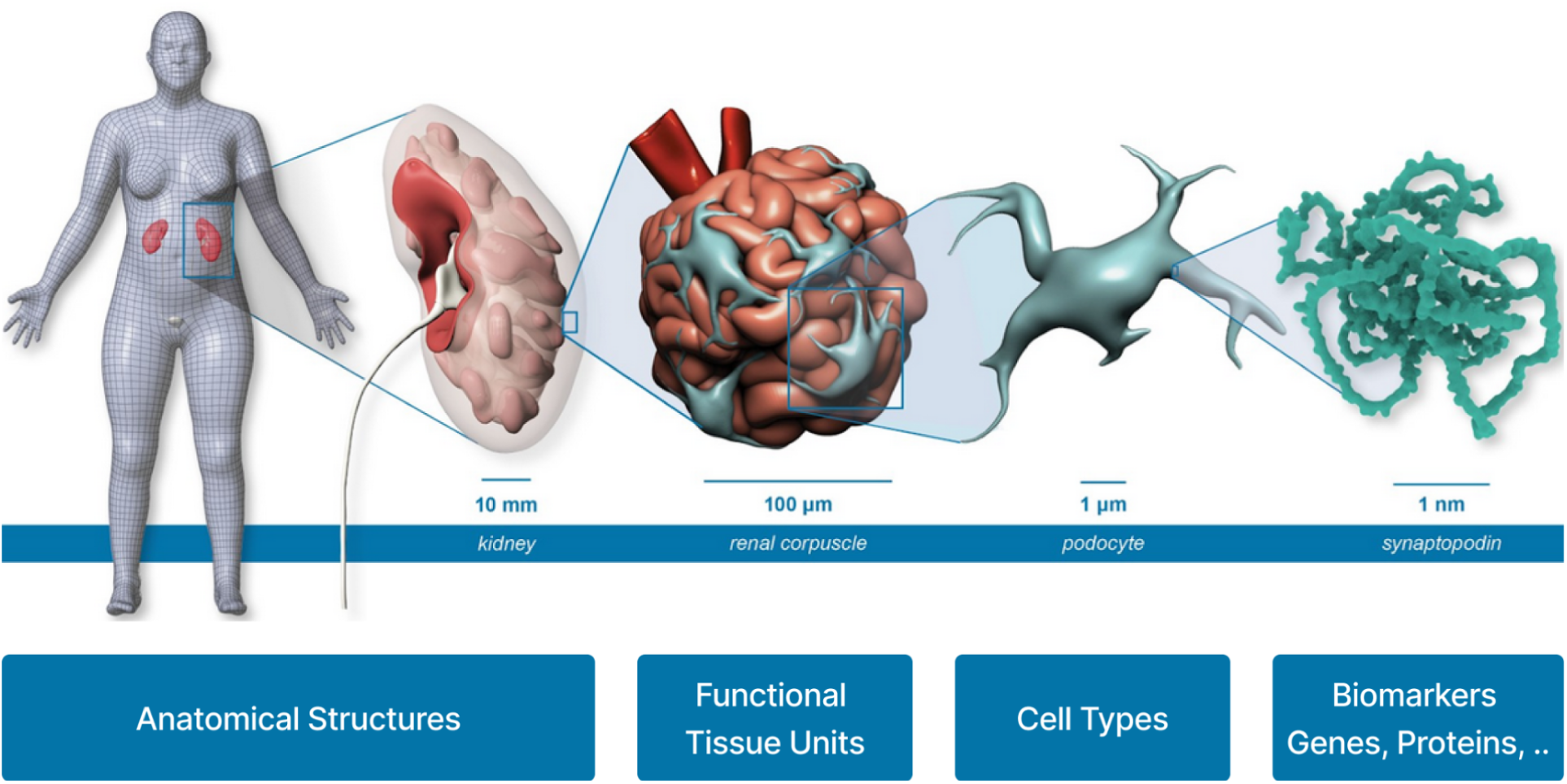
Multiscale data and user interfaces. The HRA Common Coordinate Framework (CCF)^85^ makes it possible to register and explore data in three dimensions and across scales. The HRA Organ Gallery in virtual reality (VR)^76^ supports multiscale exploration of 3D reference organs and HRA-harmonized single-cell datasets via head-mounted VR devices. VR supports interactions with HRA Digital Objects (DOs) via natural input using handheld VR controllers and enables the user to traverse the human body in an elevator-style system, from whole organs to anatomical structures, to functional tissue units (FTUS), to cell types, and down to sub-cellular structures, such as gene, protein biomarkers, see Figure 6. Abbreviations: CCF-Common Coordinate Framework, DO-digital object, HRA-Human Reference Atlas, FTUS-functional tissue units, VR-virtual reality.

**Figure 6.**
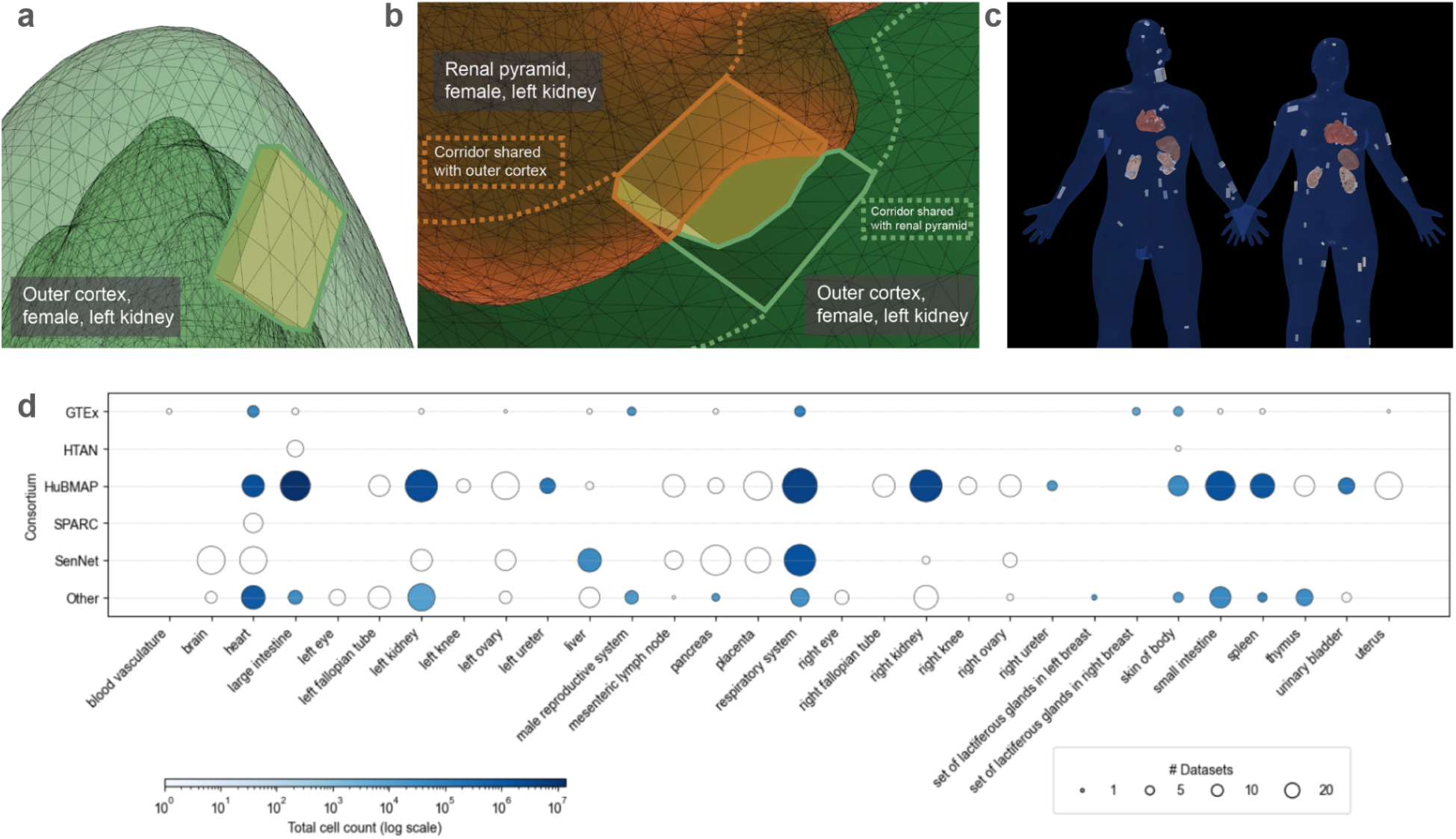
Tissue registration using the HRA Registration User Interface (RUI). **a.** Tissue registration into kidney anatomical structures using the CCF. Using the Registration User Interface, the size, position, and rotation of the tissue block (yellow cube) are adjusted until they present the properties of the tissue (e.g., biopsy) extracted from a human donor. Tissue collides completely with one anatomical structure (here the ‘outer cortex’ of the female left kidney). **b.** Example of a tissue registration in which the virtual tissue block collides with two anatomical structures (here ‘renal pyramid’ and ‘outer cortex’ of the female left kidney). **c.** All registered tissue blocks and the more than 8,500 associated datasets can be explored in the HRA Exploration User Interface. **d.** This graph shows the number of CFDE and other datasets that have been registered into the Human Reference Atlas either via 3D registration of tissue into the CCF or via crosswalking cell type annotations via HRA-compatible transcriptomics tools (e.g., Azimuth) or spatial omics tools (e.g., DeepCellTypes). The horizontal axis lists 30 anatomical structures while the vertical axis lists CFDE and other projects. Circle size denotes the number of datasets and circle color (log scale) denotes total cell count. Abbreviations: CCF-Common Coordinate Framework, CFDE-Common Fund Data Ecosystem, DO-Digital Object, GTEx-The Genotype-Tissue Expression Project, HRA-Human Reference Atlas, HTAN-Human Tumor Atlas Network, HuBMAP-The Human BioMolecular Atlas Program, FTUS-Functional Tissue Units, RUI-Registration User Interface, SenNet-Cellular Senescence Network, SPARC-Stimulating Peripheral Activity to Relieve Conditions, VR-Virtual Reality.

### Roles of standardized metadata formats in AI-readiness and lessons learned from the CFDE approach to metadata standardization

To build toward AI-readiness,^33^ CFDE has more recently encouraged DCC adoption of Croissant,^16^ a metadata format developed by an industry consortium to specify important information about datasets required for AI analytics. Thus far, Croissant has been implemented by some DCCs (Bridge2AI, GTEx, LINCS, and MoTrPAC) for subsets of data and is expected to be applied to more datasets from these and other DCCs in the future. Beyond Croissant, AI-readiness includes specifying the provenance of datasets and software and detailed characterization of statistics, semantics, recommended and prohibited use cases, ethics, and computability. CFDE has not yet adopted an ecosystem-wide set of practices to address these additional facets of AI-readiness.

In large ecosystems like CFDE, two disparate approaches may be applied to metadata formatting requirements. An ecosystem may adopt few formats^34^ or may alternatively adopt multiple formats.^35^ Benefits of the first model include standardization of practices in a common framework and reusability of tools, but barriers exist to representing multi-program data types, data models, data elements, and ontologies within a single framework. Benefits of the second model include the potential for increased community adoption, with a tradeoff of reduced interoperability. CFDE has favored the first approach, aligned with its goals of increasing interoperability across the ecosystem, but at times has engaged in judicious use of the second approach when the benefits of accessibility on disparate platforms outweigh the drawbacks (e.g., knowledge graph accessions, API definitions). Each of these formats is a “standard” within a specialized community and enables additional use cases and applications of the data.

### Knowledge graphs and infrastructure for data integration

A major goal of CFDE is to catalyze scientific hypothesis generation through integrative analyses that combine datasets generated by Common Fund programs with community-developed tools and data from other public resources. To achieve this goal, members of the CFDE community have developed infrastructure for data integration and hypothesis generation. The CFDE DRC, CWIC, and KC provide centralized locations to search, browse, and visualize high-level insights from datasets across DCCs and beyond, as described below.

To facilitate downstream analyses, datasets are processed into higher-order abstractions, such as knowledge graphs, gene set libraries, and attribute tables. CFDE knowledge graphs include the CFDE Data Distillery Knowledge Graph, ReproTox-KG, the BiomarkerKB Knowledge Graph, CFDE-GSE, Petagraph, and the HRA Knowledge Graph.^13,14,19,20,36^ The CFDE Data Distillery Knowledge Graph (DDKG; https://github.com/nih-cfde/data-distillery)^14^ currently integrates over 180 ontologies and controlled vocabularies and data from 11 CFDE DCCs and several non-CFDE programs, enabling queries across disparate biomedical dataset types.^14^ ReproTox-KG is (https://maayanlab.cloud/reprotox-kg) a knowledge graph that enables exploration of genetic or toxicologic mechanisms underlying structural birth defects.^13^ By leveraging similarities in structures and gene expression signatures of small molecules, ReproTox-KG can be used to predict drugs’ placental crossing capabilities and teratogenic risk. The BiomarkerKB (https://biomarkerkb.org) partnership has recently developed a standardized data model and controlled vocabulary for biomarkers, representing over 200,000 biomarkers and 1,000 phenotypes in multiple formats including a DDKG-integrated knowledge graph.^19^ The CFDE Gene Set Enrichment (https://gse.cfde.cloud/) provides an integrated enrichment analysis from gene set libraries derived from eight Common Fund programs.^8^

Some of these knowledge graphs were designed with interoperability in mind, and integration of many is ongoing. DDKG-UI^14^, ReproTox-KG^13^, BiomarkerKG, ProKN, and CFDE-GSE^8^ leveraged a generic KG user interface (KG-UI) template^37^ that was developed by the DRC,^37^ which uses Cytoscape as a main component for the frontend user interface and the Neo4j graph database platform (Neo4j, Inc.; https://neo4j.com/) for the backend KG database.

The Unified Biomedical Knowledge Graph (UBKG) serves as a base infrastructure that allows many assertions and entire graphs to be ingested for use across CFDE, with DDKG being the prime example that has used it (see use case section below). The CFDE Knowledge Graph Working Group is currently working on an exchange format standard consisting of a second version of UBKG with exchange in JSON, which will facilitate interoperable *ad hoc* assembly of assertions and knowledge graphs. CFDE’s UBKG-based knowledge graphs include DDKG, Petagraph, and the BiomarkerKB Knowledge Graph. Petagraph is an early version of DDKG,^20^ and the BiomarkerKB Knowledge Graph uses the same format and technology as DDKG and is integrated into DDKG in addition to a standalone format.^19^ ASCT+B, a part of the HRA Knowledge Graph, is also integrated with DDKG.^36^ The Protein Knowledge Network (ProKN; https://research.bioinformatics.udel.edu/ProKN) represents another UBKG-aligned knowledge graph, providing protein-centric data integration and use cases, leveraging the UniProt knowledgebase and associated resources. A DDKG-compatible version of C2M2 is currently being explored. The DDKG, Petagraph, HRA Knowledge Graph, and other non-CFDE KGs have previously been visualized through a bimodal network to compare the ontologies imported and served by these KGs.^36^ CFDE knowledge graphs have been interlinked to run novel queries and support multiscale data visualization and exploration (Figure 5), and machine learning analyses of CFDE knowledge graphs have generated new gene-disease association hypotheses across a dozen conditions.^38^

### CFDE collaborations with other consortia

CFDE frameworks for integration and analysis support cross-program analyses of CFDE data and, in some cases, enhance connections between CFDE and other large global repositories. For instance, the UniProt-CFDE partnership facilitates connections between protein-level data from UniProt^39^ and other resources and genome-centric data from CFDE. This partnership is developing the Protein Knowledge Network (ProKN), a protein-centric multi-omic integration knowledge graph framework building on and complementing the DDKG. The ProKN combines CFDE and non-CFDE data at the genome, protein, drug, and phenotype levels to facilitate functional genomics discovery. Similar connections with the Knockout Mouse Phenotyping Program (KOMP2)^40,41^ and the Bridge2AI Cell Maps for AI^25^ project (CM4AI) and the Bridge2AI Bridge Center are enabling connection with additional modalities. Such partnerships also leverage new tools such as KSMoFinder,^42^ a supervised neural network classifier that predicts kinases of human phosphosites at the motif and substrate levels for phosphoproteomic data analysis.

Partnerships between CFDE and non-CFDE teams can enrich the landscape for interoperable biomedical analyses. For example, the Metabotyping Humans partnership project spans the Metabolomics Workbench (MW),^26^ Molecular Transducers of Physical Activity Consortium (MoTrPAC),^43,44^ Human Health Exposure Analysis Resource (HHEAR),^45^ and Nutrition for Precision Health (NPH)^46^ programs and focuses on metabotyping. A metabotype defines a collection of metabolites or metabolite classes associated with a group of individuals based on the metabolome of humans conforming to a given phenotype. The project team is harmonizing metabolomics data across the three centers to identify metabotypes and facilitate functional interpretation. The team is developing tools that will allow exploring how perturbations such as exercise, diet, or environmental exposures shift homeostatic metabolic states.

The Gene Regulatory Working group engaged the NHGRI Clinical Genome Resource (ClinGen) project to develop and implement a registry of regulatory elements and standards for sharing gene regulatory information. This two-way collaboration leveraged ClinGen infrastructure while harnessing information from CFDE projects to identify disease-causing gene regulatory variants revealed by genome sequencing, which is a core focus of ClinGen.^15^

### Documenting data, pipelines, and standards

Communities often develop internal practices that provide information necessary to understand the key variables of an experiment or analysis without becoming overly cumbersome. As a community-of-communities, CFDE has not adopted a single ecosystem-wide approach or platform for developing and running computational pipelines or experiments. However, CFDE has developed guidelines for how data resources and experimental methods are documented, and metadata collection is shareable, public, modular, and combinable.

CFDE itself has developed and centralized multiple publicly accessible user guides, protocols, and standards (https://data.cfde.cloud/documentation). Moreover, as of March 2026, the CFDE community has developed 11 distinct recommendations for policies, standards, and design principles pertinent to an array of needs within the ecosystem (https://github.com/nih-cfde/rfcs/blob/master/adoptionstatus.md). For example, the CFDE Playbook Partnership^21^ was a two-year partnership project that involved the Data Resource Center and six DCCs. Together, partnership members created a visual platform to construct workflows from building blocks called metanodes. Each metanode can be an input form, an executable function like an API call, or an output such as a figure or a table. By combining metanodes, many workflows can be generated without the need for coding. The members of the partnership developed over 20 use cases that demonstrate how the system can perform on-the-fly analyses that integrate data from multiple Common Fund programs to form novel hypotheses or enrich users’ input data. To support this accomplishment, the partnership established standards for the development of interoperable APIs that could be used by multiple Common Fund DCCs in support of modular workflow development (https://github.com/nih-cfde/rfcs/blob/master/adoptionstatus.md).

Common Fund DCCs have also developed internal processes for documenting data and pipelines. While these are not CFDE products per se, CFDE does support the reuse and sustainability of these resources. Moreover, CFDE recognizes that some of these DCC-developed resources could be adopted more broadly across other resources to improve reproducibility and interoperability both within and outside of CFDE. Thus, a few examples are showcased here.

As one example, the GlyGen DCC approaches documentation of data and pipelines through the use of BioCompute Objects (BCOs).^47^ All data ingested into GlyGen are stored as datasets at data.glygen.org. For each dataset, a BioCompute Object (BCO) is created that conforms to the IEEE BCO specification^48^ and has been adapted for database and knowledgebase resources.^49^ These dataset-level BCOs are generated from the perspective of the integration workflow, ensuring that every processing step and its associated metadata are captured. The resulting BCO serves as a “readme” for the dataset, providing precise details on how the data were integrated. By adopting the BCO standard, GlyGen ensures granular tracking of metadata, particularly provenance, which supports proper attribution, license compliance, workflow exchange between researchers, and reproducibility. All dataset BCOs are recorded in machine-readable JSON format and are available for viewing and download at the GlyGen data portal (data.glygen.org). As datasets are updated, their corresponding BCOs are also revised and versioned. Everything visible on the GlyGen frontend (glygen.org) is documented within BCOs. This extends beyond data provenance to include software provenance, input/output files, and where needed embedded workflow languages (e.g. CWL) that enable automated execution. Together, these practices not only enhance reproducibility and transparency but also support long-term sustainability, so that even if original resource providers can no longer maintain the resource, others can continue its use and development. Wider adoption of BCOs across resources both within and outside of CFDE could provide a robust standard for workflow documentation, transparency, reproducibility, and interoperability.

Among other DCCs, the MoTrPAC bioinformatics center harmonizes data across multiple submitters, projects, and -omics analysis approaches to facilitate downstream analyses. A series of quality control and metadata harmonization scripts (https://motrpac-data.org/code-repositories) prepare data for downstream normalization and analysis. HuBMAP (https://docs.hubmapconsortium.org/data-submission/Section6.html), SenNet (https://docs.sennetconsortium.org/data-submission/Section6.html), and SPARC^50^ have established rich metadata and directory structure schemas with required and optional fields that all members of respective consortia must adhere to when submitting datasets. The metadata and directory schemas function as reference frameworks that allow for cross-program data findability and reuse. Datasets are validated against these schemas to promote data interoperability, reusability and compatibility with standardized pipelines. Central HuBMAP analysis pipelines are built with Docker containers in a Common Workflow Language (CWL) workflow to support reuse and reproducibility of these processing pipelines. KidsFirst uses CWL and Nextflow workflows and tools to enable reproducible bioinformatics. Currently, over 1000 community-sourced workflows are available for use via Cavatica (www.cavatica.org). Cavatica is also used by multiple DCCs across CFDE (e.g., LINCS, GlyGen, Kids First, exRNA)^51,52^ in the CFDE Playbook Partnership (https://playbook-workflow-builder.cloud/playbooks).^15,21^

### Tools and interfaces for data analysis, visualization and discovery

Multiple CFDE initiatives target improving cross-program data harmonization and visualization within the ecosystem. One example is the Human Reference Atlas (HRA; https://humanatlas.io), which combines multiscale datasets and donor metadata from multiple CFDE programs (HuBMAP, SenNet, and GTEx; Figure 5). This supports construction of cell atlases, and harmonized reference data is available via the HRA Knowledge Graph (KG).^29,36^ There are multiple tools for web-based access to the HRA, including the HRA KG Explorer (https://apps.humanatlas.io/kg-explorer), spatial examination via the Exploration User Interface (EUI; https://apps.humanatlas.io/eui; Figure 6c),^53^ with cell distance distributions analyzable through the Cell Distance Explorer (CDE, https://apps.humanatlas.io/cde)^31^ and immersive integration and visualization of healthy donor cell types available through the HRA Organ Gallery (https://humanatlas.io/hra-organ-gallery).^29^

Based in part on these integrative, collaboratively-developed tools, multiple CFDE Centers, in particular the Data Resource Center (DRC; https://info.cfde.cloud/)^8^ and the the Knowledge Center (KC; https://cfdeknowledge.org), build toolkits for constructing integrative resources. The ultimate goal is to enable users who lack deep computational expertise to construct similar integrative analyses through intuitive web-based interfaces.

The DRC developed the tools GeneSetCart^54^, GDLPA^55^, CFDE-GSE, the Playbook Workflow Builder (PWB)^21^, Rummagene^56^, RummaGEO^57^, and GSFM^58^, all centered around the abstraction of omics datasets from over 10 Common Fund CFDE programs, and external resources, into gene sets, and the process of combining gene sets from multiple Common Fund programs for cross DCC knowledge discovery and hypothesis generation. Furthermore, the DRC’s CFDE Workbench^8^ provides an entry point for DCCs to submit processed data and metadata about their data, and a mechanism to search and discover data, processed data, knowledge graph assertions, and tools developed by the DRC and CFDE partnerships, applicable to items discovered on the workbench. The CFDE Workbench’s search functions are exposed via an API and Model Context Protocol (MCP) server, leveraged by the CFDE Workbench chatbot to find DCC-submitted knowledge and trigger PWB workflows to respond to user queries.

The KC provides a centralized, user-friendly web entry point for the ecosystem and promotes novel disease-focused discoveries and integrative use cases. Users can gain a broad understanding of the ecosystem through curated summaries of each CFDE DCC, including its purpose, data, resources, and key publications. The KC routinely attends conferences and hosts a regular webinar series (https://cfdeknowledge.org/r/CFDE_webinars) to promote broader visibility into and integration of CFDE resources. A summary of flagship outreach activities by the KC and other centers is provided in Table 5.

**Table 5.**
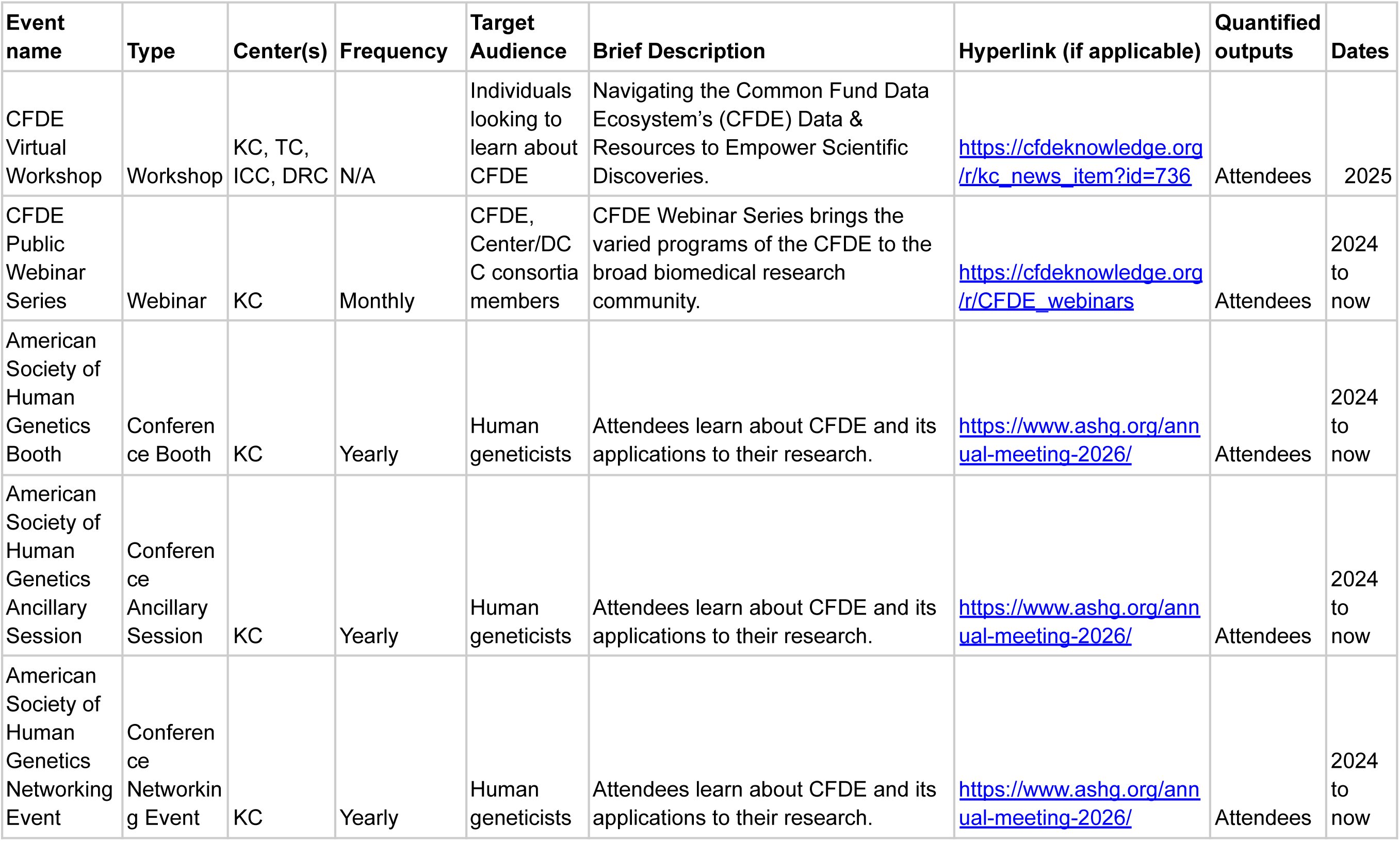
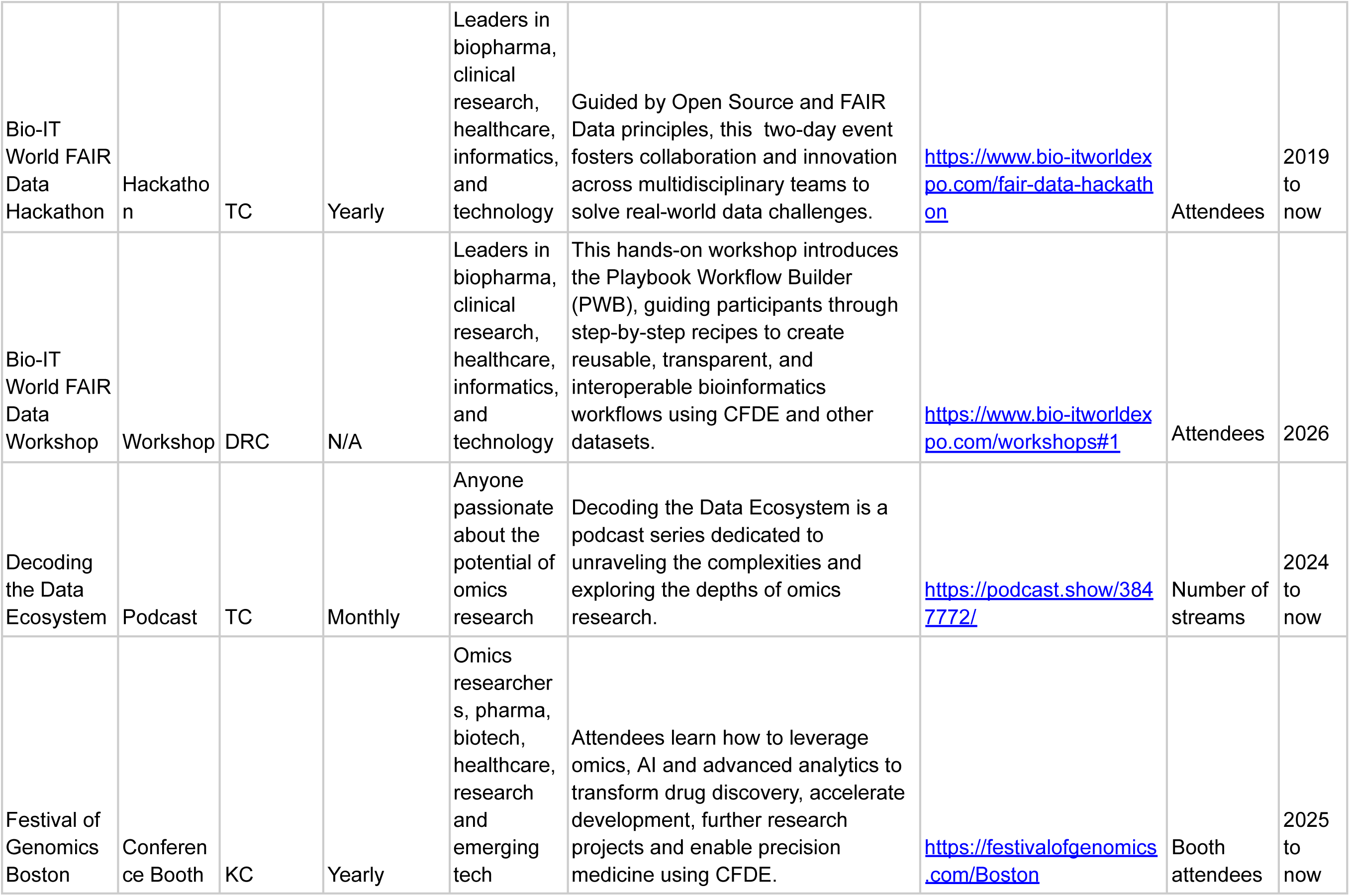
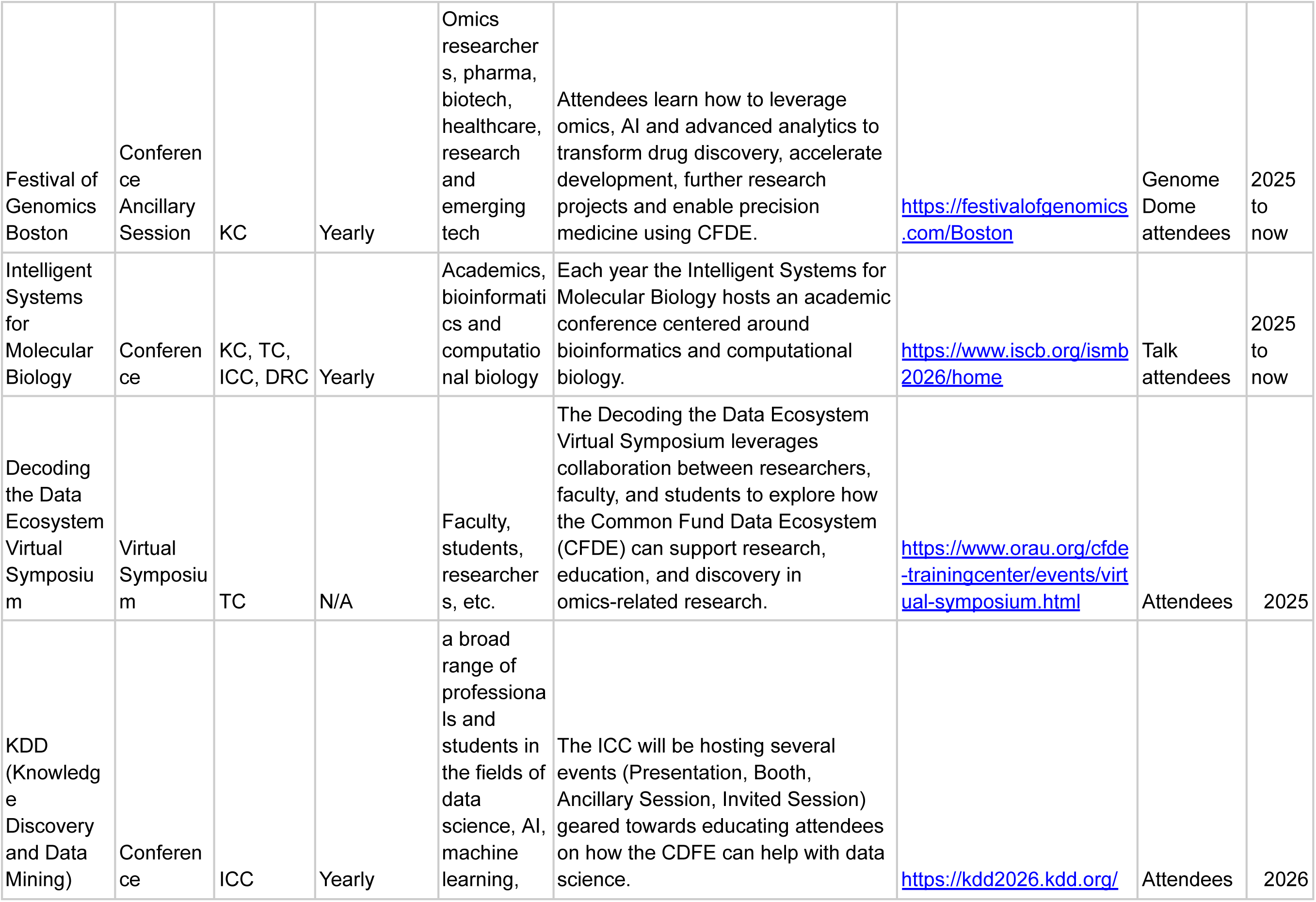
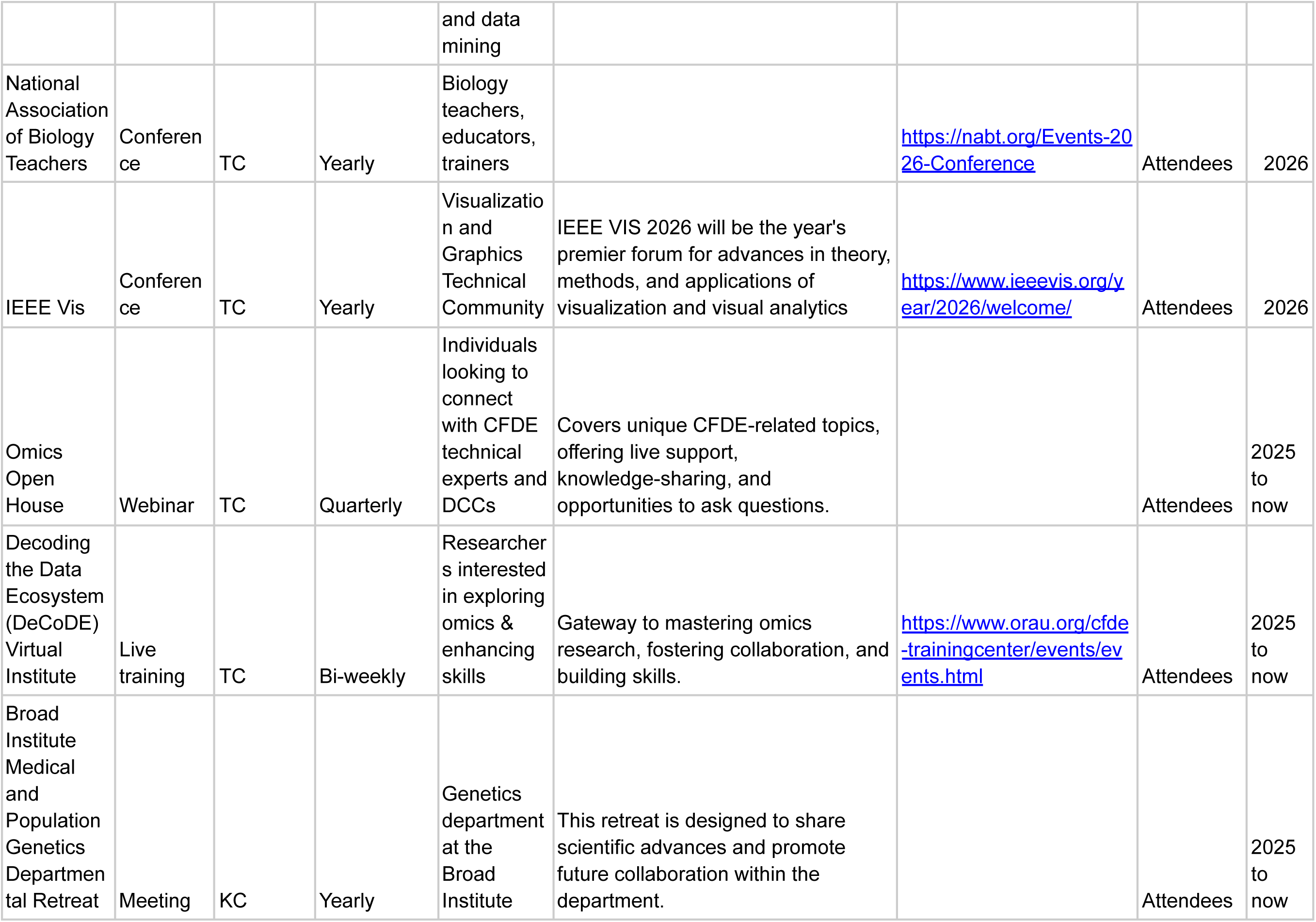
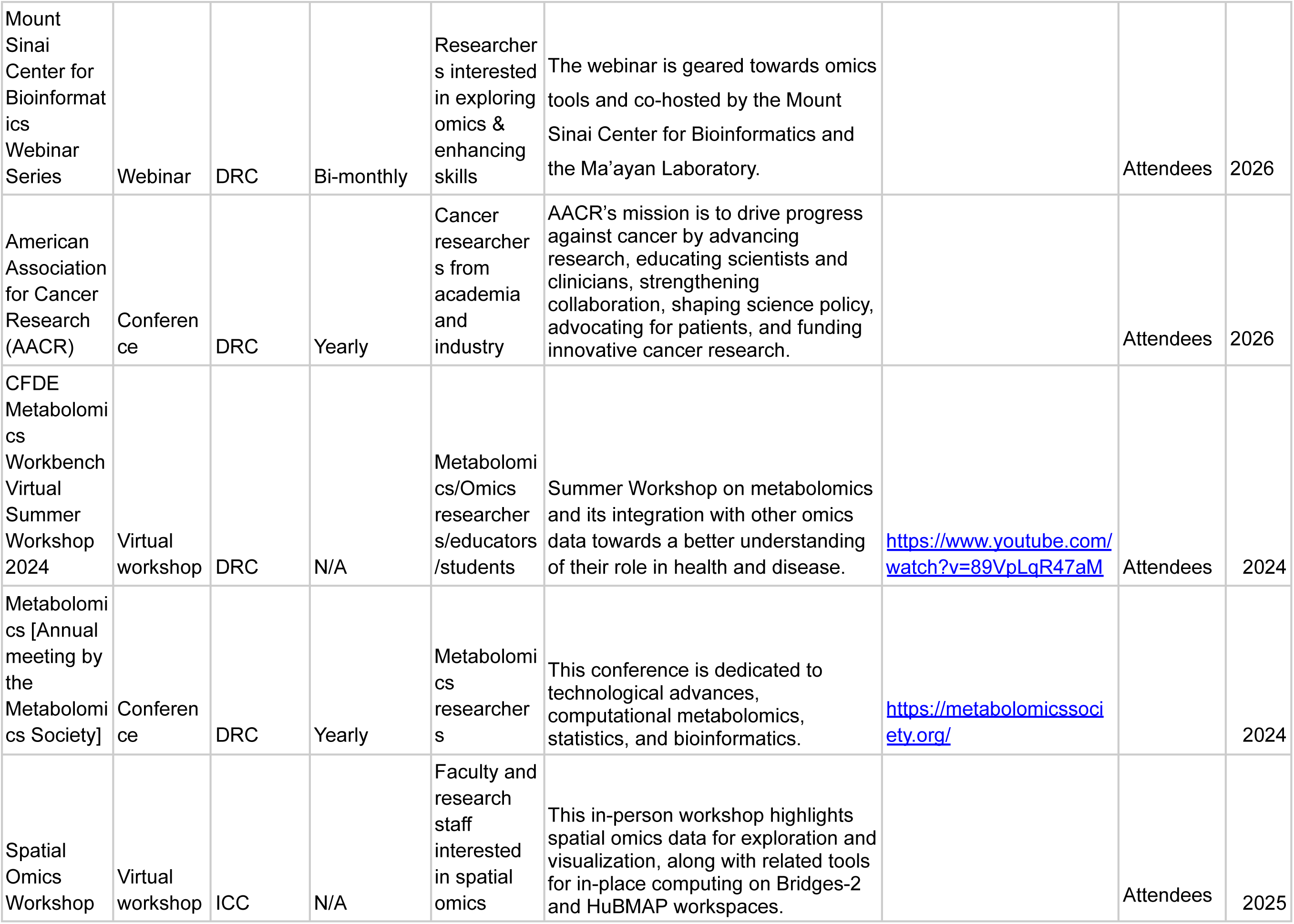
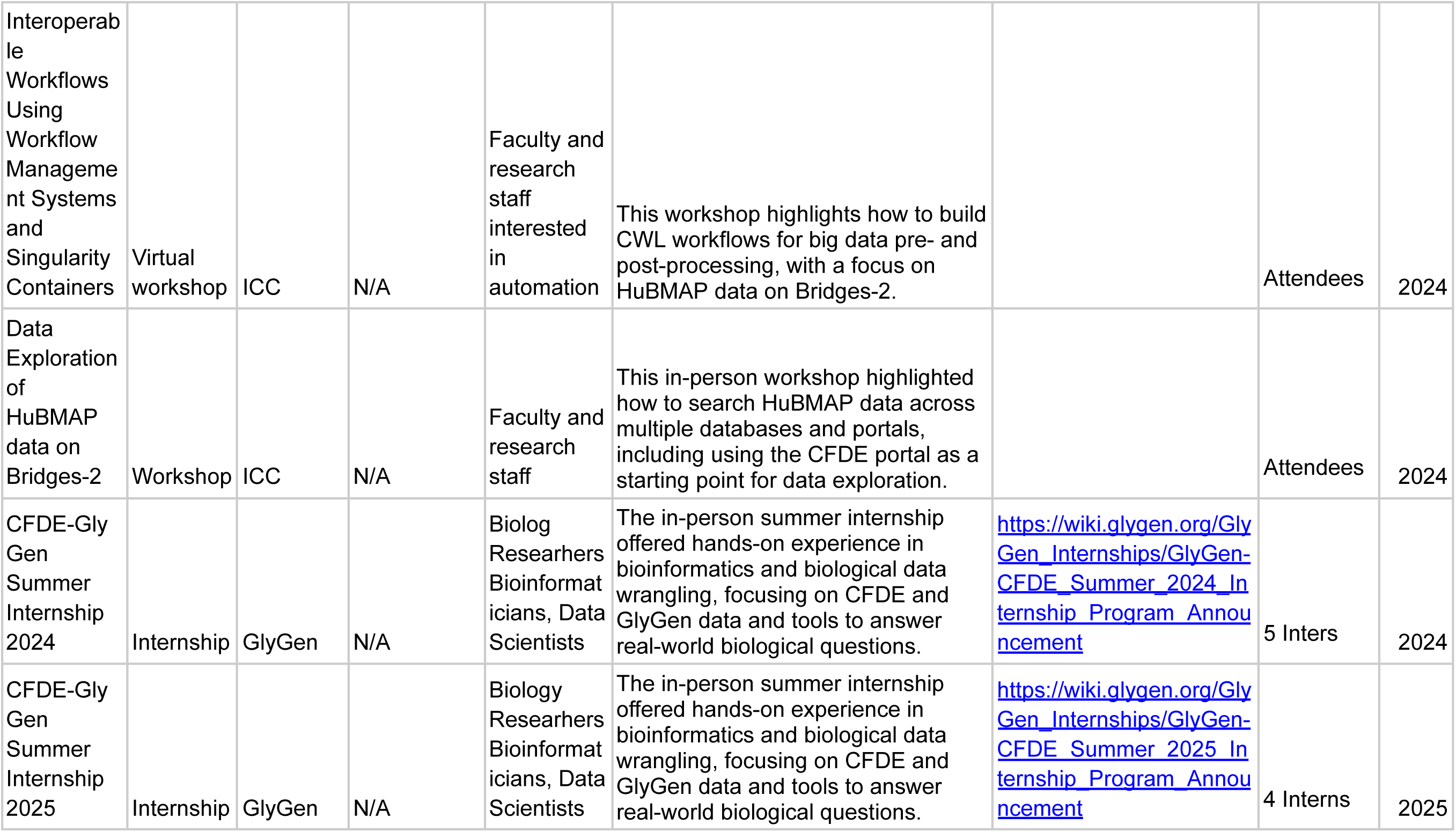
Flagship CFDE outreach and engagement activities. Key outreach and engagement activities undertaken by CFDE centers since their activities began in earnest in 2024. Abbreviations: DRC-Data Resource Center, ICC-Integration and Coordination Center, KC-Knowledge Center, TC-Training Center.

Given the expansive range of data and information across CFDE, it is important to help users identify the data, information, or analytical results most relevant to them. To guide users toward datasets or analyses of interest, the KC has implemented large language model (LLM)-assisted workflows, CFDE-REVEAL and CFDE-DESIGN, whose current implementation leverages the OpenAI GPT5-mini LLM model. CFDE-REVEAL (https://cfdeknowledge.org/r/cfde_reveal) is an interface that converts users’ questions into mechanistic hypotheses. From user-supplied natural language queries, CFDE-REVEAL performs semantic search across statistical associations of gene sets and traits from the KC Gene Set Browser (https://cfdeknowledge.org/r/kc_gsb). An LLM then reveals candidate hypotheses and mechanisms derived from these associations. For example, a user might ask, “What are druggable targets for obesity?” REVEAL extracts search terms from the query, returns the top phenotype-gene signature associations across CFDE programs, and uses these associations to generate mechanistic hypotheses such as identifying a microRNA-mediated growth and extracellular remodeling axis that connects BMP/SMAD signaling to hip circumference. Specific candidate genes are proposed (e.g., *LTBP3, HMGA2, HSPG2*), and the underlying phenotype-gene signature associations and gene set clusters are provided that support each gene. These hypotheses or independent gene set-trait associations can then be supplied to a second KC workflow, CFDE-DESIGN (https://cfdeknowledge.org/r/cfde_design), to produce customized LLM-generated experimental protocols.

### Cloud computing infrastructure

Conducting integrative analyses can be challenging when the data reside in different environments. CFDE’s standardized metadata and tools aligned with community practices create the opportunity for a cloud computing environment capable of running analyses that integrate data from the underlying data repositories. The Cloud Workspace Implementation Center (CWIC) (https://cloudworkspace.org) provides a unified, scalable, and secure computational environment designed to aid interoperability among Common Fund resources to support a broad range of biomedical researchers and data scientists. Built on the Galaxy platform (https://usegalaxy.org),^59^ CWIC offers a familiar interface that anchors a suite of integrated tools and environments. Users access CWIC using their free account that supports two-factor authentication and role-based access controls. This access is managed by the Texas Advanced Computing Center (TACC) application programming interface (Tapis).^60^ Once logged in, users can access Common Fund datasets, tools, and workflows through the feature-rich, zero-code Galaxy graphical user interface and launch interactive sessions in Jupyter and RStudio. As TACC users, individuals can also directly access high-performance computing (HPC) resources using command-line interfaces. This direct access enables advanced customization, including the selection of compute nodes, the installation of specialized packages, and the configuration of environment parameters tailored to specific workflows. Users access commercial cloud resources through CloudBank (https://www.cloudbank.org).

CWIC supports both CPU and GPU-based workloads, particularly for compute-intensive AI workloads. To minimize barriers to data-intensive research, CWIC minimizes data transfer and egress costs, aiming for seamless integration with external data repositories and services. Features such as workflow sharing, provenance tracking, and usage analytics enable users to effectively manage and optimize their computational resources. User dashboards enable usage, promoting transparency and supporting informed resource management. Furthermore, by leveraging several major GA4GH standards,^61^ such as the Directory Registry Service (DRS) and Tool Registry Service (TRS), CWIC enables users to easily integrate with datasets and resources from other biomedical cloud resources such as the NHGRI AnVIL^62^ or NHLBI BioDataCatalyst.^63^ Overall, CWIC is developed as an open-source platform, leveraging existing investments in Galaxy and related projects to support long-term sustainability and community-driven innovation.

### Training and workforce development

The CFDE Training Center (TC) serves as a hub for developing and implementing training programs that bridge knowledge gaps, improve access to CFDE resources, and foster collaboration among CFDE community members and other researchers. Building the TC required a detailed assessment of the current training landscape to understand the needs and challenges of the CFDE community. Specifically, the landscape analysis assessed the effectiveness of existing CFDE training programs, identified barriers that impact researcher engagement and learning outcomes, evaluated best practices from bioinformatics and data science training models, and developed evidence-based recommendations to improve CFDE training strategies. Recommendations included improving access to resources through a centralized platform for CFDE resources; enhancing training structure through modular, hands-on, self-paced courses; strengthening mentorship programs through “train-the-trainer” programs that equip mentors with effective teaching and communication skills; and expanding evaluation methods with pre- and post-training assessments to measure knowledge gains and real-time quizzes and coding exercises with instant feedback to reinforce learning (CFDE Training Team, personal communication).

The TC serves the CFDE community by building foundational and advanced omics analysis skills through a dynamic syllabus grounded in six aims for comprehensive development: understanding biological domains relevant to CFDE datasets; leveraging biological reference databases; automating computational tasks with scripting; using software and programming tools; applying appropriate statistical and visualization methods; and leveraging advanced computing and collaboration tools. Each aim includes clear learning outcomes structured around three cognitive levels—Remember, Understand, and Apply—to guide researchers from conceptual understanding to practical application (Figure 7a).

**Figure 7.**
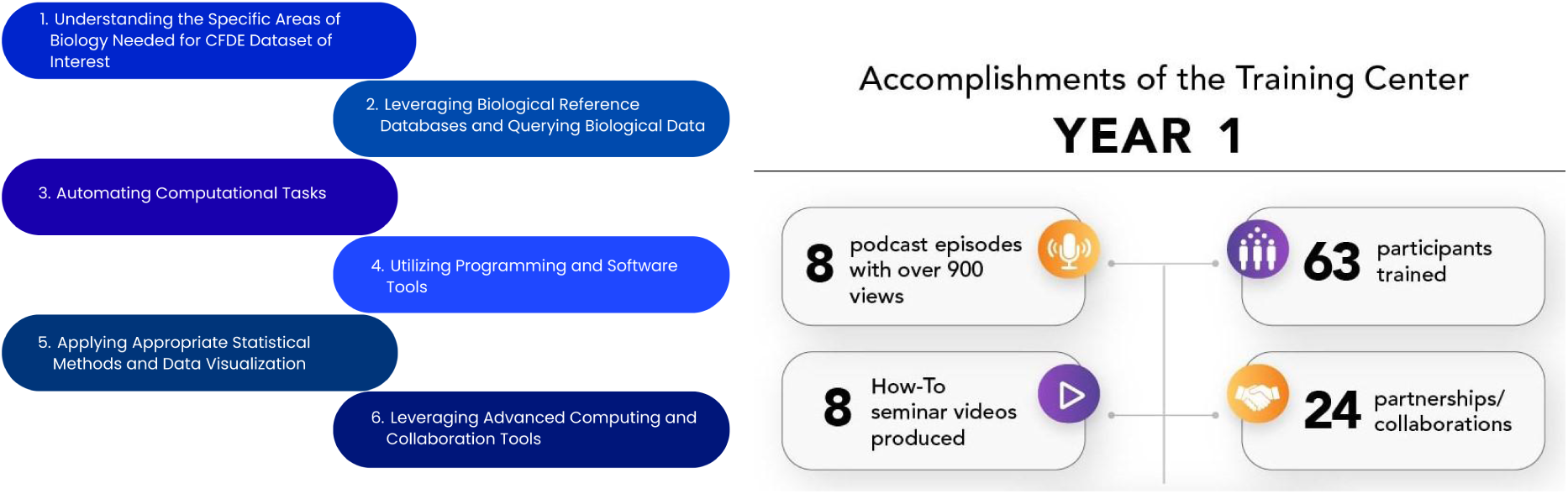
Training Center’s Aims for Omics Skills Development. **a.** The Training Center serves the CFDE community by building foundational and advanced omics analysis skills through a dynamic syllabus grounded in six comprehensive aims: understanding the specific areas of biology needed for CFDE datasets; leveraging biological reference databases; automating computational tasks; utilizing programming and software tools; applying appropriate statistical and visualization methods; and leveraging advanced computing and collaboration tools. **b.** In its first year, the TC has developed a wide range of resources and materials and hosted over 10 events. Events included Decoding the Data Ecosystem podcast (https://podcast.show/3847772/), the e-Learning Dashboard (which includes a How-To Seminar Series) (https://orau.org/cfde-trainingcenter/training/e-learning.html), a Digital Scavenger Hunt, and the Bio-IT World Hackathon (https://www.bio-itworldexpo.com/).

Together, the TC supports a cohesive and scalable learning framework for biomedical scientists. The TC has developed a wide range of resources and materials and hosted 10 events, such as the Decoding the Data Ecosystem podcast (https://podcast.show/3847772/) with over 900 views, the e-Learning Dashboard (which includes a How-To Seminar Series) (https://orau.org/cfde-trainingcenter/training/e-learning.html), a Digital Scavenger Hunt, and the Bio-IT World Hackathon (https://www.bio-itworldexpo.com/) (Figure 7b). By integrating these benchmarks into its initiatives, the TC has created an environment for the community that will empower researchers to leverage CFDE datasets and tools effectively while fostering collaboration and innovation. The TC has developed a robust program based on participant feedback to inform the development of new training resources and opportunities, while establishing partnerships with CFDE DCCs and other training and outreach efforts to adapt to changing needs.

### Coordination and Assessment

The CFDE Integration and Coordination Center (ICC) works across three core areas: administrative coordination, evaluation, and sustainability. ICC administers CFDE working groups and hosts semi-annual CFDE conferences, manages cross-CFDE communications, defines evaluation metrics and creates dashboards to help monitor CFDE to identify best practices and areas for growth, and develops and supports sustainability strategies.

The CFDE evaluation strategy encompasses three temporal scales of outcomes. Short-term metrics focus on resource availability, tracking the deployment and accessibility of datasets, software tools, web-based platforms, and training materials such as webinars and tutorials. Mid-term assessments emphasize user engagement through usage statistics, user satisfaction surveys, trainee feedback, and early indicators of resource sustainability. For example, across CFDE web properties steady growth in usage was observed through mid-2025 (40-50% growth rates), at which point access patterns, potentially driven by AI-associated systems, drove substantial growth (1972 engaged sessions per day in 2024; 6953 engaged sessions per day in 2025). Long-term impact evaluation examines transformative outcomes, including peer-reviewed publications describing or leveraging CFDE resources, sustainment efforts and funding, and qualitative or quantitative changes in research practice or workforce. This multi-tiered, shared responsibility model provides a comprehensive framework for understanding how data ecosystem infrastructure translates into scientific advancement and community transformation.

The ICC’s Evaluation Core has implemented automated workflows that supply NIH and stakeholders with standardized publication and citation metrics, social coding metrics (via GitHub), and Google Analytics. A centralized event and calendaring system forms the basis for the ongoing assessment of outreach activities, utilizing a shared survey approach. Automated collection of GitHub metrics, publication and citation metrics, Google Analytics, overall project budgets and funding timelines across funded CFDE projects, powers a dashboard that enables rapid examination of both ecosystem-wide and component-specific contributions.

The ICC Evaluation Core’s automated collection of GitHub analytics for CFDE-developed source code from December 5, 2025, indicated 127 public repositories from CFDE-supported projects, which had garnered 861 stars. Open licensing is often key for sustainability. Across the public repositories, most had explicit open source licensing with 54 repositories using MIT licensing followed by 22 GPLv3 licensing. A README file, important for community engagement for open source software, was available for 85% of repositories. Across all repositories, a total of 259 contributors had performed 108,614 commits and closed 14,767 pull requests. The TC utilized the Academic Analytics Research Insights database to examine research activities associated with 14 CFDE programs across 531 U.S. academic institutions. A total of 1,990 scholars from 195 academic institutions referenced at least one CFDE dataset in 4,438 publications. Finally, individual Common Fund programs demonstrate broad impact through their contributions to the scientific literature. For example, between 2013 and 2024, the exRNA Communication Program^64–66^ alone produced 877 publications that were cited 79,442 times.^67^

### Sustainability

The ICC’s Sustainability Core has developed a framework for improving the FAIRness, AI-readiness, and sustainable management of CFDE datasets through three guiding principles.^68^ The current implementation is informational as opposed to evaluative. First, dataset collection and management must be ethically sourced, with data privacy, safety, and integrity safeguarded throughout every operational step, from acquisition to curation and dissemination. Second, building trust must be understood as a collaborative process involving both data contributors and users, embedding co-design and stewardship principles that cultivate transparency and accountability across the data lifecycle. Finally, achieving transparency, provenance, and explainability in all tools and resources is essential to rendering datasets reproducible and sustainable over time. Together, these components embody a commitment to ethical stewardship that enables CFDE to serve as a foundation for responsible and trustworthy biomedical AI integration.

Sustainability considerations for CFDE include data, computing, code, and funding. With respect to data, the DRC accepts quarterly submission of different datasets, metadata, and code assets from the DCCs. These include dataset assets and code assets. Dataset assets include C2M2 data packages,^7^ knowledge graph assertions,^14^ gene set libraries, and attribute tables. Code assets include APIs, ETLs, landing pages templates such as gene, drug, and disease pages, and models. As of March 2026, the Data Matrix on the Data Portal contains 70 assets that include 15 C2M2 packages submitted by 15 DCCs. Submissions are assessed for FAIRness using FAIRshake^69^ (https://fairshake.cloud/) and vetted by both the DRC and the submitting DCCs. The FAIRshake evaluations are reported back to the submitter as feedback about what they can improve to make their assets more FAIR. FAIRshake evaluation checks for data formatting errors in addition to FAIRness through a complex evaluation rubric (https://fairshake.cloud/rubric/) that is inspired by the universal FAIR metrics^70^ but does not map to them precisely. While the DCCs have improved the FAIRness of their metadata, data, and code asset submissions over time, this is difficult to observe because over time new DCCs are introduced to the ecosystem and more assets are added to the Data Portal (Figure 8).

**Figure 8.**
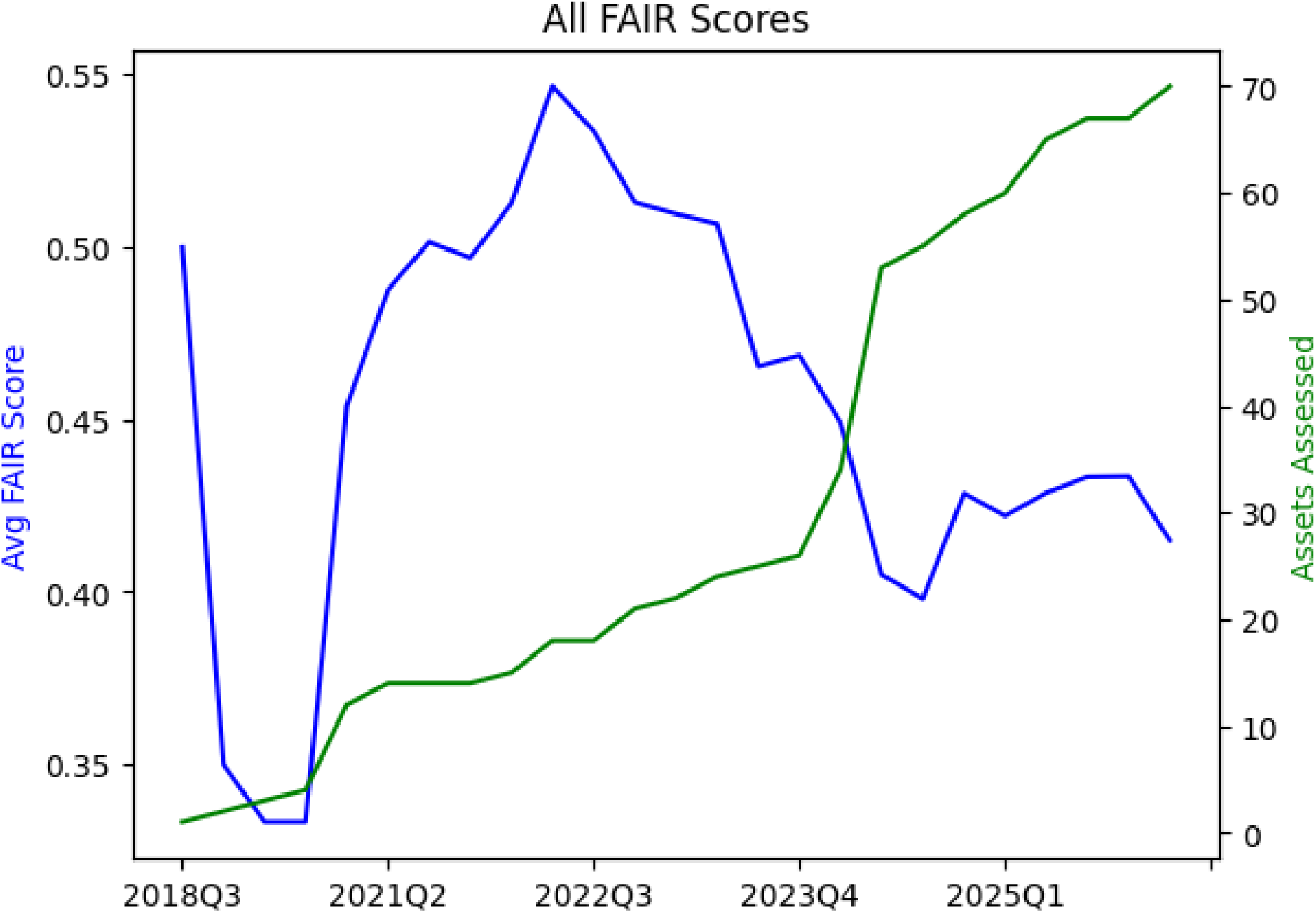
Change in average FAIRshake scores across all submitted assets from all DCCs over time. In March 2026, average FAIRshake scores were calculated across all submitted assets (n=70) from all DCCs (n=15) over time (years 2018-2026). These included dataset assets (C2M2 data packages, knowledge graph assertions, gene set libraries, and attribute tables) and code assets (APIs, ETLs, landing pages templates such as gene, drug, and disease pages, and models). Blue-average FAIR score; green-number of assets assessed.

CFDE has not yet fully addressed the funding component of sustainability, and discussions of how to address this remain ongoing. As a partial solution, the ICC has mapped key CFDE data types to potential repositories available to host them at the conclusion of awards, with the idea that CFDE data could potentially be integrated into these external repositories for sustainability (Figure 1; interactive diagram available at https://cfde-sankey.github.io/). Some DCCs have already selected a subset of these repositories to host their data given the repositories’ support of long-term sustainability. Sustainable access to computing is supported by the variety of computing resources available to users. TACC systems allow users access to 100K+ CPUs, 1K+ GPUs and 100s of petabytes of storage to allow for analysis and reproducibility of large-scale analysis. Additionally, users have access to commercial cloud computing tools via the Cloudbank partnership, which provides users access to all major cloud computing platforms. Collaborations between the KC, DRC, and CWIC enable users to identify datasets or construct workflows and then launch jobs directly onto the CWIC. The CWIC enables users to share their history, supporting the dissemination of workflows to easily reproduce analyses and validate data without having to re-conduct the underlying analyses.

### The value of an ecosystem: highlighted use cases and biological insights

Finally, CFDE aims to generate substantial community impact through three primary mechanisms: fostering cross-program collaborations among previously siloed Common Fund programs, establishing shared technical standards for metadata harmonization, and lowering barriers to entry for researchers unfamiliar with specific Common Fund program resources.

Use cases for cross-program collaborations arguably provide the most important insights for constructing analytical and data integration tools and demonstrating integrative data analysis. Most biomedical researchers develop hypotheses based on a mechanistic, conceptual, or analytical model, embellishing it with additional data and evidence. Common Fund projects offer best-in-class datasets and tools that can aid a biomedical researcher in developing novel biological hypotheses and outcomes.

As a demonstration of this cross-ecosystem discovery, here we present a use case from the Data Distillery Knowledge Graph^14^ using the Neo4j graph database platform (Neo4j, Inc.; https://neo4j.com/). We provide a Cypher query that captures mechanistic hypotheses that link genetic variation to disease through tissue-specific metabolic and transcriptional processes (Figure 9). The query follows a workflow initially proposed by the IDG Reactome Project (https://idg.reactome.org/) and was developed to extract a tightly constrained, semantically coherent subgraph from a biomedical knowledge graph by composing multiple relationship patterns across heterogeneous data sources. The query first anchors on a three-hop causal–correlational chain linking gene, metabolite, and condition concepts using source-qualified edges from Metabolomics Workbench (MW), then incrementally refines the result set by enforcing additional topological and semantic constraints. These include tissue-level provenance of metabolite production, independent evidence of gene expression from GTEx via intermediate expression entities, and shared or related tissue contexts connecting molecular activity to disease-relevant anatomy. After filtering the core graph pattern, we populate the resulting subgraph with low-cardinality annotations (preferred terms), upstream biochemical relationships from Illuminating the Druggable Genome (IDG) (gene–protein–compound), and quantitative expression metadata. This generated structure reflects a deliberate graph-querying strategy that balances expressiveness with performance while preserving provenance, semantic specificity, and interpretability of the extracted subgraph. The query is designed to use the information provided for tissue-specific diseases (utilizing content from the UMLS-based Unified Biomolecular Knowledge Graph (UBKG; https://ubkg.docs.xconsortia.org/))^20^ as a means to filter the results and provide more meaningful relationships.

**Figure 9.**
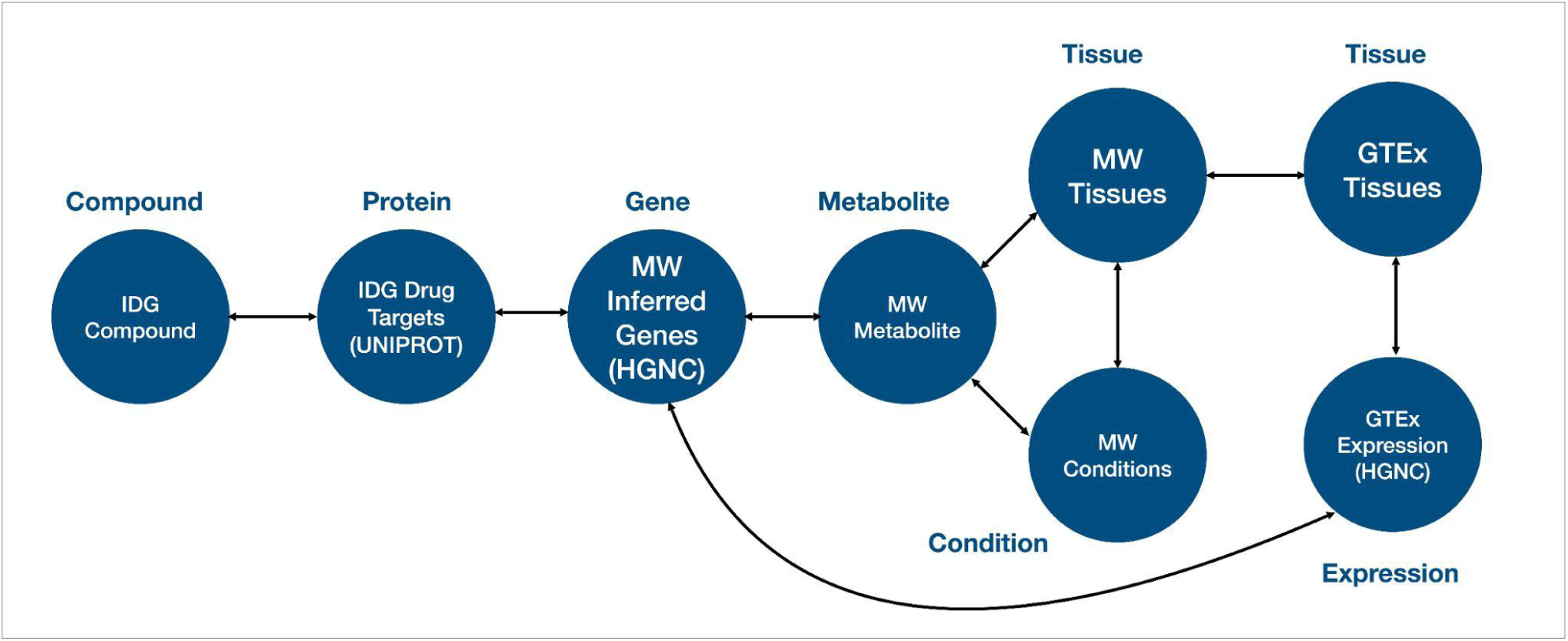
Highlighted use case from the CFDE Data Distillery Knowledge Graph: sample Cypher query and schema intersecting data from three DCCs. Schema for the Cypher query from the Data Distillery Knowledge Graph developed to obtain novel biological insights using data from three CFDE DCCs (GTEx, IDG, and MW). Abbreviations: CFDE-Common Fund Data Ecosystem, DCC-Data Coordinating Center, GTEx-The Genotype-Tissue Expression Project, HGNC-HUGO Gene Nomenclature Committee, IDG-Illuminating the Druggable Genome, MW-Metabolomics Workbench.

The results (Table 6) demonstrate sample outputs from this query, showing a candidate biological pathway in which a gene product (*MGAM*) influences a metabolite (sucrose) associated with a clinical condition (polycystic kidney disease). Both processes are grounded in a relevant tissue (kidney), where the sucrose levels may be influenced and *MGAM* is expressed. Finally, IDG suggests compounds associated with MGAM, thus highlighting candidate therapeutic targets for this pathway. While this example is exploratory and would require additional downstream supporting evidence, it does show the power of this interface to rapidly generate initial hypotheses for future validation by leveraging information from multiple CFDE programs.

**Table 6.**
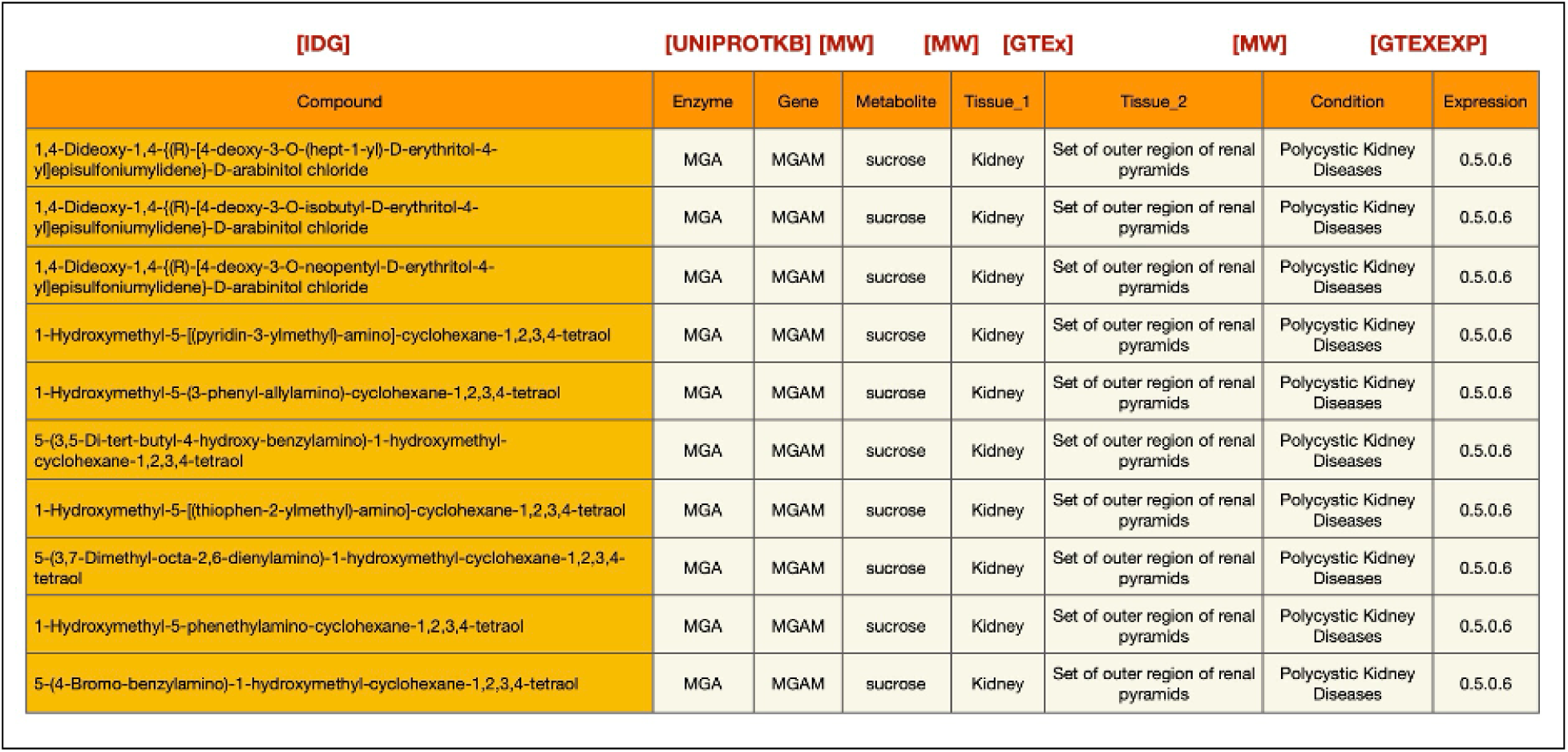
Highlighted use case from the CFDE Data Distillery Knowledge Graph: sample query results intersecting data from three CFDE DCCs. Sample outputs from a Cypher query conducted on the Data Distillery Knowledge Graph that nominates candidate mechanistic hypotheses for future validation. The query first connects gene, metabolite, and condition concepts using edges from Metabolomics Workbench (MW), then refines the results through additional constraints. These include tissue-level provenance of metabolite production, independent evidence of gene expression derived from GTEx via intermediate expression entities, and shared or related tissue contexts connecting molecular activity to disease-relevant anatomy. After filtering the core graph pattern, the resulting subgraph was enriched with low-cardinality annotations, upstream biochemical relationships from IDG (gene-protein-compound), and quantitative expression metadata. Results demonstrate a candidate biological pathway in which a gene product (*MGAM*) influences a metabolite (sucrose), and the metabolite is associated with a clinical condition (polycystic kidney disease). Processes are grounded in a relevant tissue (kidney) where the *MGAM* gene is expressed. IDG suggests compounds that are associated with MGAM, thus highlighting candidate therapeutic targets for this pathway. Abbreviations: CFDE-Common Fund Data Ecosystem, DCC-Data Coordinating Center, GTEx-The Genotype-Tissue Expression Project, IDG-Illuminating the Druggable Genome, MW-Metabolomics Workbench.

By integrating gene-metabolite-condition linkages from Metabolomics Workbench, expression evidence from GTEx, tissue anatomy from GTEx, and upstream chemical and protein interactions information from IDG, the extracted graph supports interpretation of how molecular regulation propagates across biological scales. This type of integrative representation is particularly valuable for elucidating disease mechanisms, prioritizing therapeutic targets, and identifying candidate biomarkers, as it contextualizes gene–disease associations within the metabolic, anatomical, and biochemical systems that underlie human pathophysiology. Because integrative tools from CFDE are still relatively new, efforts to disseminate these resources and demonstrate their uptake by the community are still ongoing.

### Challenges and solutions

CFDE shares certain challenges of its participating Common Fund-supported projects. These projects are ambitious, cross-disciplinary, and at the forefront of their fields. Therefore, nearly every project has elements that are both unique to CFDE and unique within their field. Across DCCs, priorities are often split between addressing the immediate needs of the project and the enabling work of the ecosystem, drawing focus away from generalizable solutions. CFDE Centers provide a mechanism to address this challenge by collaborating with DCCs on targeted initiatives. For instance, the CFDE KC and DRC are working with DCCs to develop and implement “The CFDE Navigation Wheel (Figure 1),” an interactive diagram of CFDE components that provides centralized access to the ecosystem across its many disparate web entry points. Finally, CFDE co-evolves alongside Common Fund programs to extend to broader data types, modeling, and simulations.

While CFDE aims to evaluate FAIR principles^71^ systematically through FAIRshake^69^ and to encourage the adoption of FAIR principles among DCCs, the uptake of FAIR principles across the ecosystem remains ongoing, and the ecosystem aims to balance promoting FAIRness with the timely sharing of data to encourage their uptake even if they are not yet completely FAIR. While this challenge is not unique to the CFDE, each ecosystem has its own unique needs that require tailored approaches to improving FAIRness over time.^72,73^ A previous study^74^ analyzed the FAIRness of 28 major biomedical databases, 4 of which were CFDE programs (GTEx, HuBMAP, LINCS, and Metabolomics Workbench), using a 29-question rubric. For each of the 29 questions, one point was assigned if the information could be identified from the database to answer the question. Scores across the 28 databases ranged from 7 to 29 points, and the mean FAIRness score for CFDE databases was higher than for all databases (24.25 points for CFDE databases; 18.57 points for all databases), and the only database to score 29 points was the CFDE program HuBMAP. However, most CFDE databases had room for improvement across the FAIR spectrum.

Sustaining the large scale Common Fund data resources is nontrivial, as each DCC has developed its own architecture for consent and data access. The technical solutions used by individual DCCs often reflect the state of the art in their field at the time of DCC development, but many require engineering and informatic liftover to meet interoperability standards and best practices. Research objectives and data types also range across and within DCCs, and nearly any type of biomedical data referential to nearly any organ, tissue, molecule, disease, experimental approach, organism, or analytic approach may be part of a CFDE DCC. Thus, data harmonization is required across the same and distinct data modalities, but integration must focus on the practical to remain feasible, and many data elements may ultimately not be harmonized. Integrating data types across DCCs at different stages in the life cycle of a Common Fund project is difficult post-hoc but is supported by efforts including onboarding new data types, ontologies, and standardized metadata. Analysis workflows also partially address data integration challenges by building dynamic analytic strategies. For example, the Playbook Workflow Builder enables non-computational biologists to develop modular code-free analysis workflows that draw from multiple data sources and analysis pipelines to enable reproducible, cross-DCC discovery.^21^ Workflow development is also supported through the generation of APIs by CFDE programs and centers. Visualization tools like Vitessce,^75^ the Cell Distance Explorer,^31^ the Exploration User Interface,^53^ or the HRA Organ Gallery^76^ support standard data formats and make it possible to display, explore, and download 2D and 3D data visualizations. However, they do not support all data formats, and multiscale data exploration and zoom, from the meter size human body to nm-scale biomarkers (e.g., 3D protein structures), is challenging (Figure 5).

Due to the 10-year limit of the legislation establishing Common Fund programs, many DCCs (e.g., GTEx) support ‘legacy’ datasets maintained through more limited versions of their data coordinating centers and/or deposited in data repositories but no longer supported by an underlying Consortium. Other DCCs (e.g., Metabolomics Workbench) overcome this challenge by serving as a perpetual knowledge center for particular data types. The CFDE Centers also support certain functions required for DCC sustainability. For example, the CWIC partially addresses sustainability concerns by providing computational infrastructure, lightweight long-term maintenance, multiple funding streams, and a model for data analysis that requires authentication. However, certain sustainability functions fall outside the purview of the Centers.

Finally, there are hurdles in establishing, broadening, and educating a user base for CFDE and in creating a unified experience for a wide array of user archetypes. The massive operational and domain-level scales of CFDE makes it difficult to define a single group of target users, and each CFDE DCC has its own unique consortia, which vary from one another and from other users outside of these consortia. Despite these challenges, CFDE has observed substantial increases in usage over time, as mentioned in the “Coordination and Assessment” section above. The COMPA (Communication and Outreach to Maximize Product Adoption) partnership was established to prototype market research of three test CFDE products in order to inform a CFDE-wide strategy to enhance the visibility and adoption of its range of products. The COMPA partnership studied three resources across the CFDE landscape and found the following problems across them: lack of user tracking, low awareness of CFDE resources, insufficient onboarding and training, undefined user personas, challenges in identifying and leveraging outreach and engagement venues, and concerns around sustainability. The report recommended developing a flexible toolkit of strategies and best practices in order to improve user identification, outreach and awareness, usability and support, programmatic API access, and user tracking and metrics. Additionally, new CFDE tools have been developed to identify the CFDE data or knowledge most pertinent to individual users, such as the KC tool CFDE-REVEAL (https://cfdeknowledge.org/r/cfde_reveal; see “Tools and interfaces for data analysis, visualization, and discovery”).

## Discussion

CFDE was established to directly address fundamental challenges hindering biomedical discovery across NIH Common Fund programs: siloed repositories developed by specific projects, heterogeneous formats that impede cross-study analysis, and limited sustainability of data beyond individual program lifespans. The core of CFDE’s success lies in its adoption of a hybrid federated data ecosystem model rather than a fully centralized repository or a pure runtime federated query system. This strategy avoids duplicating massive datasets by harmonizing the key elements of rich metadata while leaving the raw data in its original DCC repository, thereby respecting DCC autonomy and providing a unified discovery experience. Federation is facilitated in part by the C2M2 model, a flexible metadata schema that supports consistent search and integration across DCC resources. By integrating metadata from NIH Common Fund programs individually designed for transformational science and discovery and for re-engineering the research enterprise, CFDE provides a crucial infrastructure layer, enhancing the value of DCC resources.

Systematic cost-benefit analyses within other ecosystems outside of CFDE have shown that harmonized ontologies, tools, computational support, and training can save researchers time, streamline the research process, and lead to economic value.^77,78^ Similar future cost-benefit analyses of CFDE resources will be useful to demonstrate the impact of CFDE on these outcomes and to inform broader recommendations for other large ecosystems. Other large biomedical consortia have additionally demonstrated the value of accessible platforms for reference datasets, standards, and interoperability (e.g,, The Encyclopedia of DNA Elements (ENCODE)^79^, The Human Cell Atlas (HCA)^80^, The Monarch Consortium^81^, The International Cancer Genome Consortium^82^, All of Us^83^, and UK Biobank^84^), though these initiatives differ in their architecture (federated versus centralized) and data access models (open versus controlled).

In considering the challenge of integrative data reuse from the breadth of Common Fund programs, a single portal or commons is unlikely to be sufficient to drive transformative discoveries. CFDE is designed as an ecosystem with multiple specialized communities brought together alongside the software, skills, and data to enable success. This represents a shift in how biomedical data are managed and utilized and provides a blueprint for future efforts. By overcoming fragmentation through a rigorous and community-driven approach, the ecosystem is supporting and enhancing the value of NIH Common Fund programs, positioning them for integrative analyses that support new discoveries.

## Methods

### Preparation of CFDE Review

This review was collectively assembled by members from across the CFDE community. A small Publication Working Group with representation from DCCs, partnerships, and centers outlined the key features of CFDE to be described and assigned individual manuscript sections to community members with domain expertise in them. For a subset of sections, a Publication Writing Sprint was held at the Fall 2025 CFDE Meeting, where ∼60 members from across CFDE collectively contributed to brainstorming and drafting initial versions of the text. The text was further refined by the Publication Working Group, and then by all co-authors from across CFDE to produce the final version.

### Development of Aims for CFDE Omics Skills Development

The Training Center utilized a mixed-method approach including a review of existing CFDE trainings, key informant interviews with CFDE subject matter experts and learners, a systematic literature review, a CFDE trainer survey, a mentor survey, an analysis of CFDE datasets mentioned in academic research, and information gathering from CFDE stakeholders from Data Coordinating Centers (DCCs). Each aim includes clearly defined goals and outcomes to ensure a structured and effective learning experience. The aims are based on three core learning elements: Remember, Understand, and Apply.

### Metrics Collection

We developed software to collect metrics on resource usage on CFDE-developed software and web properties from GitHub and Google Analytics alongside publications that reference CFDE funding. For GitHub, awardees tag repositories with their grant identifier. We then use the GitHub API to find each tagged repository and gather total and over-time statistics on forks, stars, licenses, and the presence of key repository elements (README, CONTRIBUTING, and similar files). For Google Analytics, we use a similar mechanism. Awardees grant programmatic access to their existing website “properties” and tag them with grant identifiers.

We then use the Google Analytics API to list all properties we have access to, associate them with particular core projects, and gather total and over-time statistics on unique visitors, engaged interactions, most common geographic regions, and more. For publications, we gather grant-associated publications from NIH Reporter and obtain citation metrics for these publications using iCite. We present these data in a password-protected dashboard that is regularly updated, resulting in a holistic view of metrics across CFDE.

### FAIRshake Evaluation

In March 2026, average FAIRshake scores were calculated across all submitted assets (n=70) from all DCCs (n=15) over time (years 2018-2026). These included dataset assets (C2M2 data packages, knowledge graph assertions, gene set libraries, and attribute tables) and code assets (APIs, ETLs, landing pages templates such as gene, drug, and disease pages, and models). FAIRshake scores were plotted relative to the total number of assets at each time point.

### Data Distillery Knowledge Graph Query Construction

To demonstrate capabilities for cross-ecosystem discovery, we performed a query against the Data Distillery Knowledge Graph (DDKG), a biomedical knowledge graph developed from the Unified Biomedical Knowledge Graph (UBKG) and hosted through the Neo4j platform (Neo4j, Inc.; https://neo4j.com/). The DDKG incorporates multiple CFDE programs, including Metabolomics Workbench (MW), Genotype-Tissue Expression (GTEx), and Illuminating the Druggable Genome (IDG). The KG encodes provenance through a Source Abbreviation (SAB), which defines the original data source.

A Cypher query was developed to extract a mechanistically plausible subgraph. The query first uses Metabolomics Workbench annotations to generate a hypothesis for which a gene causally affects a metabolite, and the metabolite is connected with a disease. A secondary layer of constraints is then applied to require that a) the metabolite is generated within a specific tissue according to Metabolomics Workbench annotations, and b) the gene is expressed in the same tissue according to GTEx. Tissues were harmonized across data sources using UBKG-derived controlled vocabularies. Finally, the query adds upstream relationships from IDG to connect compounds to proteins and gene products, thus nominating candidate drugs for targeting the associated pathway.

~~~
//Tabular Output Query
MATCH (compound_concept:Concept)-[r1:bioactivity
{SAB:“IDGP”}]->(protein_concept:Concept)-[r2:gene_product_of
{SAB:“UNIPROTKB”}]->(gene_concept:Concept)-[r3:causally_influences
{SAB:“MW”}]->(metabolite_concept:Concept)-[r4:correlated_with_condition
{SAB:“MW”}]->(condition_concept:Concept),(metabolite_concept:Concept)-[r5:produced_by
{SAB:“MW”}]->(tissue_concept_1:Concept),(tissue_concept_2:Concept)-[r6:expresses
{SAB:“GTEXEXP”}]->(gtexexp_concept:Concept)-[r7
{SAB:“GTEXEXP”}]->(gene_concept:Concept),(gtexexp_concept:Concept)-[r8:has_expression
{SAB:“GTEXEXP”}]->(exp_concept:Concept)-[:CODE]-(exp_code:Code),(condition_concept:Concept)-[]->(t issue_concept_1:Concept),(tissue_concept_1:Concept)-[]->(tissue_concept_2:Concept) WITH *
MATCH (compound_concept:Concept)-[:PREF_TERM]-(compound:Term),
(protein_concept:Concept)-[:PREF_TERM]-(protein:Term),
(gene_concept:Concept)-[:PREF_TERM]-(gene:Term),
(condition_concept:Concept)-[:PREF_TERM]-(condition:Term),
(metabolite_concept:Concept)-[:PREF_TERM]-(metabolite:Term),
(tissue_concept_1:Concept)-[:PREF_TERM]-(tissue_1:Term),
(tissue_concept_2:Concept)-[:PREF_TERM]-(tissue_2:Term) RETURN DISTINCT
compound.name,protein.name,gene.name,metabolite.name,tissue_1.name,tissue_2.name,condition.name,ex p_code.CODE limit 10
~~~

**Query Description:** we formulate a comprehensive Cypher query that conducts a dense, multi-branch graph traversal over biomedical `Concept` nodes by first establishing a primary causal chain in which a `compound_concept` is linked to a `protein_concept` via a `bioactivity` relationship constrained by IDG data through `SAB:“IDGP”`, the protein is connected to a `gene_concept` through `gene_product_of` with `SAB:“UNIPROTKB”`, the gene is propagated downstream to a

`metabolite_concept` via `causally_influences` (`SAB:“MW”`), and the Metabolomics Workbench metabolite is further associated with a `condition_concept` through `correlated_with_condition` (`SAB:“MW”`), thereby encoding a compound-to-phenotype mechanistic pathway. In parallel within the same `MATCH` clause, we populate this backbone by linking the `metabolite_concept` to a first tissue context `tissue_concept_1` using `produced_by` (`SAB:“MW”`), while independently introducing a second tissue context `tissue_concept_2` that `expresses` a `gtexexp_concept` (`SAB:“GTEXEXP`”), which is then connected via an additional GTEx-associated relationship (also filtered by `SAB:“GTEXEXP`”, though without an explicit type) back to the same `gene_concept`, ensuring that the gene in the causal chain is the one whose expression is being measured. We further traverse from the `gtexexp_concept` to an `exp_concept` through `has_expression` (`SAB:“GTEXEXP`”) and add an expression identifier via a `:CODE` relationship to an `exp_code:Code` node, capturing the measurable expression output. To maintain biological coherence across subgraphs, we also impose relationship constraints requiring that the `condition_concept` connects to `tissue_concept_1` and that `tissue_concept_1` connects to `tissue_concept_2`, thereby implicitly aligning disease relevance, metabolite production, and gene expression across anatomically or functionally related tissues. After this extensive pattern matching, we propagate all bound variables forward using `WITH *`, preserving the full intermediate result set for subsequent transformation.

We then conduct a second `MATCH` clause that systematically resolves each `Concept` node (`compound_concept`, `protein_concept`, `gene_concept`, `condition_concept`, `metabolite_concept`, `tissue_concept_1`, `tissue_concept_2`) to its corresponding human-readable `Term` node via `PREF_TERM` relationships, effectively translating ontology-level identifiers into displayable names. Finally, we project a deduplicated (`RETURN DISTINCT`) tabular output consisting of `compound.name`, `protein.name`, `gene.name`, `metabolite.name`, `tissue_1.name`, `tissue_2.name`, `condition.name`, and the associated `exp_code.CODE`, and constrain the result cardinality with `LIMIT 10`, yielding a compact yet semantically rich integration of pharmacological activity, molecular biology, metabolic influence, disease association, tissue specificity, and transcriptomic expression measurements derived from multiple curated data sources.

### Visualization of Multi-Source Datasets Registered in the HuBMAP Human Reference Atlas

A list of all CFDE and non-CFDE datasets registered in the HuBMAP Human Reference Atlas was obtained. These included datasets integrated through 3D registration of tissue into the CCF or through crosswalking cell type annotations via HRA-compatible transcriptomic tools (e.g., Azimuth) or spatial omics tools (e.g., DeepCellTypes). The top sources of these datasets were plotted against 30 anatomical structures from which they were derived, and the number of datasets and total cell counts within each was plotted (Figure 6d). Analyses and visualizations were generated using reproducible Python scripts accessible through a public GitHub repository (https://github.com/cns-iu/hra-cfde-marker-visualizations/blob/main/hra-pop-scatter-checkpoint.ipynb). Dataset-level metadata were mapped to standardized anatomical structures defined by the Human Reference Atlas to enable cross-consortium comparisons of organ coverage.

## Resource Availability

### Lead contact

Requests for further information and resources should be directed to and will be fulfilled by the lead contact, Julie Jurgens.

### Materials availability

This study did not generate new unique reagents.

### Data and code availability

This paper analyzes existing, publicly available data. Common Fund programs’ metadata and processed datasets are available through https://data.cfde.cloud/. CFDE has developed a host of Standards, which are available at https://github.com/nih-cfde/rfcs/blob/master/adoptionstatus.md, and additional user guides, protocols, and standards available through https://data.cfde.cloud/documentation.

Data and original code for the HuBMAP analysis within this manuscript have been deposited at GitHub and are publicly available at https://github.com/cns-iu/hra-cfde-marker-visualizations/blob/main/hra-pop-scatter-checkpoint.ipynb as of the date of publication.

## Acknowledgments

Dedicated to the memory of Philip Bourne (1953-2026), first Associate Director for Data Science at the National Institutes of Health, whose efforts were critical to the development of CFDE. Article processing fees were supported by the CFDE Knowledge Center (OT2OD036440). This work was supported in part by funds from the NIH Office of the Director / NIH Common Fund (grants OT2OD036440, U54OD036472, OT2OD036435, OT2OD037922, OT2OD037936, OT2OD030160, OT2OD030162, OT2OD030544, OT2OD032619, OT2OD030541, OT2OD030546, U24OD036598, U24OD038425, U24OD038422, U24OD038421, U24OD038424, U24OD038423, OT2OD032119, U24OD026629, OT2OD030161, OT2OD032092, OT2OD030547, OT2OD032720, OT2OD032701, OT2OD032644, OT2OD032742, UM1OD023221, OT2OD030545, OT2OD026671, OT2OD033756, OT2OD030545, 1R03OD039970-01), the National Institute of General Medical Sciences (1R24GM146616, R35GM141873), the National Institute on Drug Abuse (3U54DA049098, 1U54DA036134-01, 1U54DA049098-01, 1U54DA049098-01S1, UM1DA058230, U54DA049110, U54DA049116, U54DA049115, U54DA049113), the National Human Genome Research Institute (U24HG010423, U24HG007822, U54HG012510, U54HG012513, U54HG012517, U54HG0006364, U01HG004080, U24HG006620), the National Heart Lung and Blood Institute (U24HL168712, K23HL164980), the National Institute of Neurological Disorders and Stroke (UM1NS118922, UM1NS112874, U24NS112873), the National Center for Research Resources (U42RR024244), the National Cancer Institute (U24CA268108), the National Institute on Aging (U54AG076043), NIH Office of the Director (U24OD038423, 1U24OD038422, 1R03OD030600), the National Institute of Diabetes and Digestive and Kidney Diseases (U01DK133090, U2CDK114886, U24DK135157, U2CDK119886, U24DK141185), and the American Heart Association (23CDA1040581). K.B. is a co-director of and funded by the MacMillan Multiscale Human program by the Canadian Institute for Advanced Research (CIFAR). K.B. is also supported via a Stiftung Charité Visiting Fellowship via Berlin Institute of Health at Charité (BIH). A.B. was supported by OT2OD026675 and OT2OD033759 under the NIH JumpStart Fellowship and by a supplement award on U54AG076043. S.K.T. is a Scholar of Blood Cancer United, holds the Joshua Kahan Endowed Chair in Pediatric Leukemia Research at the Children’s Hospital of Philadelphia, and is/was supported by the NIH/NCI (1U01CA232486, 1U01CA243072, 1R01CA293587) and a Pennsylvania Department of Health Commonwealth Universal Research Enhancement award. P.P. acknowledges Startup Funds from the College of Medicine at Phoenix, University of Arizona. We acknowledge the U.S. National Institutes of Health (NIH) Common Fund, all CFDE participating Data Coordinating Centers, the Somatic Cell Genome Editing (SCGE) Consortium, and the dedicated teams across CFDE.

## Author contributions

Conceptualization, J.A.J., A.B., T.M.A., B.S., N.B., J.P.C., J.C., P.P., S.D., D.M.T., K.B. (Katy Börner), A.D., K.B. (Kelli Bursey), A.M. (Avi Ma’ayan), R.M., M.E.R., C.S.G.; Data Curation, B.J.H.; Formal analysis, A.B., T.M.A., B.S., J.B. (Jennifer Burnette), A.D., K.B. (Kelli Bursey), K.B. (Katy Börner), and C.S.G.; Funding acquisition, A.B., N.B., J.P.C., L.G., J.C., P.P., S.D., D.M.T., K.B. (Katy Börner), A.M. (Avi Ma’ayan), R.M., M.E.R., M.T.W., C.H.W, T.I.O., and C.S.G.; Investigation, A.B., T.M.A., B.S., E.Z., D.W., C.R., J.B. (Jennifer Burnette), L.G., A.D., K.B. (Kelli Bursey), P.P., D.M.T., K.B. (Katy Börner), S.D., and C.S.G.; Methodology, A.B., T.M.A., B.S., E.Z., D.W., C.R., J.B. (Jennifer Burnette), L.G., A.D., K.B. (Kelli Bursey), P.P., D.M.T., K.B. (Katy Börner), C.S.G., M.A., C.C., K.E.R, C.H.W.; Project administration, J.A.J., R.M., M.E.R., C.S.G; Software, A.B., T.M.A., B.S., K.B. (Katy Börner), J.A.S.; Supervision, J.A.J., R.M., M.E.R., C.S.G. C.C, M.A; Visualization, J.A.J., A.B., M.R.M., T.M.A., B.S., S.L.J., E.Z., D.W., C.R., M.B., A.S., P.P., D.M.T., K.B. (Katy Börner), A.D., and K.B. (Kelli Bursey); Writing – original draft, J.A.J., A.B., J.V., M.R.M., T.M., E.Z., B.S., D.W., C.R., S.R., A.N., N.C., M.B., S.T., D.H.K., M.C.M., I.D., H.E.C., J.A.S., S.K.T., J.E.E., H.D., S.S. (Surya Saha), M.A.L., V.G., C.N., J.Z., K.E.R., A.I.B., A.S., V.T.M., S.D., C.G.B., S.S. (Sumana Srinivasan), D.J. (Dongkeun Jang), P.K., L.D.T., M.P.L., V.P., M.A., M.K., D.J.C. (Daniel J. B. Clarke), A.I., D.J.C. (Daniel J. Crichton), D.B., C.C., A.J.S., A.M. (Ashish Mahabal), I.C., S.K., D.M., K.Y., K.J.G., D.J-M (David Jimenez-Morales), J.R., B.J.H., W.W., C.H.W., A.M. (Aleksandar Milosavljevic), P.D.B., J.B. (Jyl Boline), T.I.O., C.G.L., B.d., P.J.P., J.C.S., J.F., J.J.Y., J.S.G., S.S. (Shankar Subramaniam), M.T., T.C., M.T.W., A.K., J.B. (Jennifer Burnette), R.R., M.C.S., L.G., N.P.B., J.P.C., J.C., P.P., D.M.T., K.B. (Katy Börner), A.D., K.B. (Kelli Bursey), A.M. (Avi Ma’ayan), R.M., M.E.R., and C.S.G.; Writing –review & editing, J.A.J., A.B., J.V., M.R.M., T.M., E.Z., B.S., S.D., D.W., C.R., S.R., A.N., N.C., M.B., S.T., D.H.K., M.C.M., I.D., H.E.C., J.A.S., S.K.T., S.L.J., J.E.E., H.D., S.S. (Surya Saha), M.A.L., V.G., C.N., J.Z., K.E.R., A.I.B., A.S., V.T.M., C.G.B., S.S. (Sumana Srinivasan), D.J. (Dongkeun Jang), P.K., L.D.T., M.P.L., V.P., M.A., M.K., D.J.C. (Daniel J. B. Clarke), A.I., D.J.C. (Daniel J. Crichton), S.S. (Shava Smallen), D.B., C.C., A.J.S., A.M. (Ashish Mahabal), I.C., S.K., D.M., K.Y., K.J.G., D.J-M (David Jimenez-Morales), J.R., B.J.H., W.W., C.H.W., A.M. (Aleksandar Milosavljevic), P.D.B., J.B. (Jyl Boline), T.I.O., C.G.L., B.d., P.J.P., J.C.S., J.F., J.J.Y., J.S.G., S.S. (Shankar Subramaniam), M.T., T.C., M.T.W., A.K., J.B. (Jennifer Burnette), R.R., M.C.S., L.G., N.P.B., J.P.C., J.C., P.P., D.M.T., K.B. (Katy Börner), A.D., K.B. (Kelli Bursey), A.M. (Avi Ma’ayan), R.M., M.E.R., C.S.G., T.C. (Timothy Clark) and B.J.H.

## Declarations of interest

S.K.T. receives/d research funding from Incyte Corporation and Kura Oncology, serves/d on scientific advisory boards for Aleta Biotherapeutics, Amgen, AstraZeneca, Jazz Pharmaceuticals, Kura Oncology, and Syndax Pharmaceuticals, and received travel support from Amgen (all for unrelated studies). M.T.W. receives/d research funding from Astra Zeneca, Alexion, Bristol Myers Squibb, Salubris Bio, and Nihon Kohden, all outside the submitted work. T.I.O. is an advisor to InSilico Medicine, Chair of the Scientific Advisory Board for ExScalate, Dompé farmaceutici SpA, and CEO of Expert Systems, Inc. The remaining authors declare no competing interests related to this study.

## Consortia

The members of the CFDE Consortium are Enis Afgan, Nasheath Ahmed, Jain Aluvathingal, Eugenia Ampofo, Lisa Anderson, Jessie E. Arce, Kristin Ardlie, Sena Arpinar, Euan Ashley, Ahmed Awan, Dannon Baker, Clara Bakker, Alexander Barnes, Michele Berselli, Lynette Bower, Alan Bradley, Arthur Brady, C. Titus Brown, Laurel Brown, Miguel Brown, David Burns, Saranya Canchi, Robert Carter, Amanda L. Charbonneau, Kyle Chard, David Chen, Jing Chen, John Chilton, Asif T. Chinwalla, Sonal Choudhary, Christopher Churas, David Clary, Kevin Coakley, Kristina A. Cole, Tyler Collins, Nate Coraor, Sierra Corban, Ryan J. Corbett, Jeremy Costanza, Maria Costanzo, Jonathan Crabtree, Heather H. Creasy, Gilbert Curbelo III, Karl Czajkowski, Mike D’Arcy, Maytal Dahan, Vlado Dancik, John Davis, Pieter de Jong, Eden Z. Deng, Jack DiGiovanna, Sharon J. Diskin, Leslie Duffy, Eoin Fahy, Tierra Farris, Victor Felix, Erik Ferlanti, Vincent Ferretti, John Fonner, Ian Foster, Robert Fullem, Steven Gage, Vikram Adithya Ganesh, Aydan Gasimova, Michelle G. Giglio, Alicia Gingrich, Aysam Guerler, Yiran Guo, Shakti Gupta, Jan N. Hansen, Andrew Hardy, Rayna M. Harris, Allison P. Heath, David M. Higgins, Dave Durell Hill, Quy Hoang, Theresa K. Hodges, Paul Hoover, Michelle R. Hribar, Olukemi Ifeonu, Andrew Jackson, Minji Jeon, Bosko Jevtic, Dubravka Jevtic, Ethan Ji, Robel Kahsay, Kate Kaya, Carl Kesselman, Christine Kirkpatrick, Ryan Koesterer, Diane Krause, Eryk Kropiwnicki, Natalie Kucher, Parul Kudtarkar, Sujeet Kulkarni, Alexander Lachmann, Aditya Lahiri, Ethan M. Lange, Louise Lanoue, Aaron Y. Lee, Cecilia S. Lee, Yuna Lee, Qi Li, Marisa C.W. Lim, R. Lee Liming, K. C. Kent Lloyd, Jessica Lumian, Emma Lundberg, Anup A. Mahurkar, Meisha Mandal, Giacomo B. Marino, Julia Markowski, Maryann E. Martone, Michele Mattioni, David A. Mayer, Colin McKerlie, Susan L. McRitchie, Nathaniel Mendoza, Vishal Midya, Aleksandar Mihajlovic, Vuk Milinovic, Daniel Miller, Richard S. Morgan, Stephen Mosher, Juan Muerto, James B. Munro, Kay Métis, Stephen B. Montgomery, Suvarna Nadendla, Sara Narayanaswamy, Rahi Navelkar, Jared Nedzel, Thanh Long Nguyen, Trang Nguyen, Lauryl Nutter, Lucila Ohno-Machado, Joseph Okonda, Stephanie Olaiya, Bhavesh Patel, Wimal Pathmasiri, Lauren Petrick, Hedda Porchaska, Melody Porterfield, Abinanda Prabhakaran, Sara Rahiminejad, Pritham Ram, Shashidhar Rao, Regina Renfro, Adam Resnick, Jessica Respicio, Rudyard Richter, Philippe Rocca-Serra, Jo Lynne Rokita, Cia Romano, William Ronchetti, Alexander J. Ropelewski, Jared Rozowsky, Vincent Rubinetti, Blake R. Rushing, Kristi Sadowski, Ethan Sanchez, Susanna-Assunta Sansone, Michelle Savage, Michael Schor, Robert E. Schuler, Zhandos Sembay, Bill Shirey, Max Sibilla, Alex Sickler, William Skarnes, Patrick Smadbeck, Neela Srinivasan, Shivaramakrishna Srinivasan, Matt Stelmaszek, Desmond Stubbs, Joe Stubbs, Keith Suderman, Patricia J. Sullivan, Susan J. Sumner, Ying Sun, Hongsuda Tangmunarunkit, DeBran Tarver, Deanne Taylor, Susan L. Teitelbaum, Jean-Philippe Thibert, Casey Thomas, Luca Tucciarone, Joshua Urrutia, Alexander D. Veit, Jennifer Vessio, Rick Wagner, Alex Waldrop, Jeremy Walter, Eric Wenger, Natalie Whitaker-Allen, Owen White, Cris Williams, Karen Word, Zhuorui Xie, Chase Yakaboski, Zhou Yuan, Shiping Zhang, Yuanchao Zhang, Haoquan Zhao, Yuankun Zhu, and Felix Zuo.

## Notes

### Summary of Updates

Minor updates in wording were made to the manuscript body and program description in Table 1.

## References

1. Collins, F.S., Wilder, E.L., and Zerhouni, E. (2014). Funding transdisciplinary research. NIH Roadmap/Common Fund at 10 years. Science 345, 274–276.

2. Zerhouni, E. (2003). Medicine. The NIH Roadmap. Science 302, 63–72.

3. Piwowar, H.A., and Vision, T.J. (2013). Data reuse and the open data citation advantage. PeerJ 1, e175.

4. Manzoni, C., Kia, D.A., Vandrovcova, J., Hardy, J., Wood, N.W., Lewis, P.A., and Ferrari, R. (2018). Genome, transcriptome and proteome: the rise of omics data and their integration in biomedical sciences. Brief Bioinform 19, 286–302.

5. Shome, M., MacKenzie, T.M.G., Subbareddy, S.R., and Snyder, M.P. (2024). The Importance, Challenges, and Possible Solutions for Sharing Proteomics Data While Safeguarding Individuals’ Privacy. Mol Cell Proteomics 23, 100731.

6. Sharing and re-using open data: A case study of motivations in astrophysics (2019). International Journal of Information Management 49, 228–241.

7. Charbonneau, A.L., Brady, A., Czajkowski, K., Aluvathingal, J., Canchi, S., Carter, R., Chard, K., Clarke, D.J.B., Crabtree, J., Creasy, H.H., et al. (2022). Making Common Fund data more findable: catalyzing a data ecosystem. Gigascience 11. 10.1093/gigascience/giac105.

8. Evangelista, J.E., Clarke, D.J.B., Byrd, A.I., Srinivasan, S., Srinivasan, S., Maurya, M.R., Jenkins, S.L., Diamant, I., Sanchez, E., Xie, Z., et al. (2026). The CFDE Workbench: Integrating Metadata and Processed Data from Common Fund Programs. J Mol Biol, 169631.

9. Ye, Y., Barapatre, S., Davis, M.K., Elliston, K.O., Davatzikos, C., Fedorov, A., Fillion-Robin, J.-C., Foster, I., Gilbertson, J.R., Lasso, A., et al. (2021). Open-source Software Sustainability Models: Initial White Paper From the Informatics Technology for Cancer Research Sustainability and Industry Partnership Working Group. J Med Internet Res 23, e20028.

10. Chandras, C., Weaver, T., Zouberakis, M., Smedley, D., Schughart, K., Rosenthal, N., Hancock, J.M., Kollias, G., Schofield, P.N., and Aidinis, V. (2009). Models for financial sustainability of biological databases and resources. Database (Oxford) 2009, bap017.

11. Wilcox, A., Randhawa, G., Embi, P., Cao, H., and Kuperman, G.J. (2014). Sustainability considerations for health research and analytic data infrastructures. EGEMS (Wash DC) 2, 1113.

12. Hinkson, I.V., Davidsen, T.M., Klemm, J.D., Kerlavage, A.R., Kibbe, W.A., and Chandramouliswaran, I. (2017). A Comprehensive Infrastructure for Big Data in Cancer Research: Accelerating Cancer Research and Precision Medicine. Front Cell Dev Biol 5, 83.

13. Evangelista, J.E., Clarke, D.J.B., Xie, Z., Marino, G.B., Utti, V., Jenkins, S.L., Ahooyi, T.M., Bologa, C.G., Yang, J.J., Binder, J.L., et al. (2023). Toxicology knowledge graph for structural birth defects. Commun Med (Lond) 3, 98.

14. Ahooyi, T.M., Stear, B., Alan Simmons, J., Metzger, V.T., Kumar, P., Evangelista, J.E., Clarke, D.J.B., Xie, Z., Kim, H., Jenkins, S.L., et al. (2025). The Data Distillery: A Graph Framework for Semantic Integration and Querying of Biomedical Data. bioRxiv, 2025.08.11.666099. 10.1101/2025.08.11.666099.

15. Yu, K., Zhao, H., Wilderman, A.S., Farris, T.R., Arce, J.E., Chen, D., Jackson, A.R., Guo, Y., Li, Q., Jevtic, B., et al. (2026). Aggregation of gene regulatory information and knowledge on FAIR principles enables discovery of pathogenic gene regulatory variants. Bioinformatics 42. 10.1093/bioinformatics/btag013.

16. Akhtar, M., Benjelloun, O., Conforti, C., Foschini, L., Giner-Miguelez, J., Gijsbers, P., Goswami, S., Jain, N., Karamousadakis, M., Kuchnik, M., et al. (2024). Croissant: A Metadata Format for ML-Ready Datasets. 10.1145/3650203.3663326.

17. Jacobsen, A., de Miranda Azevedo, R., Juty, N., Batista, D., Coles, S., Cornet, R., Courtot, M., Crosas, M., Dumontier, M., Evelo, C.T., et al. (2024). FAIR Principles: Interpretations and Implementation Considerations. Data Intelligence 2, 10–29.

18. Common Fund Data Ecosystem Working Group https://dpcpsi.nih.gov/council/working-groups/cfde.

19. Masood, D., Kim, M., Vora, J., Kahsay, R., McNeeley, P., Kim, S., Kulkarni, S., Natale, D.A., Ramachandran, S., Gupta, S., et al. (2026). BiomarkerKB: An Integrated Knowledgebase Supporting Biomarker-Centric Exploration of Biomedical Data. bioRxiv, 2026.01.26.701395. 10.64898/2026.01.26.701395.

20. Stear, B.J., Mohseni Ahooyi, T., Simmons, J.A., Kollar, C., Hartman, L., Beigel, K., Lahiri, A., Vasisht, S., Callahan, T.J., Nemarich, C.M., et al. (2024). Petagraph: A large-scale unifying knowledge graph framework for integrating biomolecular and biomedical data. Sci Data 11, 1338.

21. Clarke, D.J.B., Evangelista, J.E., Xie, Z., Marino, G.B., Byrd, A.I., Maurya, M.R., Srinivasan, S., Yu, K., Petrosyan, V., Roth, M.E., et al. (2025). Playbook workflow builder: Interactive construction of bioinformatics workflows. PLoS Comput Biol 21, e1012901.

22. Higgins, D., Thibert, J.-P., Mattioni, M., DiGiovanna, J., Grossman, R.L., Farrow, B.K., Wenger, E., Volchenboum, S., Carroll, R.J., Haendel, M.A., et al. (2023). Abstract 6576: Gabriella Miller Kids First Data Resource Center (KFDRC): Empowering discovery across germline and somatic variation in pediatric cancer. Cancer Res. 83, 6576–6576.

23. Dekker, J., Belmont, A.S., Guttman, M., Leshyk, V.O., Lis, J.T., Lomvardas, S., Mirny, L.A., O’Shea, C.C., Park, P.J., Ren, B., et al. (2017). The 4D nucleome project. Nature 549, 219–226.

24. Reiff, S.B., Schroeder, A.J., Kırlı, K., Cosolo, A., Bakker, C., Mercado, L., Lee, S., Veit, A.D., Balashov, A.K., Vitzthum, C., et al. (2022). The 4D Nucleome Data Portal as a resource for searching and visualizing curated nucleomics data. Nat. Commun. 13, 2365.

25. Clark, T., Mohan, J., Schaffer, L., Obernier, K., Al Manir, S., Churas, C.P., Dailamy, A., Doctor, Y., Forget, A., Hansen, J.N., et al. (2024). Cell Maps for Artificial Intelligence: AI-Ready Maps of Human Cell Architecture from Disease-Relevant Cell Lines. bioRxiv. 10.1101/2024.05.21.589311.

26. Sud, M., Fahy, E., Cotter, D., Azam, K., Vadivelu, I., Burant, C., Edison, A., Fiehn, O., Higashi, R., Nair, K.S., et al. (2016). Metabolomics Workbench: An international repository for metabolomics data and metadata, metabolite standards, protocols, tutorials and training, and analysis tools. Nucleic Acids Res 44, D463–D470.

27. Sluka, K.A., Wager, T.D., Sutherland, S.P., Labosky, P.A., Balach, T., Bayman, E.O., Berardi, G., Brummett, C.M., Burns, J., Buvanendran, A., et al. (2023). Predicting chronic postsurgical pain: current evidence and a novel program to develop predictive biomarker signatures. Pain 164, 1912–1926.

28. Urrutia, J., Vaughn, M.W., Scherer, P., Taub, M.A., Ansari, B., Hackman, A., Gherman, A., Garcia, C., Stubbs, J., Carson, J.P., Lindquist, M.A., Kahn, A.B. The Virtual Biospecimen Repository. In Submitted to The Proceedings of the Practice and Experience in Advanced Research Computing Conference (PEARC 2026).

29. Börner, K., Blood, P.D., Silverstein, J.C., Ruffalo, M., Satija, R., Teichmann, S.A., Pryhuber, G.J., Misra, R.S., Purkerson, J.M., Fan, J., et al. (2025). Human BioMolecular Atlas Program (HuBMAP): 3D Human Reference Atlas construction and usage. Nat Methods 22, 845–860.

30. Mungall, C.J., Torniai, C., Gkoutos, G.V., Lewis, S.E., and Haendel, M.A. (2012). Uberon, an integrative multi-species anatomy ontology. Genome Biol 13, R5.

31. Jain, Y., Jepson, J., Chen, R., Maier, E., Herr, B.W., Puig-Barbe, A., Quardokus, E.M., Qaurooni, D., Yapp, C., Ewing, S.L., et al. (2025). Exploring endothelial cell environments across organs in spatially resolved omics data. bioRxiv, 2025.09.23.678129. 10.1101/2025.09.23.678129.

32. Bueckle, A., Herr, B.W., Chen, L., Bolin, D., Qaurooni, D., Ginda, M., Jain, Y., Puig-Barbe, A., Ardlie, K., Wang, F., et al. (2025). Cell type populations for 3D anatomical structures of the Human Reference Atlas. bioRxiv, 2025.08.14.670406. 10.1101/2025.08.14.670406.

33. Clark, T., Caufield, H., Parker, J.A., Al Manir, S., Amorim, E., Eddy, J., Gim, N., Gow, B., Goar, W., Haendel, M., et al. (2024). AI-readiness for Biomedical Data: Bridge2AI Recommendations. bioRxiv. 10.1101/2024.10.23.619844.

34. International Organization for Standardization (2004). ISO/IEC 11179-1: information technology - metadata registries (MDR) - part 1: framework.

35. 35. Data Catalog Vocabulary (DCAT) - Version 3 https://www.w3.org/TR/2024/REC-vocab-dcat-3-20240822/.

36. Bueckle, A., Herr, B.W., 2nd, Hardi, J., Quardokus, E.M., Musen, M.A., and Börner, K. (2025). Construction, Deployment, and Usage of the Human Reference Atlas Knowledge Graph. Sci Data 12, 1100.

37. Evangelista, J.E., Lutsky, A.D., Byrd, A.I., Clarke, D.J.B., Prabhakaran, A., Jenkins, S.L., and Ma’ayan, A. (2025). Creating an Interactive Web Interface for Networks Stored in Knowledge Graph Databases. Curr Protoc 5, e70200.

38. Kumar, P., Metzger, V.T., Purushotham, S.T., Kedia, P., Bologa, C.G., Lambert, C.G., and Yang, J.J. (2025). KG2ML: integrating knowledge graphs and positive unlabeled learning for identifying disease-associated genes. Front Bioinform 5, 1727953.

39. UniProt Consortium (2025). UniProt: the Universal Protein Knowledgebase in 2025. Nucleic Acids Res 53, D609–D617.

40. Austin, C.P., Battey, J.F., Bradley, A., Bucan, M., Capecchi, M., Collins, F.S., Dove, W.F., Duyk, G., Dymecki, S., Eppig, J.T., et al. (2004). The knockout mouse project. Nat Genet 36, 921–924.

41. Meehan, T.F., Conte, N., West, D.B., Jacobsen, J.O., Mason, J., Warren, J., Chen, C.-K., Tudose, I., Relac, M., Matthews, P., et al. (2017). Disease model discovery from 3,328 gene knockouts by The International Mouse Phenotyping Consortium. Nat Genet 49, 1231–1238.

42. Anandakrishnan, M., Ross, K.E., Chen, C., Vijay-Shanker, K., and Wu, C.H. (2025). KSMoFinder - Knowledge graph embedding of proteins and motifs for predicting kinases of human phosphosites. bioRxiv, 2025.10.21.683733. 10.1101/2025.10.21.683733.

43. Sanford, J.A., Nogiec, C.D., Lindholm, M.E., Adkins, J.N., Amar, D., Dasari, S., Drugan, J.K., Fernández, F.M., Radom-Aizik, S., Schenk, S., et al. (2020). Molecular Transducers of Physical Activity Consortium (MoTrPAC): Mapping the Dynamic Responses to Exercise. Cell 181, 1464–1474.

44. MoTrPAC Study Group, Lead Analysts, and MoTrPAC Study Group (2024). Temporal dynamics of the multi-omic response to endurance exercise training. Nature 629, 174–183.

45. Viet, S.M., Falman, J.C., Merrill, L.S., Faustman, E.M., Savitz, D.A., Mervish, N., Barr, D.B., Peterson, L.A., Wright, R., Balshaw, D., et al. (2021). Human Health Exposure Analysis Resource (HHEAR): A model for incorporating the exposome into health studies. Int J Hyg Environ Health 235, 113768.

46. Lee, B.Y., Ordovás, J.M., Parks, E.J., Anderson, C.A.M., Barabási, A.-L., Clinton, S.K., de la Haye, K., Duffy, V.B., Franks, P.W., Ginexi, E.M., et al. (2022). Research gaps and opportunities in precision nutrition: an NIH workshop report. Am J Clin Nutr 116, 1877–1900.

47. Simonyan, V., Goecks, J., and Mazumder, R. (2017). Biocompute Objects-A Step towards Evaluation and Validation of Biomedical Scientific Computations. PDA J Pharm Sci Technol 71, 136–146.

48. IEEE 2791-2020 IEEE Standards Association. https://standards.ieee.org/.

49. Kahsay, R., Vora, J., Navelkar, R., Mousavi, R., Fochtman, B.C., Holmes, X., Pattabiraman, N., Ranzinger, R., Mahadik, R., Williamson, T., et al. (2020). GlyGen data model and processing workflow. Bioinformatics 36, 3941–3943.

50. Bandrowski, A., Grethe, J.S., Pilko, A., Gillespie, T., Pine, G., Patel, B., Surles-Zeigler, M., and Martone, M.E. (2021). SPARC Data Structure: Rationale and Design of a FAIR Standard for Biomedical Research Data. bioRxiv, 2021.02.10.430563. 10.1101/2021.02.10.430563.

51. Murillo, O.D., Thistlethwaite, W., Rozowsky, J., Subramanian, S.L., Lucero, R., Shah, N., Jackson, A.R., Srinivasan, S., Chung, A., Laurent, C.D., et al. (2019). exRNA Atlas Analysis Reveals Distinct Extracellular RNA Cargo Types and Their Carriers Present across Human Biofluids. Cell 177, 463–477.e15.

52. LaPlante, E.L., Stürchler, A., Fullem, R., Chen, D., Starner, A.C., Esquivel, E., Alsop, E., Jackson, A.R., Ghiran, I., Pereira, G., et al. (2023). exRNA-eCLIP intersection analysis reveals a map of extracellular RNA binding proteins and associated RNAs across major human biofluids and carriers. Cell Genom 3, 100303.

53. Börner, K., Bueckle, A., Herr, B.W., 2nd, Cross, L.E., Quardokus, E.M., Record, E.G., Ju, Y., Silverstein, J.C., Browne, K.M., Jain, S., et al. (2022). Tissue registration and exploration user interfaces in support of a human reference atlas. Commun Biol 5, 1369.

54. Marino, G.B., Olaiya, S., Evangelista, J.E., Clarke, D.J.B., and Ma’ayan, A. (2025). GeneSetCart: assembling, augmenting, combining, visualizing, and analyzing gene sets. Gigascience 14. 10.1093/gigascience/giaf025.

55. Clarke, D.J.B., Kuleshov, M.V., Xie, Z., Evangelista, J.E., Meyers, M.R., Kropiwnicki, E., Jenkins, S.L., and Ma’ayan, A. (2022). Gene and drug landing page aggregator. Bioinform Adv 2, vbac013.

56. Clarke, D.J.B., Marino, G.B., Deng, E.Z., Xie, Z., Evangelista, J.E., and Ma’ayan, A. (2024). Rummagene: massive mining of gene sets from supporting materials of biomedical research publications. Commun. Biol. 7, 482.

57. Marino, G.B., Clarke, D.J.B., Lachmann, A., Deng, E.Z., and Ma’ayan, A. (2024). RummaGEO: Automatic mining of human and mouse gene sets from GEO. Patterns (N. Y.) 5, 101072.

58. Clarke, D.J.B., Marino, G.B., and Ma’ayan, A. (2025). A Gene Set Foundation Model Pre-Trained on a Massive Collection of Diverse Gene Sets. bioRxiv. 10.1101/2025.05.30.657124.

59. Galaxy Community (2024). The Galaxy platform for accessible, reproducible, and collaborative data analyses: 2024 update. Nucleic Acids Res 52, W83–W94.

60. Stubbs, J., Cardone, R., Packard, M., Jamthe, A., Padhy, S., Terry, S., Looney, J., Meiring, J., Black, S., Dahan, M., et al. (2021). Tapis: An API Platform for Reproducible, Distributed Computational Research. Advances in Information and Communication, 878–900.

61. Rehm, H.L., Page, A.J.H., Smith, L., Adams, J.B., Alterovitz, G., Babb, L.J., Barkley, M.P., Baudis, M., Beauvais, M.J.S., Beck, T., et al. (2021). GA4GH: International policies and standards for data sharing across genomic research and healthcare. Cell Genom 1. 10.1016/j.xgen.2021.100029.

62. Schatz, M.C., Philippakis, A.A., Afgan, E., Banks, E., Carey, V.J., Carroll, R.J., Culotti, A., Ellrott, K., Goecks, J., Grossman, R.L., et al. (2022). Inverting the model of genomics data sharing with the NHGRI Genomic Data Science Analysis, Visualization, and Informatics Lab-space. Cell Genom 2. 10.1016/j.xgen.2021.100085.

63. Ahalt, S., Avillach, P., Boyles, R., Bradford, K., Cox, S., Davis-Dusenbery, B., Grossman, R.L., Krishnamurthy, A., Manning, A., Paten, B., et al. (2023). Building a collaborative cloud platform to accelerate heart, lung, blood, and sleep research. J Am Med Inform Assoc 30, 1293–1300.

64. Ainsztein, A.M., Brooks, P.J., Dugan, V.G., Ganguly, A., Guo, M., Howcroft, T.K., Kelley, C.A., Kuo, L.S., Labosky, P.A., Lenzi, R., et al. (2015). The NIH Extracellular RNA Communication Consortium. J. Extracell. Vesicles 4, 27493.

65. Das, S., Extracellular RNA Communication Consortium, Ansel, K.M., Bitzer, M., Breakefield, X.O., Charest, A., Galas, D.J., Gerstein, M.B., Gupta, M., Milosavljevic, A., et al. (2019). The extracellular RNA Communication Consortium: Establishing foundational knowledge and technologies for extracellular RNA research. Cell 177, 231–242.

66. Mateescu, B., Jones, J.C., Alexander, R.P., Alsop, E., An, J.Y., Asghari, M., Boomgarden, A., Bouchareychas, L., Cayota, A., Chang, H.-C., et al. (2022). Phase 2 of extracellular RNA communication consortium charts next-generation approaches for extracellular RNA research. iScience 25, 104653.

67. Amolegbe, S.M., Johnston, N.C., Ambrosi, A., Ganguly, A., Howcroft, T.K., Kuo, L.S., Labosky, P.A., Rudnicki, D.D., Satterlee, J.S., Tagle, D.A., et al. (2025). Extracellular RNA communication: A decade of NIH common fund support illuminates exRNA biology. J Extracell Vesicles 14, e70016.

68. Sankar, B.S., Gilliland, D., Rincon, J., Hermjakob, H., Yan, Y., Adam, I., Lemaster, G., Wang, D., Watson, K., Bui, A., et al. (2024). Building an Ethical and Trustworthy Biomedical AI Ecosystem for the Translational and Clinical Integration of Foundation Models. Bioengineering (Basel) 11. 10.3390/bioengineering11100984.

69. Clarke, D.J.B., Wang, L., Jones, A., Wojciechowicz, M.L., Torre, D., Jagodnik, K.M., Jenkins, S.L., McQuilton, P., Flamholz, Z., Silverstein, M.C., et al. (2019). FAIRshake: Toolkit to Evaluate the FAIRness of Research Digital Resources. Cell Syst 9, 417–421.

70. Wilkinson, M.D., Sansone, S.-A., Schultes, E., Doorn, P., Bonino da Silva Santos, L.O., and Dumontier, M. (2018). A design framework and exemplar metrics for FAIRness. Scientific Data 5, 180118.

71. Wilkinson, M.D., Dumontier, M., Aalbersberg, I.J.J., Appleton, G., Axton, M., Baak, A., Blomberg, N., Boiten, J.-W., da Silva Santos, L.B., Bourne, P.E., et al. (2016). The FAIR Guiding Principles for scientific data management and stewardship. Sci Data 3, 160018.

72. Babb, L., Bult, C., Carey, V.J., Carroll, R.J., Hitz, B.C., Mungall, C.J., Rehm, H.L., Schatz, M.C., Wagner, A., and NHGRI Resource Workshop Community (2025). Improving the FAIRness and sustainability of the NHGRI resources ecosystem. arXiv [q-bio.GN]. 10.48550/arXiv.2508.13498.

73. Murphy, F., Bar-Sinai, M., and Martone, M.E. (2021). A tool for assessing alignment of biomedical data repositories with open, FAIR, citation and trustworthy principles. PLoS One 16, e0253538.

74. [No title] https://commonfund.nih.gov/sites/g/files/mnhszr341/files/Complement-ARIE-Landscape-An alysis-29-Feb-2024-508.pdf.

75. Keller, M.S., Gold, I., McCallum, C., Manz, T., Kharchenko, P.V., and Gehlenborg, N. (2025). Vitessce: integrative visualization of multimodal and spatially resolved single-cell data. Nat Methods 22, 63–67.

76. Bueckle, A., Qing, C., Luley, S., Kumar, Y., Pandey, N., and Börner, K. (2023). The HRA Organ Gallery affords immersive superpowers for building and exploring the Human Reference Atlas with virtual reality. Front Bioinform 3, 1162723.

77. Catalano, G., Delugas, E., Vignetti, S., Caputo, A., Martins Grapengiesser, I., Sousoni, D., Balsyte, E., and Pappas, D. UniProt case study and factsheet. 10.5281/zenodo.15732022.

78. Beagrie, N., and Houghton, J. Data-driven discovery: The value and impact of EMBL-EBI managed data resources. https://www.embl.org/documents/wp-content/uploads/2021/10/EMBL-EBI-impact-report-2021.pdf.

79. Kagda, M.S., Lam, B., Litton, C., Small, C., Sloan, C.A., Spragins, E., Tanaka, F., Whaling, I., Gabdank, I., Youngworth, I., et al. (2025). Data navigation on the ENCODE portal. Nat Commun 16, 9592.

80. Regev, A., Teichmann, S.A., Lander, E.S., Amit, I., Benoist, C., Birney, E., Bodenmiller, B., Campbell, P., Carninci, P., Clatworthy, M., et al. (2017). The Human Cell Atlas. Elife 6. 10.7554/eLife.27041.

81. Putman, T.E., Schaper, K., Matentzoglu, N., Rubinetti, V.P., Alquaddoomi, F.S., Cox, C., Caufield, J.H., Elsarboukh, G., Gehrke, S., Hegde, H., et al. (2024). The Monarch Initiative in 2024: an analytic platform integrating phenotypes, genes and diseases across species. Nucleic Acids Res 52, D938–D949.

82. International Cancer Genome Consortium, Hudson, T.J., Anderson, W., Artez, A., Barker, A.D., Bell, C., Bernabé, R.R., Bhan, M.K., Calvo, F., Eerola, I., et al. (2010). International network of cancer genome projects. Nature 464, 993–998.

83. All of Us Research Program Investigators, Denny, J.C., Rutter, J.L., Goldstein, D.B., Philippakis, A., Smoller, J.W., Jenkins, G., and Dishman, E. (2019). The “All of Us” Research Program. N Engl J Med 381, 668–676.

84. Sudlow, C., Gallacher, J., Allen, N., Beral, V., Burton, P., Danesh, J., Downey, P., Elliott, P., Green, J., Landray, M., et al. (2015). UK biobank: an open access resource for identifying the causes of a wide range of complex diseases of middle and old age. PLoS Med 12, e1001779.

85. Herr, B.W., Hardi, J., Quardokus, E.M., Bueckle, A., Chen, L., Wang, F., Caron, A.R., Osumi-Sutherland, D., Musen, M.A., and Börner, K. (2023). Specimen, biological structure, and spatial ontologies in support of a Human Reference Atlas. Scientific Data 10, 171.

86. Bandrowski, A., Brinkman, R., Brochhausen, M., Brush, M.H., Bug, B., Chibucos, M. C., Clancy, K., Courtot, M., Derom, D., Dumontier, M., et al. (2016). The Ontology for Biomedical Investigations. PLoS One 11, e0154556.

87. Diehl, A.D., Meehan, T.F., Bradford, Y.M., Brush, M.H., Dahdul, W.M., Dougall, D.S., He, Y., Osumi-Sutherland, D., Ruttenberg, A., Sarntivijai, S., et al. (2016). The Cell Ontology 2016: enhanced content, modularization, and ontology interoperability. J Biomed Semantics 7, 44.

88. Sarntivijai, S., Lin, Y., Xiang, Z., Meehan, T.F., Diehl, A.D., Vempati, U.D., Schürer, S.C., Pang, C., Malone, J., Parkinson, H., et al. (2014). CLO: The cell line ontology. J Biomed Semantics 5, 37.

89. Kim, S., Chen, J., Cheng, T., Gindulyte, A., He, J., He, S., Li, Q., Shoemaker, B.A., Thiessen, P.A., Yu, B., et al. (2025). PubChem 2025 update. Nucleic Acids Res 53, D1516–D1525.

90. Ison, J., Kalas, M., Jonassen, I., Bolser, D., Uludag, M., McWilliam, H., Malone, J., Lopez, R., Pettifer, S., and Rice, P. (2013). EDAM: an ontology of bioinformatics operations, types of data and identifiers, topics and formats. Bioinformatics 29, 1325–1332.

91. Schriml, L.M., Mitraka, E., Munro, J., Tauber, B., Schor, M., Nickle, L., Felix, V., Jeng, L., Bearer, C., Lichenstein, R., et al. (2019). Human Disease Ontology 2018 update: classification, content and workflow expansion. Nucleic Acids Res 47, D955–D962.

92. Ashburner, M., Ball, C.A., Blake, J.A., Botstein, D., Butler, H., Cherry, J.M., Davis, A.P., Dolinski, K., Dwight, S.S., Eppig, J.T., et al. (2000). Gene ontology: tool for the unification of biology. The Gene Ontology Consortium. Nat Genet 25, 25–29.

93. Gargano, M.A., Matentzoglu, N., Coleman, B., Addo-Lartey, E.B., Anagnostopoulos, A.V., Anderton, J., Avillach, P., Bagley, A.M., Bakštein, E., Balhoff, J.P., et al. (2024). The Human Phenotype Ontology in 2024: phenotypes around the world. Nucleic Acids Res 52, D1333–D1346.

94. Smith, C.L., and Eppig, J.T. (2009). The mammalian phenotype ontology: enabling robust annotation and comparative analysis. Wiley Interdiscip Rev Syst Biol Med 1, 390–399.

95. Schoch, C.L., Ciufo, S., Domrachev, M., Hotton, C.L., Kannan, S., Khovanskaya, R., Leipe, D., Mcveigh, R., O’Neill, K., Robbertse, B., et al. (2020). NCBI Taxonomy: a comprehensive update on curation, resources and tools. Database (Oxford) 2020. 10.1093/database/baaa062.

